# Rapid host response to an infection with Coronavirus. Study of transcriptional responses with Porcine Epidemic Diarrhea Virus

**DOI:** 10.1101/2020.07.28.224576

**Authors:** Wei Hou, Fei Liu, Wim H.M. van der Poel, Marcel M. Hulst

## Abstract

The transcriptional response in Vero cells (ATCC^®^ CCL-81) infected with the coronavirus Porcine Epidemic Diarrhea Virus (PEDV) was measured by RNAseq analysis 4 and 6 hours after infection. Differential expressed genes (DEGs) in PEDV infected cells were compared to DEGs responding in Vero cells infected with Mammalian Orthoreovirus (MRV). Functional analysis of MRV and PEDV DEGs showed that MRV increased the expression level of several cytokines and chemokines (e.g. IL6, CXCL10, IL1A, CXCL8 [alias IL8]) and antiviral genes (e.g. IFI44, IFIT1, MX1, OASL), whereas for PEDV no enhanced expression was observed for these “hallmark” antiviral and immune effector genes. Pathway and Gene Ontology “enrichment analysis” revealed that PEDV infection did not stimulate expression of genes able to activate an acquired immune response, whereas MRV did so within 6h. Instead, PEDV down-regulated the expression of a set of zinc finger proteins with putative antiviral activity and enhanced the expression of the transmembrane serine protease gene TMPRSS13 (alias MSPL) to support its own infection by virus-cell membrane fusion (Shi et al, 2017, Viruses, 9(5):114). PEDV also down-regulated expression of Ectodysplasin A, a cytokine of the TNF-family able to activate the canonical NFKB-pathway responsible for transcription of inflammatory genes like IL1B, TNF, CXCL8 and PTGS2. The only 2 cytokine genes found up-regulated by PEDV were Cardiotrophin-1, an IL6-type cytokine with pleiotropic functions on different tissues and types of cells, and Endothelin 2, a neuroactive peptide with vasoconstrictive properties. Furthermore, by comprehensive datamining in biological and chemical databases and consulting related literature we identified sets of PEDV-response genes with potential to influence i) the metabolism of biogenic amines (e.g. histamine), ii) the formation of cilia and “synaptic clefts” between cells, iii) epithelial mucus production, iv) platelets activation, and v) physiological processes in the body regulated by androgenic hormones (like blood pressure, salt/water balance and energy homeostasis). The information in this study describing a “very early” response of epithelial cells to an infection with a coronavirus may provide pharmacologists, immunological and medical specialists additional insights in the underlying mechanisms of coronavirus associated severe clinical symptoms including those induced by SARS-CoV-2. This may help them to fine-tune therapeutic treatments and apply specific approved drugs to treat COVID-19 patients.

## Introduction

The lack of knowledge for treating hospitalized SARS-CoV-2 infected patients is one of the pressing problems of the current COVID-19 pandemic. The SARS-CoV-2 virus shows a close genetic similarity to the in April 2003 identified SARS virus (SARS-CoV-1) and to other SARS-related coronaviruses isolated from humans and bats.

SARS-CoV-2 induces clinical respiratory symptoms familiar to the 2003 virus, mostly in persons with underlying diseases like COPD, heart failure, diabetes and obesity (1: Wu *et al.* 2020). Despite the 2003 SARS-CoV-1 virus has been extensively studied in the last two decades, there are no vaccines available yet, neither there are effective prophylactic and therapeutic treatment regimens with drugs that work equally well for each individual patient with SARS-induced respiratory problems. Such treatments might prevent development of severe disease patterns like “acute respiratory distress syndrome” (ARDS) and other, often fatal complications, and may decrease the case-fatality rate of SARS-CoV-2 infections.

In our lab we study the alpha-coronavirus PEDV. PEDV was first detected in pig herds in 1977 in Europe (2: Pensaert and de Bouck 1978). However, this virus reemerged in the spring of 2013 in North America causing a massive outbreak among pig herds, resulting in the death of about 30% of the suckling piglets due to severe diarrhea and dehydration (3: Huang *et al.* 2013, 4: Jung K. and Saif LJ. 2015). In the following years, PEDV was detected in many countries around the world. Intensified surveillance programs for PEDV frequently detected Mammalian Orthoreovirus serotype 3 (MRV3) in the fecal samples of diarrheic piglets tested positive for PEDV. Firstly in the USA during the PEDV epidemic in 2013 (5: Nayaranappa *et al.* 2015), and in the following years also in fecal samples of PEDV infected piglets in Europe (6: Lelli *et al.* 2016, 7: Hulst *et al.* 2017). MRV is able to infect many types of mammals and is widely distributed around the world, with bats as the main animal reservoir.

Infections of MRV have been reported in humans and can cause mild (often asymptotic) to severe respiratory symptoms, gastroenteritis, or encephalitis in young children (8: Steyer *et al.* 2013, 9: Tyler 1998, 10: Chua *et al.* 2008). Although several studies concluded that these clinical symptoms were caused by MRV itself, in concordance with the co-existence of MRV3 in PEDV infected piglets, also other MRV serotypes were isolated from hospitalized patients with airway problems diagnosed positive for SARS-CoV-1 (11: Cheng *et al.* 2009, 12: Duan *et al.* 2003, 13: Zuo *et al.* 2003). Recently, a cross-family recombinant coronavirus was isolated in China from bat faeces in which an RNA sequence originating from the S1 segment of MRV was inserted in the coronavirus genome between the N and Ns7a genes, indicating that both viruses were replicating simultaneously in a single cell in bats (14: Huang *et al.* 2016). A prevalence study showed that this cross-family recombinant coronavirus circulated in an isolated bat colony in a cave in China (15: Obameso *et al.* 2017).

This cooccurrence of MRV with coronaviruses raised the questions whether a synergistic effect between both viruses exists and if such coexistence plays a role in viral pathogenesis. Therefore we studied the host response in cultured cells early (4 and 6 hours) after PEDV and MRV infection using RNAseq. Our original goal was to identify early factors and processes induced by PEDV or MRV that could stimulate or influence the replication and pathogenesis of the other virus.

The host, tissue and cell tropism of PEDV differs from SARS-CoV-1 and −2. However, the genomic organization, replication strategy and function of a part of the viral non-structural proteins share common features among all coronaviruses (16: Brian and Baric 2005). This applies particularly for interactions in infected cells of non-structural coronavirus proteins with specific host proteins. Host proteins that are recruited or silenced to support virus replication, assembly and release. In our experiment we used Vero cells (Cercopithecus aethiops epithelial kidney cell line; ATCC® CCL-81) because these cells support efficient infection and replication of both MRV and PEDV. Vero cells are susceptible for many coronaviruses, including SARS-CoV-1 and −2 (17: Chu *et al.* 2020). They originate from epithelial tissue, in part resembling nasal and bronchial epithelium cells, the prime target cells infected by SARS-CoV-2 in the airways of humans. Recent research showed that SARS-CoV-2 is also able to replicate in epithelial cells of human small intestinal organoids (18: Lamers *et al.* 2020). A disadvantage of Vero cells is a deletion in the type I interferon (IFN) gene cluster on chromosome 12 (19: Osada *et al.* 2014). Therefore, these cells lack expression of type I IFNs important for activation of antiviral defense mechanisms. However, research has shown that Vero cells by-pass this IFN-activation route and could mount an antiviral response mediated by interferon regulatory factor 3 (20: Chew *et al.* 2009).

Single infections with PEDV or MRV3 alone and simultaneous (double) infections of Vero cells with both viruses were performed using a maximum multiplicity of infection (MOI) to achieve a synchronized infection of all cells. By RNAseq measured expression levels of mRNA transcripts/genes in infected cells were compared to RNAseq profiles measured from similar treated mock-infected cells harvested at the same time point after infection. The detected sets of differential expressed genes (DEGs) for PEDV and MRV were analyzed by gene set enrichment analysis (GSEA) using functional bioinformatic programs to retrieve biological processes (pathways and Gene Ontology terms [GO-term]) and associations with chemical compounds, including drugs. In addition, we searched the literature for functional information of the PEDV-DEGs to find possible associations with SARS-CoV-2 pathogenesis. Because of the COVID-2 pandemic we gave priority to publish the results of this functional bioinformatical analysis and datamining for the single infected Vero cells with PEDV separate from the results of the double infections with MRV3. In this report we focused on the “very early” host response of epithelial cells to an infection with the coronavirus PEDV and pay less attention to the role of specific viral proteins in this host response to PEDV. In part our results were in agreement with results of a previous RNAseq study comparing SARS-CoV-2 and Influenza host responses by RNAseq (21: Blanco-Melo *et al.* 2020). But we also found associations with biological processes, and pivotal genes/proteins acting in these processes, that had not been recognized before. This information may contribute to the search for novel or alternative preventive or therapeutic drugs and treatment protocols for this devastating COVID-19 disease.

## Results

### RNA transcription profiling

A time-dependent infection experiment was performed with cultured Vero cells. Details are described in supplementary file 1 (material and methods) and visually displayed in this file. Briefly, overnight cultured Vero cells grown in 2 cm^2^ wells were mock-infected, infected with MRV3 strain WBVR (7: Hulst *et al.* 2017) or PEDV strain CV777 (2: Pensaert and de Bouck 1978, 22: Rasmussen *et al.* 2018]) with a multiplicity of infection of ≥1 for 30 min at 4°C. For PEDV and corresponding mock-infected cells, 10 μg/ml of trypsin in serum-free medium was used to facilitate infection of Vero cells during the whole experiment. All virus and mock-infected timepoints were performed in quadruplicate. After incubation for 30 min at 4°C, virus was discarded and cells were washed twice and supplied with fresh culture medium. Cells were incubated for 0, 2, 4, 6, and 16h at 37°C and 5% CO_2_. After incubation for the indicated times, cells were placed on ice before total RNA was isolated from three of the quadruplicate wells. The replication of both viruses in Vero cells was monitored using virus-specific RT-qPCR tests (Fig.1: methods and primers used for PCR are provided in supplementary file 1). In addition, cells in one of the quadruplicate wells incubated for 16h were fixated and stained with antibodies directed against the S2 spike protein of PEDV and the S1 attachment protein (α1) of MRV3. A decrease in CT-values for PEDV was not observed before 6 h post inoculation (6 h.p.i), indicating that replication in PEDV infected cells started later than was observed for MRV (at 4 h.p.i). Staining of the cells after 16h indicated that nearly all Vero cells were infected with MRV3 and more than 50% with PEDV. Also more than 50% of the cells in 16h-wells appeared as fused cells (syncytia), confirming that more than 50% of the cells were infected with PEDV. Quality control of the total RNA isolated from infected cells using an Agilent Bioanalyzer showed that RNAs isolated from PEDV infected wells at 16 h.p.i. were partially degraded (RIN values below 9), making them unsuitable for RNAseq analysis. Therefore, only 0, 4 and 6h timepoints were analyzed using RNAseq.

**Fig. 1.**
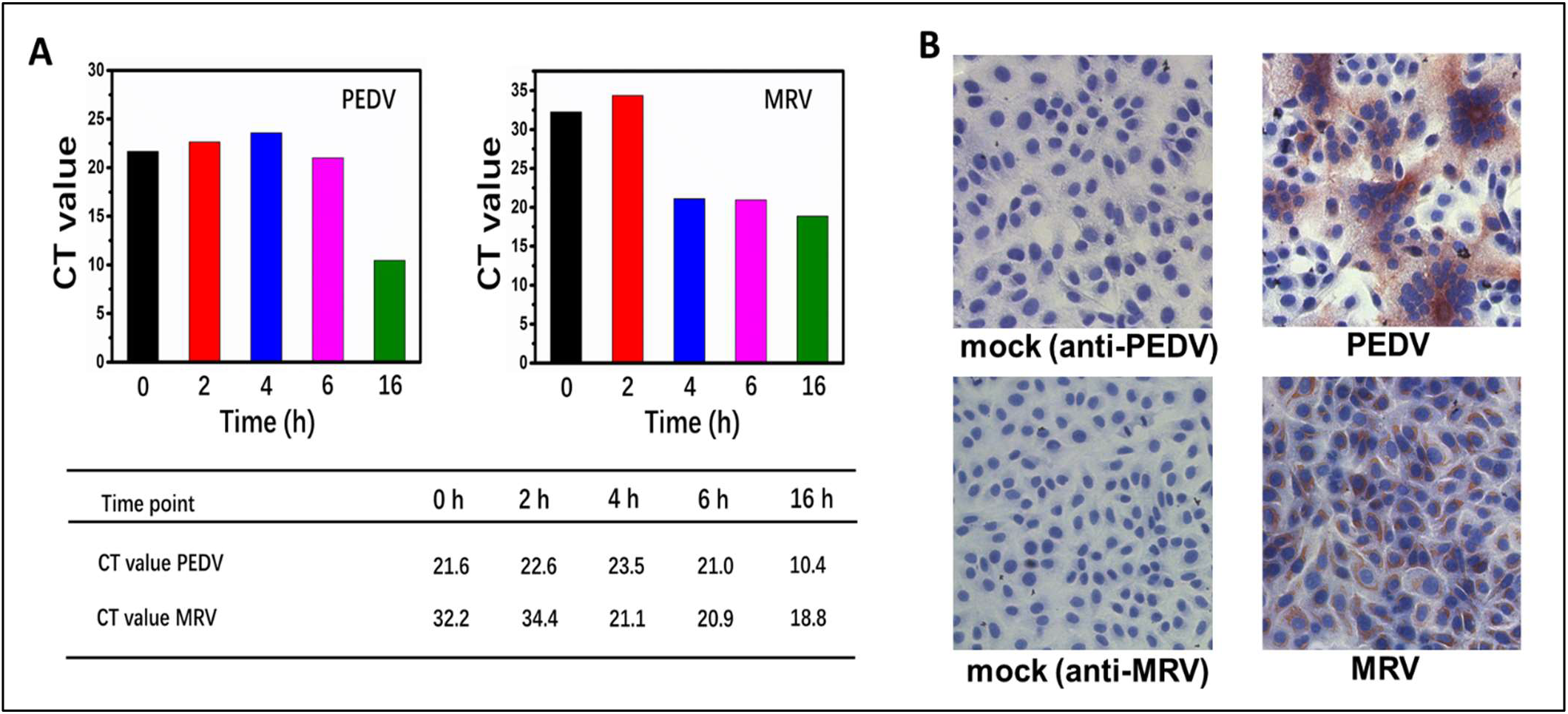
(A) Detection of PEDV and MRV replication in Vero cells at 0, 2, 4, 6 and 16h post infection by RT-qPCR. CT values are provided in the table beneath the graphs. (B) Immuno-Peroxidase Monolayer Assay (IPMA) of virus infected cells and mock infected cells (control) 16h after infection. PEDV and mock infected cells were stained with a monoclonal antibody directed against the S2 spike protein. MRV and mock infected cells were stained with a polyclonal rabbit serum raised against a peptide sequence of the S1-attachment protein of MRV serotype 3. Nuclei were stained blue with the Hoechst, 4’,6-diamidino-2-phenylindole dye.

Equal amounts of total RNA isolated from triplicate wells were pooled and subjected to RNAseq analysis by GenomeScan B.V.(Leiden, The Netherlands) using Next Generation Sequencing (NGS) (see supplementary file 2a for details). Mapping of NGS reads to the Cercopithecus aethiops reference genome and preparation of datafiles with calculated Fold Change (FC) of expression levels of mapped mRNAs, were performed for each comparison at 0, 4, and 6h by GenomeScan (see supplementary file 2b). From these datafiles we extracted lists of DEGs with a FC>2 and p-value of <0.05. After accessing the NCBI, Panther or KEGG databases for human orthologs, not annotated Cercopithecus aethiops DEGs were annotated with an HUGO official gene symbols (http://www.genenames.org). In supplementary file 3 sheet “PEDV-MRV DEGs FC>2”, lists of all annotated PEDV and MRV DEGs are presented with their FC. In a separate sheet “PEDV-DEGs functional info” all 266 individual DEGs regulated by PEDV at 4 and 6 h.p.i. are presented with their FC, information about their function and the types of human cells in which expression of the gene is relatively high compared to other human cells (retrieved from the “Primary Cell Atlas” dataset of BioGPS: http://biogps.org/). Note that all tables in these Excel sheets of supplementary file 3 are sortable using the headers. In all results paragraphs beneath information about the biological function of DEGs was retrieved by consulting the “GeneCards” (Weizmann Institute of Science: https://www.genecards.org/) and NCBI Gene reports (Entrez Gene: https://www.ncbi.nlm.nih.gov/gene/), and literature linked to these reports (for references about these biological functions of genes/proteins we refer to publications cited in these reports: “GeneCards” weblinks are provided in supplementary file 3).

Sets of PEDV and MRV DEGs were analyzed using the GSEA program GeneAnalytics (LifeMap Sciences, Inc.) and pathways (for MRV and PEDV), GO-terms (not for MRV), and associations with compounds/drugs (not for MRV) with a high or medium score (p-value <0.05) were retrieved and listed in 3 separate sheets in supplementary file 3 (sheets “MRV-PEDV pathways”, “PEDV G0-terms” and “PEDV Compounds”). Similar and related pathways retrieved for both PEDV and MRV, and remarkable PEDV pathways, GO-terms and compound associations are summarized in Table 1. For PEDV all DEGs within these pathways are provided with their regulation, up (green) or down (red). For MRV only DEGs in common with PEDV-DEGs were listed in Table 1 (see sheet “MRV-PEDV pathways” in supplementary file 3 for all MRV-DEGs acting in these pathways). Subsets of PEDV-DEGs were selected matching the terms “chemokines-cytokines”, “antiviral” , and terms related to the pathogenesis of COVID-2 (explained below) using the genotyping program VarElect (LifeMap Sciences, Inc.) and displayed in supplementary file 3 in separate sheets: “chemokines-cytokines”, “(anti)-viral”, etc. Based on these selections we prepared a set of PEDV KEY-DEGs consisting of genes regulated with a FC of >10 (up) or <−10 (down) or playing an important role in biological processes induced by PEDV and related to COVID-19 pathology. In beneath results sections we tried to give as much as possible meaningful information about the function of KEY-DEGs for which we found an association with SARS-CoV-2 infections. We emphasize that further dedicated experimental and in-silico research is necessary to confirm the involvement of the proteins encoded by these genes for pathogenesis of this viral disease.

**Table 1.** Enriched pathways, GO-terms and compound associations of PEDV-DEGs

| (A) PEDV and related MRV pathways (h.p.i.) | *PEDV and similar or related MRV enriched Pathways | Matched Genes (Symbols)<sup>5</sup> |
| --- | --- | --- |
| PEDV-4h/MRV-4 and 6h | Transcription_Role of VDR in Regulation of Genes Involved in Osteoporosis (Vitamin-D metabolism) | CYP24A1, COL1A1 (MRV-4h 5 genes, MRV-6h 6 genes, no common genes to PEDV) |
| PEDV-4h/MRV-6h | Tacrolimus/Cyclosporine Pathway (drug-induced regulation of NFAT transcription factor activity in T-cells by immunophilin [PPIA]; down-regulation of inflammatory cytokine expression) | PPIA, NRL (MRV-6h 8 genes; no common genes to PEDV) |
| PEDV-6h/MRV-4h | Signaling By Rho GTPases (ARHGAP42 and its paralogs OPHN1; inhibition of RhoA activity in regulating vascular tone and blood pressure and down-regulated expression of autoantigens) | HIST2H2BE, H2AFZ, ARHGAP42, HIST4H4, NOX3, SFN, OPHN1 (MRV-4h 14 genes; OPHN1 in common) |
| PEDV-4h/MRV-6h | Innate Immune System (transcription of genes for vesicle formation) | EIF4E3, AHSB, HLA-H, HSP90B1, HIST1H3A, AOC1, GAB2, EDARADD, ATP8B4, MAOA, BLNK, PPIA, SPRED3, RORC, CD53, RASA4, PI3, CXCR1, RASGEF1A, MUC5AC, EDA (MRV-6h 69 genes; RORC, RASA4 in common with PEDV) |
| PEDV-6h/MRV-6h | Developmental Biology (transcription of genes involved in forming "cornified cell envelopes"; mediated by ephrin ligands) | LCE3B, HIST2H2BE, COL9A3, KRT25, H2AFZ, EPHA10, HSPA8, HIST4H4, SHC3, MEF2B, PIK3CB, RND1, SCN4B (MRV-6h 42 genes; PIK3CB in common with PEDV) |
| PEDV-4h/MRV-4 and 6h | Cytokine Signaling in Immune System (PEDV-RORC up-regulation/MRV-RORC down-regulation) (PEDV up-regulation of Endothelin 2 [EDN2] at 4h and Cardiotrophin 1 [CTF1] at 6h) | EIF4E3, HLA-H, HSP90B1, HIST1H3A, GAB2, EDARADD, MAOA, BLNK, SPRED3, RORC, RASA4, RASGEF1A, EDA (MRV-4h 21 genes, MRV-6h 41 genes; RORC, RASA4, HIST1H3A in common with PEDV) |
| PEDV-4h/MRV-4 and 6h | NF-kappa B Signaling Pathway (down-regulated EDA-EDARADD signaling: reduced NF-kappaB induced IL1B, IL8, TNF, PTGS2, NFKBIA, TNFAIP3 transcription) | EDARADD, BLNK, EDA (MRV-6h 22 genes; no common genes to PEDV) |
| PEDV-6h/MRV-6h | PI3K / Akt Signaling (ERAS activates the phosphatidylinositol 3-kinase signal transduction pathway) | BORCS6, ERAS, PIK3CB (MRV-6h 28 genes; PIK3CB in common with PEDV) |
| PEDV-4h/MRV-4 and 6h | TNFs Bind Their Physiological Receptors (down-regulated EDA-EDARADD signaling: reduced activation of NF-kappaB induced IL1B, IL8, TNF, PTGS2, NFKBIA, TNFAIP3 transcription) | EDARADD, EDA (MRV-6h 14 genes; no common genes to PEDV) |
| PEDV-4h | Cell Adhesion_ECM Remodeling (CXCR1; high affinity receptor for CXCL8=IL8, involved in neutrophil activation) | COL1A1, COL3A2, CXCR1 |
| PEDV-4h | Diseases of Glycosylation (down-regulation of forming of an epithelial mucus layer) | SEMA5A, MUC5AC, SSPO |
| PEDV-4h | Histidine Metabolism (biogenic amine metabolism; histamine-[derivates]) | AOC1, MAOA |
| PEDV-4h | Inflammatory Response Pathway (COL1A1 and COL3A1; inflammation fibrotic response) | COL1A1, COL3A1 |
| PEDV-4h | Influenza Viral RNA Transcription and Replication (RPS genes; ribosomal proteins involved in translation) | RPS2, RPL36, RPL35A, RPS15 |
| PEDV-4h | Phenylalanine Metabolism (biogenic amine metabolism) | AOC1, MAOA, IL4I1 |
| PEDV-4h | Tryptophan Metabolism (biogenic amine metabolism) | AOC1, MAOA, IL4I1 |
| PEDV-6h | C-MYB Transcription Factor Network (PPIA forms a complex with H2AFZ in response to DNA damage) | CSF1R, H2AFZ, HSPA8 |
| PEDV-6h | Eicosanoid Synthesis (conversion of prostaglandin endoperoxide H2 [PGH2] to prostaglandin E2 [PGE2]; involved in pathogenesis of collagen-induced arthritis and acute pain during inflammatory) | GGT1, PTGES |
| PEDV-6h | Histidine Metabolism (biogenic amine metabolism; histamine-[derivates]) | AMDHD1, HNMT |
| PEDV-6h | Histidine, Lysine, Phenylalanine, Tyrosine, Proline and Tryptophan Catabolism (biogenic amine metabolism) | AMDHD1, HNMT, PRODH |
| PEDV-6h | Cell Surface Interactions at The Vascular Wall (PPIA forms a complex with H2AFZ in response to DNA damage) | JCHAIN, PPIA, PIK3CB, PSG7, SLC7A9 |
| (B) PEDV h.p.i. | *PEDV-GO-terms | Matched Genes (Symbols)<sup>5</sup> |
| 4 | Translational Initiation (EIF4E3; Binds the 7-methylguanosine-mRNA cap in early step of translation and may act as an inhibitor of EIF4E1 activity) | EIF4E3, RPS2, RPL36, RPL35A, RPS15 |
| 4 | Viral Transcription (RPS genes; ribosomal proteins involved in translation) | RPS2, RPL36, RPL35A, RPS15 |
| 4 | Platelet Activation (F2RL2 down-regulation; reduced thrombotic and/or inflammatory response) | COL1A1, COL3A1, F2RL2, GNG2 |
| 4 | Primary Amine Oxidase Activity (biogenic amine metabolism; histamine-[derivates]) | AOC1, MAOA |
| 4 | Platelet-derived Growth Factor Binding (COL1A1 and COL3A1; inflammation fibrotic response) | COL1A1, COL3A1 |
| 4 | Non-motile Cilium (ANO2 down-regulation; reduction olfactory sensory amplification of odorant receptors present on non-motile cilia of dendritic knobs) | MCHR1, ANO2, (added DEG; OR2A14) |
| 6 | Axon Guidance (FEZ1; Fasciculation and/or normal axonal bundling and elongation within axon bundles) | FEZ1, CSF1R, EPHA10, FLRT2, SHC3, PIK3CB, OPHN1 |
| 6 | Cilium (CDK20 and MAK, inhibitors of cilia formation) | GPR19, DNAH10, CCDC114, CDK20, MAK, TULP2 |
| 6 | Axoneme (MAK; regulates intraflagellar transport (IFT) speed and negatively regulates cilia length) | DNAH10, CCDC114, MAK |
| 6 | Histidine Catabolic Process (biogenic amine metabolism; histamine-[derivates]) | AMDHD1, HNMT |
| 6 | Secretory Granule Lumen (ATPase activity of HSPA8 contributes to disassembly of clathrin-coated vesicles in the cell) | EEF1A1, GHDC, HSPA8, PPIA |
| (C) PEDV h.p.i. | *PEDV compound association | Matched Genes (Symbols)<sup>5</sup> |
| 4 | Ammonia (GCSH, AOC1, MAOA, IL4I1, enzymatic by-product) | GCSH, AOC1, MAOA, IL4I1 |
| 4 | Bepridil (calcium-blocking drug/calmodulin antagonist with antihypertensive and selective anti-arrhythmia activities) | KCNQ4, CACNA1H |
| 4 | Hydrogen Peroxide (AOC1, MAOA, IL4I1, enzymatic by-product) | AOC1, MAOA, IL4I1 |
| 4 | Indoleacetaldehyde (enzymatic product catalysis by AOC1, MAOA) | AOC1, MAOA |
| 4 | NPPA (prohormone natriuretic Peptide A (ANP); inhibition cardiac hypertrophy in heart failure as well as fibrosis) | COL1A1, COL3A1 |
| 4 | NPPB (cardiac hormone; regulates natriuresis, diuresis, vasorelaxation, and has a key role in cardiovascular homeostasis) | COL1A1, COL3A1 |
| 4 | Phenylacetaldehyde (enzymatic product catalysis by AOC1, MAOA) | AOC1, MAOA |
| 4 | Tele-methylhistamine (histamine methyltransferase antagonist) | AOC1, MAOA |
| 6 | 4-methylumbelliferone (choleoretic drug; promoting bile secretion by the liver) | CES1, UGT1A9 |
| 6 | Dabigatran Etexilate (Thrombin inhibitor drug, blocking thrombogenic activity) | CES1, UGT1A9 |
| 6 | Dihydroartemisinin (malaria drug) | EEF1A1, HSPA8, PPIA, UGT1A9, NPM1 |
| 6 | Excitatory Amino Acid Antagonists (synaptic transmission) | ERAS, SFN |
| 6 | Tacrine (Anticholinesterase drug) | HNMT, CES1 |

*\*PEDV enriched pathways (A), GO-terms (B), and associations with compounds and drugs (C) with a high and medium score and with at least 2 matching genes were retrieved from GeneAnalytics.*
\*Official Gene-symbols (HUGO abbreviations) are listed for DEGs. Down-regulated DEGs were colored red and up-regulated DEGs were colored green. In section A the number of DEGs regulated by MRV in a pathway and the common DEGs are provided between brackets. DEGs regulated by both PEDV and MRV are underlined.

### Regulation of immune and antiviral genes

Compared to MRV, only a few genes involved in “cytokine/chemokine signaling” were regulated at 4 and 6h by PEDV. In Fig. 2A the regulation of cytokines/chemokines in PEDV and MRV infected Vero cells are displayed. This indicated that MRV increased the transcription of a broad set of cytokines/chemokines, including interferon-mediated cytokines like CXCL10 and CXCL8 (alias IL8), already at 4 h.p.i., whereas PEDV did not, even not when replication of PEDV RNA was detected by RT-qPCR at 6 h.p.i.. For MRV, this cytokine/chemokine response at 4 h.p.i. was followed by high up-regulation of “hallmark” antiviral genes at 6 h.p.i. (see Fig. 2B: e.g. interferon-induced genes [IFI] and OASL) and chemokines that attract T cells, monocytes, granulocytes, including basophils (e.g. CXCL8, CXCL11 and CCL2). PEDV infection up-regulated only a few genes coding for proteins with cytokine activity (CTF1 and EDN2), and also did not elevated gene expression of these “hallmark” antiviral genes. In contrast, PEDV down-regulated expression of 6 genes (out of 9 in total) coding for Zinc Finger Proteins (out of 9 in total), all 6 with an antiviral activity towards Herpes simplex virus 1 (Fig 2C) and of the gene coding for Heat Shock Protein Family A (Hsp70) Member 8 (HSPA8; 12-fold at 6h). Binding of HSPA8 to the measles virus nucleocapsid protein (N) negatively affected virus replication. An indication that PEDV provoked a transcriptional response in Vero cells was the elevated expression of the “Transmembrane Serine Protease 13” gene (TMPRSS13, alias MSPL). It was recently reported that expression of this protease was stimulated in response to PEDV infection (23: Shi *et al.* 2017). TMPRSS13 cleaves the PEDV spike protein resulting in virus-cell membrane fusion and subsequent entry of virus particles into cells. A similar mechanism of virus entry into cells is used by SARS coronaviruses and influenza A viruses (24: Zmora *et al.* 2014, 25: Matsuyama *et al.* 2010). The only 2 genes with cytokine activity found up-regulated were Endothelin 2 (EDN2, a neuroactive secreted peptide with vasoconstrictive properties) and Cardiotrophin 1 (CTF1). The latter codes for a protein that binds to the IL6-signal transducer (IL6ST, alias GP130) receptor and leukemia inhibitory factor receptor (LIFR). Both these receptors initiate phosphorylation of STAT3, a transcriptional activator of interferon (IFN) genes and IL6. In a recent study EDN2 gene expression was also up-regulated in alveolar basal epithelial cell line A549 in response to SARS-CoV-2 infection (21: Blanco-Melo *et al.* 2020, see also below). CTF1 induces pleiotropic physiological effects in the body, among them, induction of cardiomyocyte hypertrophy, regulation of energy metabolism, reduction of lipid accumulation and insulin sensitivity in skeletal myocytes, and regulation of inflammation (see also below). PEDV down-regulated the expression of the cytokine Ectodysplasin A (EDA), a cytokine of the TNF-family, and of the receptor to which EDA binds (EDAR Associated Death Domain: EDARADD). EDA-EDARADD signaling activates canonical-NFKB transcription of inflammatory genes like IL1B, TNF, CXCL8 and Prostaglandin-Endoperoxide Synthase 2. PTGS2 is responsible for synthesis of prostaglandin H2, a chemical immune modulator.

**Fig. 2.**
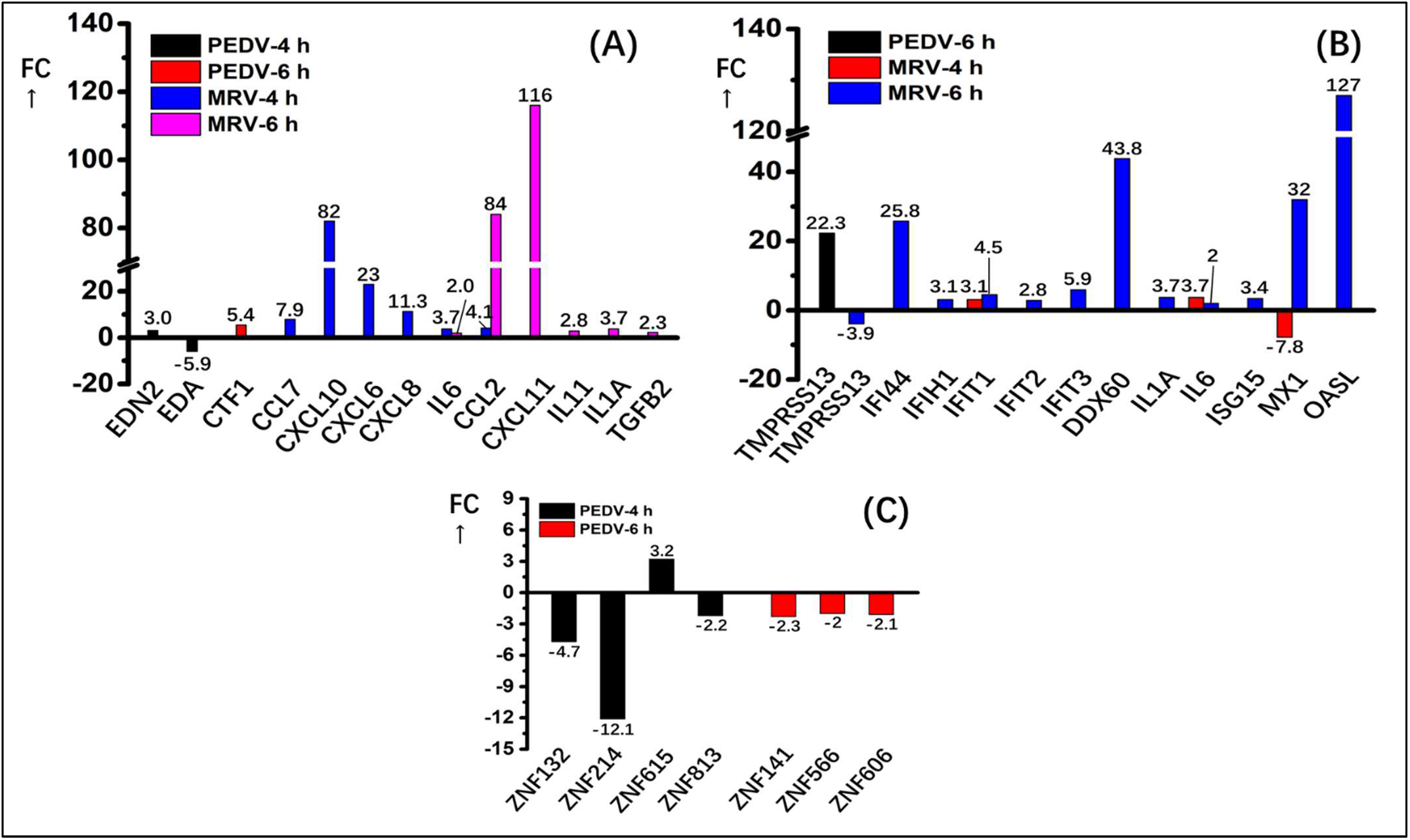
Fold Change (FC) of cytokine and chemokine gene expression (A), antiviral gene expression (B) and zinc finger genes with a putative antiviral activity (C; only shown for PEDV) measured in MRV and PEDV infected Vero cells at 4 and 6 h.p.i. The regulation of TMPRSS13 gene expression by MRV and PEDV was included in graph B. Note that negative values represent the FC of down-regulated genes.

Pathway analysis indicated that MRV enhanced the expression of a broad set of genes in Vero cells, within 6h after infection, involved in activation of acquired immune responses by T-cells. In contrary, only a few genes related to T-cell activation were regulated by PEDV after 6h (see sheet “MRV-PEDV pathways” in supplementary file 3). Therefore, we searched for PEDV-DEGs which regulation could contribute to dysregulation of T-cell responses. PEDV-DEGs with potential to do so are the receptors SLAMF8, CXCR1, and F2RL2 (thrombin receptor-like 2, alias PAR3), all down-regulated 5 to 10-fold at 4 h.p.i., and the upregulated genes Colony Stimulating Factor 1 Receptor (CSF1R; 7-fold at 6h), B cell Linker (BLNK; up-regulated 10-fold at 4h) and nuclear receptor/transcription factor RORC (up-regulated 10-fold at 4h). Transcription factor RORC was down-regulated by MRV at 6h. RORC mediates the expression of IL17 in helper Th17 cells (Th17). However, it was reported that specific isoforms of RORC exist that can suppress expression of IL17, IL2, and FASLG in T cells (26: Rauen *et al.* 2012). Binding of measles virus hemagglutinin to SLAMF surface receptors inhibits IL12 expression by dendritic cells (DCs). The S1 spike glycoprotein of specific PEDV strains and the majority of beta-coronaviruses (SARS-CoV-2 included) also induce hemagglutination of human erythrocytes (27: de Groot *et al.* 2006). CXCR1 is a high-affinity receptor for CXCL8 (IL8) on the surface of mast and T cells. Binding of thrombin to F2RL2 reduces inflammation, activates platelets and increases vasodilation and permeability of the vascular wall (see also below in the section “platelets activation”). CSF1R is a receptor for the cytokine colony stimulating factor 1, a cytokine that regulates differentiation and function of macrophages, and in the CNS, the density and distribution of microglia cells. The BLNK gene codes for a cytosolic protein that passes on B-cell receptor signals in the signaling cascade that activates B-cell development and function. Gene expression of genes coding for essential components of this B-cell signaling, like “spleen associated tyrosine kinase” (SYK) and “LYN proto-oncogene”(LYN) were not regulated by PEDV, nor by MRV.

### Genes involved in amino acid, protein translation and metabolism of immuno-active compounds

PEDV induced a 2-fold up-regulation of the gene coding for Eukaryotic Translation Initiation Factor 4E Family Member 3 (EIF4E3) at 4 h.p.i. and strong down-regulation of Eukaryotic Translation Elongation Factor 1 Alpha 1 (EEF1A1, 20-fold) at 6 h.p.i.. EIF4E3 interacts with 5-prime cap structures of mRNA’s and recruits these capped RNA’s to the ribosomes. Coronavirus genomic and sub-genomic RNA’s are also capped at the 5-prime ends and methylated by the non-structural viral protein nsp14, which possesses guanine-N7-methyltransferase (N7-Mtase) activity (28: Chen *et al.* 2009). It was shown that additional 2’-O methylation of viral caps by the coronavirus nsp16/nsp10 2’-O-Mtase counteracts the innate antiviral response of IFIT family members (Interferon Induced Protein With Tetratricopeptide Repeats) in host cells (29: Daffis *et al.* 2010). EEF1A1 directs aminoacyl-tRNA’s to the ribosomes during translation and is also a part of transcription factor-complex in T-helper 1 cells (Th1) that binds to the promotor of the IFNγ gene.

DEGs coding for enzymes involved in the metabolism of the non-essential amino acids histidine, phenylalanine, tryptophan and proline were found enriched in the PEDV dataset (see supplementary file 3, sheet PEDV-compounds). Remarkable were the DEGs coding for amine oxidases involved in the catabolism of the biogenic amines histamine, tryptamine and phenylethylamine, their derivates and related substrates/products of these enzymes (Fig. 3, AOC1, MAOA, IL4I1). None of these amino oxidase genes were regulated by MRV. Using the genotyping program VarElect, PEDV-DEGs with an association with these biogenic amines were retrieved (supplementary file 3, sheet “Biogenic amines”). Three enzymes clustered in the “Histidine metabolism” pathway (https://www.kegg.jp/kegg-bin/show_pathway?hsa00340+4128) with histamine and reaction products generated from this biogenic amine (Fig. 3). Also most association of PEDV-DEGs were found by VarElect for histamine. The gene coding for the amine oxidase “Interleukin 4 Induced 1” (IL4I1) was strongly down-regulated (30-fold) 4h after infection with PEDV. Besides catalysis of L-Phenylaniline into Phenylpyruvate (Fig.3), IL4I1 also fulfills an important role in signaling in “synaptic clefts” formed between antigen presenting cells (APC) and T cells (so-called “immune cleft”: see also below)(30: Boulland *et al.* 2007, 31: Molinier-Frenkel *et al.* 2019).

**Fig. 3.**
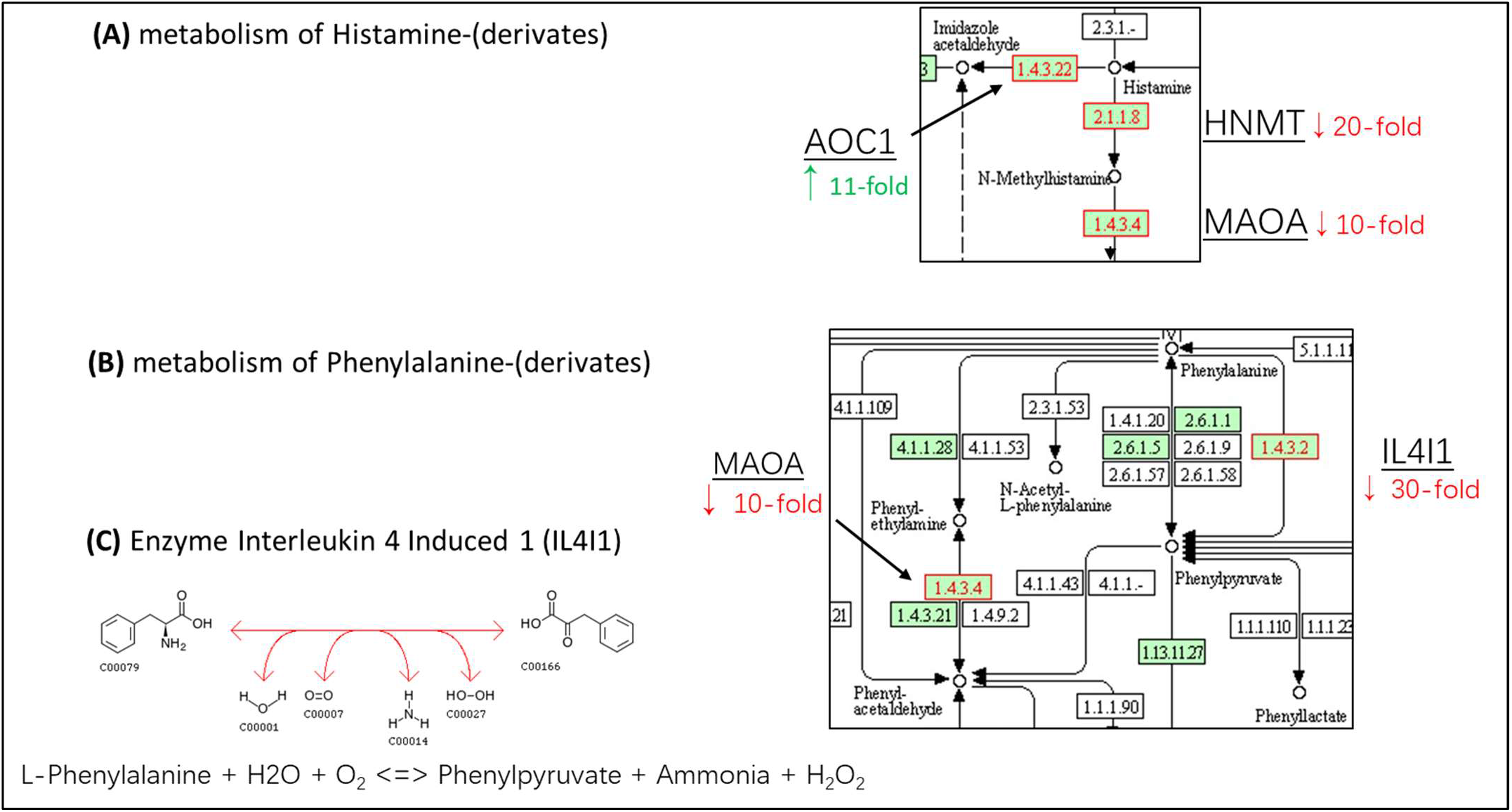
Regulation of genes involved in the metabolism of biogenic amines. Snapshots taken from the Kyoto Encyclopedia of Genes and Genomes pathway schemes “Histidine metabolism” (A) and “Phenylalanine metabolism” (B). Regulated genes are shown in red lined boxes. Up-regulation is indicated by a green arrows (↑) and down-regulation by red arrows (↓). (C) Substrates and reaction products of the reversible enzyme IL4I1.

Expression of the Prostaglandin-Endoperoxide Synthase 2 (PTGS2) gene was up-regulated by MRV at 6h p.i. PTGS2 synthesizes prostaglandin endoperoxide H2 (PGH2), an compound with a short half-life and the precursor of many biological active prostaglandins: e.g. Thromboxane-A2 (mediates activation of platelets), PGI2 and PGE2. In contrast, PEDV increased the expression of the gene coding for Prostaglandin E Synthase (PTGES) which converts PGH2 into PGE2. PGE2 is a direct vasodilator, but does not inhibit platelet aggregation. PGE2 also suppresses T cell receptor signaling. PEDV decreased expression of the Gamma-Glutamyltransferase 1 gene (GGT1) after 4h (2-fold), but increased expression of this gene 2 hours later to a 4-fold level compared to mock infected cells. GGT synthesizes Leukotriene D4 (LTD4) from LTC4. IgE-activated mast cells may secrete LTD4 and LTC4, together with histamine and platelets activating factor (PAF). This vesicle mediated secretion by mast cells (degranulation) results in stimulation of mucus production, and similar to histamine, increases the permeability and smooth muscle contraction of the vascular wall. In persons suffering from asthma this degranulation leads to an immediate allergic response (bronchospasm, airflow obstruction and forming of edema).

### Genes involved in “Cilia and Synaptic cleft” formation

GSEA detected “Axon Guidance” as the GO-term with the highest score for PEDV (see Table 1). In addition, PEDV-DEGs were enriched coding for proteins involved in calcium ion-dependent exocytosis from vesicles into the “synaptic clefts” between two cells (e.g. between axons and dendrites), and DEGs coding for proteins involved in formation of cilia. Cilia protruding from cells are found in many forms. They can have a static (structural) function, e.g. in forming of clefts between two cells (see Fig. 4), or a motile function. Motile cilia on the surface of ciliated cells lining up the epithelial layers in the nose, trachea and bronchia sweep out superfluous mucus containing dirt from the airways. PEDV DEGs matching the terms “Cilia” and “Synaptic cleft” retrieved form the genotyping program VarElect were further examined by consulting functional information in the NCBI Gene and GeneCards reports in order to evaluate their association with these processes (see supplementary file 3, sheet “Cilia and Synaptic cleft”). Based on this analysis we identified genes in the set of PEDV DEGs which can i) negatively regulate cell adhesion (RND1 and SEMA5A), ii) inhibit formation of cilia (kinases MAK1 and CDK20, highly up-regulated at 6h), and iii) regulate cytoskeleton rearrangements that facilitate axon growth and growth and stabilization of dendritic spines (F2RL2, FEZ1, OPHN1 and PIK3CB). Except for PIK3CB (Phosphatidylinositol-4,5-Bisphosphate 3-Kinase Catalytic Subunit Beta) and OPHN1 (Oligophrenin 1), all these matching PEDV-DEGS were not regulated after infection with MRV. The gene coding for “Fasciculation And Elongation Protein Zeta 1” (FEZ1) was highly up-regulated at 4 h.p.i. (21-fold). This protein associates with microtubules and kinesins to generate the force needed to transport intracellular organelles (e.g. synaptic vesicles or granules) to the cell membrane. The gene Oligophrenin 1 (OPHN1) was down-regulated at 4h and up-regulated at 6 h. OPHN1 is also involved in regulation of endocytosis of synaptic vesicles at presynaptic terminals.

**Figure 4.**
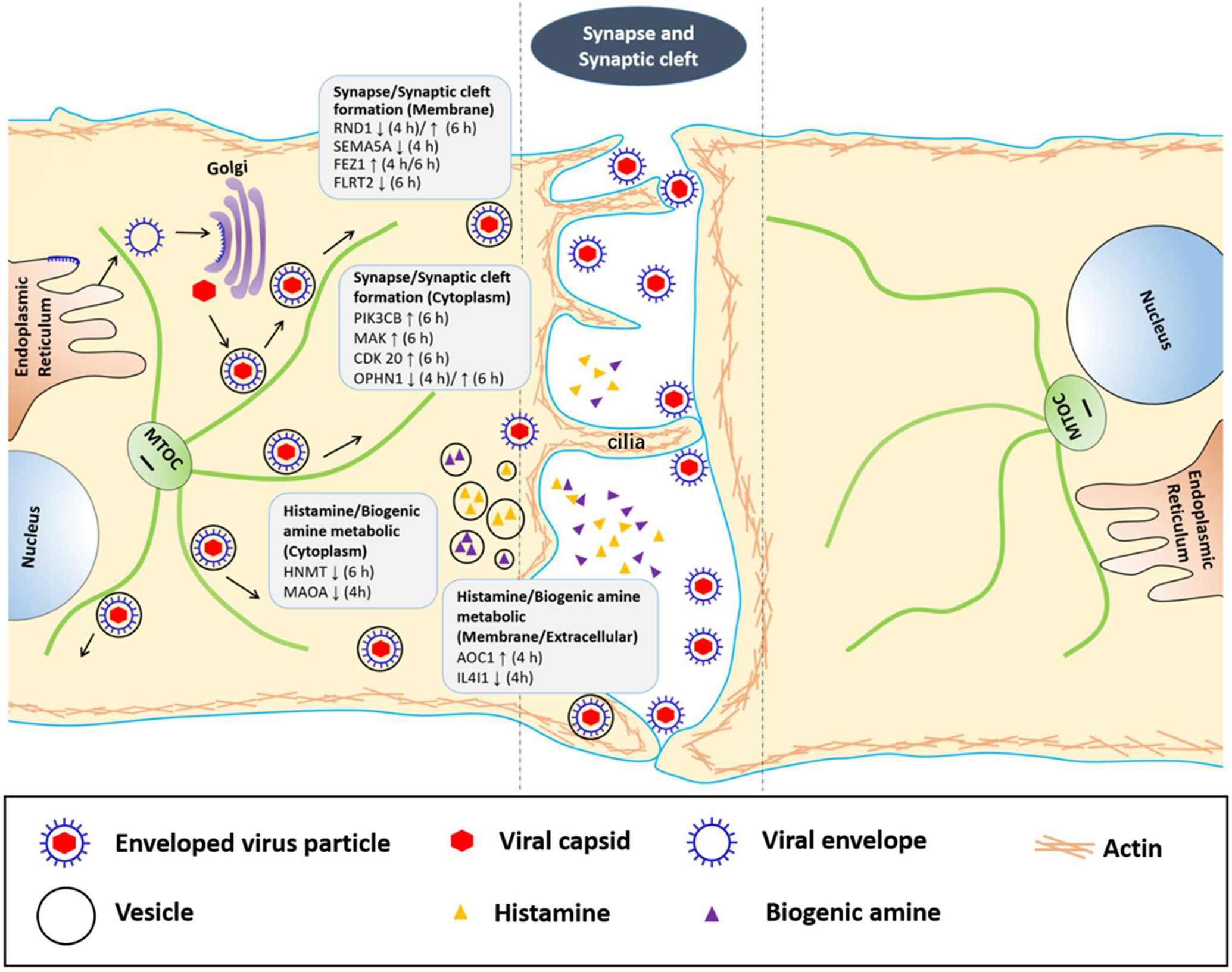
Proposed involvement of regulated genes/proteins in the process of cilia/synaptic-cleft formation and metabolism/vesicle-mediated secretion of biogenic amines (including histamine). Genes regulated by PEDV with an important role in these processes are displayed in grey boxes, with their subcellular location, regulation, up (↑) or down (↓), and time-point of regulation. Subcellular locations of genes/proteins were retrieved from UniProtKB/Swiss-Prot. data provided in the GeneCards reports of these genes/proteins. The drawing was inspired by figures presented in the publications of Molinier-Frenkel et al. (31:2019) and Sattentau (32: 2008).

A number of studies in recent years have shown that cell to-cell spread of enveloped viruses can use structures with similar characteristics as the synaptic clefts between neurons (32: reviewed in Sattentau *et al.* 2008). The space within these so-called “virological clefts” eliminates fluid-phase diffusion and facilitates more effective infection of neighboring cells. Synaptic cleft structures are also formed between immune cells, e.g. between APC’s and T cells. Transmission of immune signals by vesicle-mediated exocytosis of chemical stimuli from APC’s into the cleft space between the APC and donor immune cell or by receptor-ligand binding in this space, only activates the immune cell that forms the cleft with the APC. Regulation of genes involved in histamine/biogenic amines (see above) and formation of cilia/clefts suggested that gene expression related to this intersynaptic signaling between immune cells could be affected in response to infection with PEDV (Fig 4). In particular, the highly down-regulated gene IL4I1 (30-fold at 4h p.i.) is of interest (see also above). IL4I1 is believed to be secreted from APC’s (e.g. DC’s) in the immune cleft formed with T cells (31: Molinier-Frenkel *et al.* 2019). The mechanism how IL4I1 transmits its signal to T cells is not completely understood. It could bind to a receptor that concentrates this amino oxidase in the cleft, resulting in elevated H_2_O_2_ and ammonia production, phenylalanine depletion and phenylpyruvate production in the cleft space. These alteration in the concentration of these chemicals in the cleft are sensed by the T cell. The paralog of IL4I1, amino oxidase MAOA (down-regulated 11-fold by PEDV) could also play a similar role in this signaling process. Remarkable was also the strong down-regulation of genes coding for the Olfactory Receptor Family 2 Subfamily A Member 14 (OR2A14; 17-fold at 4h) and Anoctamin 2 (ANO2, alias CaCC;14-fold at 4h). ANO2 is a calcium-activated chloride channel imbedded in the basal membranes of neurons that harbor apical membrane receptors like OR2A14 that sense odorants. By importing chloride ions into the cytosol ANO2 contributes to the depolarization of these neurons (https://www.kegg.jp/kegg-bin/show_pathway?hsa04740+57101). Loss of smell and taste is one of the first noticeable symptoms of COVID-19. Genetic defects in the ANO2 gene are associated with Von Willebrand disease, a bleeding disorder due to defective platelet aggregation (33: Schneppenheim *et al.* 2007).

### Associations of PEDV-DEGs with COVID-19 pathology

Development of pneumonia in COVID-19 patients may progress into respiratory failure, i.e. acute respiratory distress syndrome (ARDS), often leading to septic shock and death (34: Xu *et al.* 2020). Numerous collateral complications were reported.

Failure of vital organs, like liver and kidneys, and numerous neurological complications (seizures, strokes, encephalitis and lameness symptoms [e.g. Guillain–Barré syndrome]). In addition, cardiovascular-related complications may occur, including arrhythmia, inflammation of the cardiac muscles, complete heart failure and abnormal blood clotting and thrombosis. Recent data of COVID-19 cases around the world indicate that children and adolescents are less at risk and elderly above 70 years-old and persons with underlying diseases as COPD, diabetes/obesity, hypertension and heart diseases show an increased risk for developing severe symptoms (35: Onder *et al.* 2020, 1 Wu *et al.* 2020). Also disease incidence in adult males is significantly higher than in females of the same age.

To assess whether PEDV-DEGs relate to these pathological symptoms, the genotyping program VarElect was used to identify genes matching the terms “ARDS”, “Cardiomyopathy”, “Obesity (Diabetic)”, and “Platelets activation”. Detailed information about all matching DEGs is provided in supplementary file 3 in separate sheets for all 4 queried terms. DEGs matching to more than one query term are displayed in Fig. 5. Remarkable associations of DEGs with these terms are mentioned in sections beneath.

**Fig. 5.**
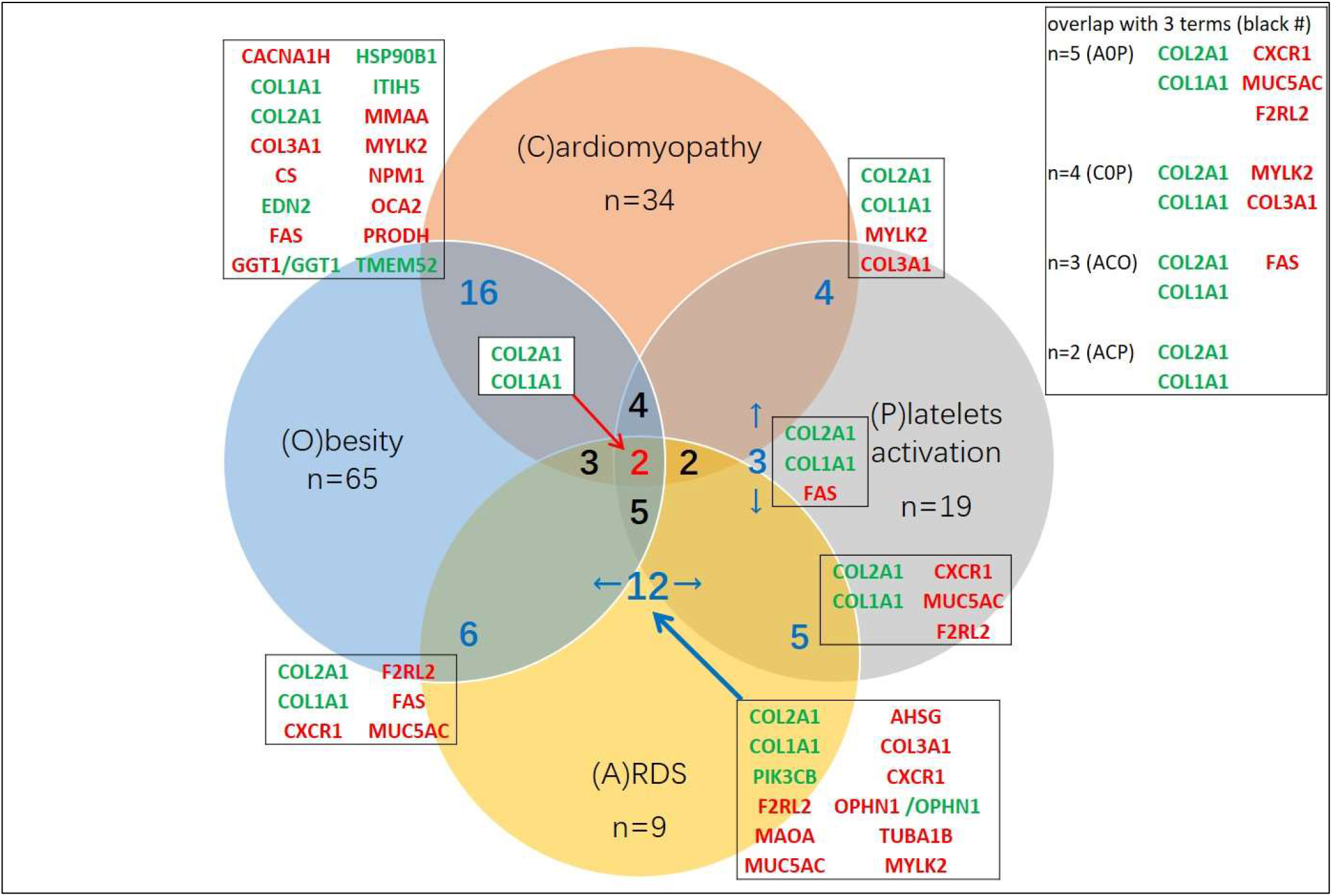
Number of PEDV-DEGs that matched to the terms “ARDS”, “Cardiomyopathy”, “Obesity (Diabetic)”, and “Platelets activation” according to the genotyping program “VarElect” (n=total per term). The number of overlapping DEGs between two or three terms are depicted in blue and black, respectively. Gene-symbols of DEGS matching 2 terms are provides in boxes in the Venn diagram and with 3 term in a separate box outside the diagram. The DEGs COL1A1 and COL21A (in the center) matched to all 4 terms (red number). Up-regulated genes are shown in green and down-regulated genes are shown in red.

#### ARDS

PEDV-DEGs coding for proteins directly associated with ARDS according to the VarElect program were i) Mucin 5AC (MUC5AC, down-regulated 14-fold at 4h), a gel forming component of the mucus layer of gastric and airway epithelial layers and which expression is regulated by IL6 or IL17, ii) transcription factor RORC (up-regulated 11-fold at 4h), iii) CXCR1, the high-affinity receptor for CXCL8 (alias IL8: down-regulated 8-fold at 4h), iv) the Thrombin-like 2 receptor F2RL2 (down-regulated 16-fold at 4h) receptor, and v) Collagens 1A (COL1A, 8-fold upregulated at 4h), 2A (COL2A, 19-fold upregulated at 6h) and 9A3 (15-fold upregulated at 6h).

See the paragraph “Regulation of immune genes” (above) for functional information about the genes RORC, F2RL2 and CXCR1. Further datamining and literature searches revealed additional associations of specific PEDV-DEGs to respiratory dysfunction. Related to the strong down-regulation of MUC5A at 4h, expression of the gene coding for the enzyme “Polypeptide N-Acetyl-galactosaminyltransferase 16 (GALNT16) was up-regulated 23-fold by PEDV at 6h. GALNT16 couples the first galactose residue of a mucin type O-linked glycan to proteins, resulting in formation of the so-called “T-antigen” (GalNaca1-Ser/Thr moiety)(36: Ju *et al.* 2011). MRV infection did not altered expression of this gene or of other GALNT orthologs.

Instead, MRV highly stimulated gene expression of the enzyme “Core 1 Synthase, Glycoprotein-N-Acetylgalactosamine 3-Beta-Galactosyltransferase 1” (C1GALT1, 26-fold at 6h). C1GALT1 couples the second galactose residue to the T-antigen moiety, further extending the mucin type O-linked glycan chain to a so-called “Tn-antigen” (Galb1-GalNaca1-Ser/Thr moiety). Somatic mutations in the gene coding for the essential chaperone of C1GALT1 (C1GALT1C1), causes Tn polyagglutination syndrome (https://omim.org/entry/300622), a disorder caused by binding of natural anti-Tn antibodies to the Tn-antigen expressed on the surface of erythrocytes and other blood cells (including platelets and leukocytes). The high elevated expression of GALNT16 may also support O-linked glycosylation of structural proteins of PEDV (37: Fung *et al.* 2018). Recently, glycosylation sites on the spike protein of SARS-CoV-2 were predicted, suggesting that the spikes of SARS-CoV-2 could bear an unique N- and O-linked glycosylation architecture different from that of SARS-CoV-1 (38: Vankadari *et al.* 2020). It was proposed that this unique glycosylation architecture plays a role in protecting SARS-CoV-2 for the hosts immune defense.

PEDV also elevated expression of genes coding for Extracellular Matrix Protein 2 (ECM2; 10-fold at 4h), and COLs 1A, 2A and 9A3 (see above for the FC of these COL genes). Prolonged deposition of ECM proteins and collagens fibers synthesized by fibroblast may lead to formation of excessive scar tissue. In the lungs this could result in pulmonary fibrosis and, eventually, to interstitial lung diseases leading to acute or chronic respiratory failure (39: Cutroneo *et al.* 2007, 40: Gagiannis *et al.* 2020).

#### Platelets activation

According to the BioGPS “Primary Cell Atlas” dataset, which provides information of expression levels measured in different type of cells/tissues, about 20 percent of the PEDV-DEGs code for proteins highly expressed in platelets and/or in cells of the vascular system. Among them, also the COL1A1, 2A1 and 9A3 genes. Binding of COLs to integrin subunit alpha on the surface of platelets mediates release of calcium from the ER into the cytosol, resulting in activation of functional processes of platelets. Processes like shape change, induced by phosphorylation of Myosin by Myosin Light Chain Kinase 2 (MYLK2; down-regulated 12-fold by PEDV), and platelets aggregation activated by Talin 1 and/or Fermitin Family Member 3 (FERMT3, alias KINDLIN3). In response to PEDV infection, an increased gene expression level was only observed for FERMT3 (18-fold at 4h), and not for Talin 1. Gene expression of the F2RL2 surface receptor in response to PEDV infection was reduced 15-fold at 6h (see also above). Binding of thrombin to this receptor also activates functional processes in platelets and promotes vasodilation and permeability of the vascular wall.

#### Cardiomyopathy

A clear association was found for the PEDV up-regulated cytokine genes coding for Cardiotrophin-1 (CTF1)(41: Wollert *et al.* 1996) and for the preproprotein of the neuroactive peptide hormone Endothelin 2 (EDN2)(42: Nagai *et al.* 2007) with diseases of the heart-muscle. The active forms of the peptide hormone EDN1 and 2 are secreted from endothelial cells and bind to the same class of G-protein coupled receptors as the Angiotensin peptide hormones (AGT1-7). EDN2 and AGT2 are vasoactive peptides and binding of EDN2 and AGT2 to their receptors on granular cells of the juxtaglomerular apparatus in the kidney raises free calcium levels in the cytosol, leading to inhibition of cAMP-mediated secretion of the aspartyl-protease renin (REN), the key regulator of renin-angiotensin-aldosterone system (RAAS) (https://www.kegg.jp/kegg-bin/show_pathway?hsa04924+1907). REN converts pre-angiotensinogen (AGT) to the endocrine peptide-hormone AGT1 (https://www.kegg.jp/kegg-bin/show_pathway?hsa04614+5972). AGT1 is further cleaved to variants with specific endocrine activity by Angiotensin I Converting Enzymes (e.g. ACE and ACE2:). On the surface of bronchial epithelial cells ACE2 was identified as entry receptor for SARS-CoV-1 and −2 (43: Hoffmann *et al.* 2020). The octamer peptide AGT2 stimulates secretion of the mineralocorticoid hormone aldosterone by the adrenal glands. Aldosterone, and the AGT and EDN peptide hormones regulate an array of physiological processes in the body, e.g. vascular smooth muscle contraction, blood pressure, fluid and electrolyte homeostasis (44: Agapitov *et al.* 2002). All processes that are important for proper functioning of the vascular system, heart muscles and kidneys.

CTF1 is directly involved in the pathology of numerous cardiovascular diseases by promoting cardiac myocyte hypertrophy (41: Wollert *et al.* 1996), which may lead to the onset of heart-diseases like “hypertrophic cardiomyopathy” or “dilated cardiomyopathy”, and eventually, to (lethal) heart failure. PEDV induced a strong down-or up-regulation of several other genes directly involved in the function of cardiomyocytes. Sodium voltage-gated channel subunit 4 (SCN4B) was strongly up-regulated (15-fold) and MYLK2 (see above), Citrate Synthase (CS; down-regulation 24-fold) and Ankyrin Repeat Domain 1 (ANKRD1) were strongly down-regulated.

Down-regulation of CS may reduce oxidative capacity in cardiomyocytes. Gene expression of ANKRD1 was down-regulated 12-fold in response to MRV infection at 4 h.p.i, but reverted to a 24-fold up-regulation 2h later. ANKRD1 is a putative transcription factor involved in regulation of gene expression in hypertrophic myocytes (https://www.wikipathways.org/index.php/Pathway:WP516)

Regulation of cytosolic calcium levels in the cytosol of cardiomyocytes, e.g. by binding of COL1A1 and COL2A (up-regulated by PEDV, see above) to integrin subunit alpha on the surface of cardiomyocytes or after import of calcium ions mediated by calcium voltage-gated channels (e.g. by CACNA1H; down-regulated 5-fold at 4h by PEDV) may also trigger myocyte hypertrophy. PEDV strongly down-regulated gene expression of a potassium voltage-gated channel (KCNQ4;16-fold). This in contrast to a strong up-regulation of the sodium symporter SCN4B. For the potassium channel CACNA1H (alias Kv7.4) it was reported that this channel regulates the membrane potential and Ca^2+^ permeability of mitochondria located in the vicinity the sarcoplasmic reticulum in rat cardiomyocytes (45: Testai *et al.* 2016). All three above mentioned ion channels are also involved in the process of excitation, contraction/relaxation and repolarization of cardiac myocytes. MRV also down-regulated gene expression 3 to 4-fold for the potassium (KCNQ4) and calcium channel CACNA1H, but did not increased expression of the sodium symporter gene SCN4B or orthologs of this gene.

### Obesity/Diabetic

Chronic hypertension and heart disease/failure is a complication frequently observed in obese/diabetic patients. In accordance with this, 16 out of the 65 PEDV DEGs matching the term “Obesity” also matched with the term “Cardiomyopathy” (see Fig.5 and supplementary file 3, sheet “A-C-O-P). The gene coding for receptor activity modifying protein 1 (RAMP1; a calcitonin-like receptor) matching term “Obesity” was 12-fold up-regulated in response to PEDV. Binding of the vasodilator adrenomedullin (ADM) to this receptor inhibits MYLK2-mediated contraction of vascular smooth muscle cells (46: Terata *et al.* 2000). The neuropeptide hormone ADM is, inter ilia, expressed in lungs, adrenal glands, kidney, heart and vascular smooth muscle cells. It was reported that paracrine and autocrine effects of ADM contribute to prevent cardiovascular and renal damage (47: Dobrzynski *et al.* 2002 48: Tsuruda *et al.* 1998). Increased expression of ADM is also observed in blood monocytes in Kawasaki disease (KD; https://www.malacards.org/card/kawasaki_disease?). This acute inflammatory disorder of the blood vessels is observed in hospitalized children with SARS-CoV-2.

Susceptibility to KD is associated with a polymorphism in the Inositol-Trisphosphate 3-Kinase C gene (ITPKC). ITPKC is important for activation of the phosphatidylinositol 3-kinase-AKT signaling. A signal transduction cascade that steers an array of cellular processes, and in which the enzyme activities of phosphoinositide 3-kinases (e.g. PIK3CB, 3-fold up-regulated by PEDV) are executers of this cascade. (https://www.kegg.jp/kegg-bin/show_pathway?hsa04151+5291). Related to this, expression of the gene coding for “ES Cell Expressed Ras” (ERAS; a paralog of NRAS), an activator of PIK3CB, was 13-fold down-regulated in response to PEDV infection.

The gene coding for H4 histone 16 (H4-16) was 15-fold up-regulated by PEDV at 6 h.p.i. H4-16 was found to be associated with familial hyperinsulinemia and hypoglycemia in infants, and disturbance of modifications of core histones (including H4-16) were also observed in diabetic hearts (49: Yerra *et al.* 2018, 50: Krishnan *et al.* 2011). Like other core histones, H4-16 is also an autoantigen in autoimmune diseases. Expression of the genes citrate synthase (CS) and 6-Phosphofructo-2-Kinase/Fructose-2,6-Biphosphatase 1 (PFKFB1), both coding for enzymes directly involved in glucose homeostasis, were down-regulated by PEDV. Enzyme activity of PFKFB1 stimulates glycolysis and inhibits gluconeogenesis and is a key-regulator of glucose homeostasis. Expression of the gene coding for transcription factor “single-minded BHLH Transcription Factor 2” (SIM2) was 10-fold up-regulated by PEDV at 6h. SIM transcription factors act in the paraventricular nucleus, a group of neurons in the hypothalamus regulating energy homoeostasis and the feeding response. SIM1 and 2 variants with low transcriptional activity were identified in persons with early-onset obesity (51: Hung *et al.* 2007). However, the mechanism how this transcriptional regulation by SIM’s affects feeding and energy homeostasis is not known. With regard to neurological regulation by the hypothalamus of energy homoeostasis and feed intake, PEDV also up-regulated (16-fold) expression of the gene coding for the “Protein Phosphatase, Mg2+/Mn2+ Dependent 1E (PPM1E) at 6 h.p.i.. PPM1E deactivates the AMPK-kinase (Protein Kinase AMP-Activated Catalytic Subunit Alpha 1), a kinase that stimulates secretion of neuroendocrine peptides (NP’s) that regulate lipogenesis and cholesterol metabolism in the body. PEDV highly up-regulated (11-fold at 4h) gene expression of Neuropeptide B (NPB). Besides feed intake, energy homeostasis and enhancement of cortisol secretion in adrenocortical cells, NPB also influences the hypothalamic-pituitary-adrenal stress responses and increases heart rate and blood pressure. Moreover, recent studies have revealed that expression of NPB by, yet, not identified types of neurons in the Japanese rice fish (Medaka) is female-specific, i.e. is mediated only by binding of ovarian sex hormones (estrogens) to receptors of the androgen/estrogen signaling cascade (52: Hiraki-Kajiyama *et al.* 2019).

### Additional remarkable PEDV-DEGs

Highly up- or down-regulated PEDV-DEGs not mentioned in the text, and to our opinion interesting with regard to coronavirus infection, are briefly described in Table 2. Among these DEGs several genes coding for transcription factors and genes transcribed in antisense RNA’s that inhibit translation of their coding counterparts. For more functional information about these DEGs we refer to the weblinks provided in supplementary file 3, sheet “PEDV KEY DEGs”.

**Table 2.** Remarkable PEDV-DEGS not mentioned in the text

| Description | GENESYMBOL | h.p.i | FC (up or down ) | function |
| --- | --- | --- | --- | --- |
| Keratin 25 | KRT25 | 6 | 13.7 | cornified envelope formation |
| Late Cornified Envelope 3B | LCE3B | 6 | 16.3 | cornified envelope formation |
| endosome-lysosome associated apoptosis and autophagy regulator family member 2 | KIAA1324L | 0 / 6 | -2.3/16.3 | estrogen-induced gene, regulating autophagy and promotes cell survival under stress |
| NADPH Oxidase 3 | NOX3 | 6 | -11.7 | generation of superoxide, stress fiber formation |
| Fibrinogen Like 1 | FGL1 | 6 | -21.8 | immune suppressive, inhibits antigen-specific T-cell activation |
| CTBP1 Divergent Transcript (CTBP1-antisense transcript) | CTBP1-DT | 4 | -23.2 | inhibition CTBP1 mRNA translation; CTBP1=co-repressor notch-mediated transcription |
| INHBA antisense RNA 1 | INHBA-AS1 | 0 / 4 | -26.6/16.3 | inhibition INHBA mRNA translation (INHBA=cytokine TGF family) |
| Purinergic Receptor P2X 5 | P2RX5 | 6 | 16.2 | ligand-(ATP)-gated ion channel family for calcium import in cells |
| Aldo-Keto Reductase Family 1 Member C1 | AKR1C1 | 6 | -11.6 | lipid and steroid hormone metabolism |
| Apolipoprotein L2 | APOL2 | 6 | 15.4 | lipid and steroid hormone transport |
| Carboxylesterase 1 | CES1 | 6 | 15.4 | enzyme in storage lipid droplets hydrolyzing cholesteryl esters to cholesterol (prevents forming of macrophage foam cells). |
| Protocadherin beta 12 | PCDHB12 | 0 / 6 | -36.1/-5.3 | calcium-dependent cell-adhesion protein |
| Neurotrimin | NTM | 7 | -17.5 | synthesis of GPI-anchored proteins |
| Kruppel like factor 14 | KLF14 | 0 / 6 | -4.9/-17.7 | transcription co-repressor TGF-genes |
| Hes Family BHLH Transcription Factor 7 | HES7 | 6 | 10.3 | transcription factor (expression regulated by notch-signaling) |
| POU Class 2 Homeobox 3 | Pou2f3 | 6 | 9.3 | transcription factor, notch downstream signaling |
| Homeobox A7 | HOXA7 | 4 | 29.0 | transcription factor; active in basophils |
| Forkhead Box Q1 | FOXQ1 | 6 | 15.1 | transcription factor; active in NK and CD8+ T cells |
| Y-Box Binding Protein 1 | YBX1 | 4 | -15.5 | transcription factor; cold shock domain protein, activated by MAPK (ERK) signaling. |

## Discussion

In this report we measured the transcriptional response of Vero cells shortly after infection with the coronavirus PEDV. The function of the responding host genes and the biological processes in which they act were studied in detail by us to find plausible relations to COVID-19 pathology. Because of the differences in genomic organization and expression of viral proteins between SARS-CoV-2 and PEDV, we paid less attention to couple the response of specific host genes to the function of specific coronavirus proteins.

We were able to infect the majority of Vero cells (>50%) with PEDV and MRV synchronically. This resulted in a unique set of highly up- and down-regulated DEGs for PEDV. Not more than 14% of the 266 PEDV-DEGs (n=37) were similar to MRV-DEGS (total of 727 MRV-DEGs). In contrast to MRV, we observed no typical response of antiviral genes and related cytokine/chemokine genes in Vero cells within 6 h.p.i. for PEDV. For MRV these processes started already before 4 h.p.i.. We have to notice that PEDV replication started 2h later than MRV3 replication, which could in part be the reason for not detecting transcriptional regulation of specific cytokine, chemokine and antiviral genes for PEDV. Longer incubation times than 6h were not planned in the original design of our experiments and would have resulted in a set of PEDV-DEGs dominated by genes involved in syncytia forming and apoptotic/necrotic cell death. Nevertheless, at 6h replication of PEDV RNA was detected by RT-PCR, indicating that dsRNA was present in the cells and could have be sensed by cytosolic pattern recognition receptors of the RIG-I-like family to initiate an antiviral and related cytokine/chemokine response. Similar as observed in another study with PEDV and Vero cells, and in analogy with SARS-CoV-1 and −2, we observed a high up-regulation of the transmembrane serine protease gene (TMPRSS13) that acts as a co-factor in the infection process of cells (25: Matsuyama *et al.* 2010). Therefore, we are confident that PEDV infection delayed or suppressed an early antiviral response and related cytokine/chemokine response, similar as was observed in intestinal epithelial cells (53: Cao *et al.* 2015). In recent *in vivo* studies with SARS-CoV-2, a delayed IFN I response was observed accompanied by only low levels of type I and III IFNs expression and a reduced or relatively late (not before 24 h.p.i.) antiviral response (19: Blanco-Melo *et al.* 2020, 20: Lamers *et al.* 2020). We observed a reduced expression of EEF1A1, as part of a transcription factor-complex that binds and activates the promotor of IFNγ, and of the cytokine EDA and its receptor (EDARADD) involved in activation of canonical-NFKB transcription of antiviral cytokine-chemokine genes like CXCL8 (alias IL8) an CXCL10. This reduced expression of EEF1A1 and EDA and its receptor may play a crucial role in delaying or downgrading an IFN-mediated antiviral and cytokine/chemokine response in our Vero cell system.

The elevated transcription of many cytokine and chemokine genes in Vero cells by MRV suggests that replication of this RNA virus in epithelial cells induces secretion of these immune effectors (more information about the genes/processes that responded to MRV infection will be published elsewhere). PEDV, and also the MRV3 strain we used both replicate in enterocytes lined up in the intestinal mucosal layer. In the intestinal and bronchial epithelial layer, microfold (M) cells are imbedded between these lined up epithelial cells. M-cells sense and engulf foreign pathogens/antigens from the lumen to present them to residing immune cells.

According to pathway analysis, most T cell related immune genes regulated in response to MRV infection were part of the pathways “T-helper 17 (Th17) differentiation/activation” and “IL17 signaling”. Antigen presentation by M cells to Th17 cells in the submucosal layers stimulate secretion of different types of IL17 cytokines (IL17A-D, IL25 and Il17F) resulting in activation of different types of innate immune cells and T cells, including Th1/Th2 cells. Dysregulation of this pathogen-induced IL17 response may disturb the balance between Th1/Th2-cell mediated immune responses, resulting in excessive inflammation, damage to the epithelial layer and on the long term, to autoimmune reactions. TF RORC (or specific isoforms of this TF, see above) plays a pivotal in controlling IL17 expression and secretion by Th17 cells. PEDV strongly up-regulated gene expression of RORC in Vero cells whereas MRV down-regulated expression of this TF. This difference in regulation of TF RORC suggests that virus-induced activation or suppression of IL17 secretion by Th17 cells in submucosal layers of airway epithelium can be an important mechanism to dysregulate the activation of T cell responses. Therefore, TF RORC might be a potential target for drug treatment/development. Drugs affecting expression of RORC, like the fluorinated steroid “Dexamethasone” and the synthetic tetracycline derivative “Doxycycline” (http://ctdbase.org/basicQuery.go?bqCat=gene&bq=RAR+related+orphan+receptor+C) are already under investigation in relation to SARS-CoV-2 pathogenesis. Transcriptional regulation of a set of genes coding for enzymes involved in biogenic amine metabolism was unique for PEDV, and not observed for MRV. Most associations of these PEDV-DEGs were found with histamine, a compound produced by mast cells and basophils, and released by these cells in response to allergens and pathogen-induced inflammation. The 10-fold up-regulation of the enzyme AOC1 suggests that histamine is converted to imidazole-acetaldehyde (see Fig. 3).

However, without data of intra- and extracellular concentrations of the chemicals this remains speculative. Recent reports indicated that submucosal mast cells in the lungs were triggered by SARS-CoV-2 infection to release pro-inflammatory cytokines (IL1, IL6 and TNF-alpha) as well as histamine, leukotrienes (LCD4) and prostaglandins (PGD2) (54: Kritas *et al.* 2020). This release by mast cells or basophils can induce excessive inflammation in the lungs and bronchoconstriction, leading to acute respiratory failure. As proposed by Raymond *et al.* (55: 2020), testing of existing antihistamines and mast cell/basophil stabilizers in clinical trials to evaluate their effectiveness as therapeutic drug for COVID-19, has potential.

An interesting finding was the strong down-regulation of the gene coding for the amino acid oxidase IL4I1 4 h.p.i with PEDV. IL4I1 enzyme activity plays an important role in “chemical signaling” in the cleft space between APCs and immune effector cells. Recent studies showed that down-regulation of IL4I1 expression resulted in inhibition of differentiation of monocytic myeloid-derived suppressor cells (Mo-MDSC) to polymorphonuclear MDSC (PMN-MDSC)(56: Gao *et al.* 2019). Mo- and PMN-MDSCs use different mechanisms to suppress the activity of antigen-specific T cells, NK cells, DCs and macrophages. Levels of MDSCs increase in the spleen, lymph nodes, or in blood in response to inflammation of tissues or tumors in the body (57: Movahedi *et al.* 2008). MDSCs infiltrate these tumors and inflamed tissues to suppress the local activity of specific immune cells. Therefore, the role of infiltrating MDSCs at inflammatory sites in the lungs of COVID-19 patients, as part of an SARS-CoV-2 immune-evading strategy, and the role of IL4I1 in this process, is worthwhile to investigate in more detail.

Expression of genes that promote or inhibit the formation and motility function of cilia were time-dependently regulated by PEDV. The 20-fold up-regulation of FEZ1 gene expression at 4h (20-fold) descended within 2h to a moderate 6-fold up-regulation. This descend occurred simultaneously with elevation of gene expression of the kinases MAK and CDK20, both involved in inhibition of cilia formation. Because PEDV efficiently replicates in enterocytes that carry ciliated membrane protrusions (microvilli) on their luminal surface, regulation of these genes could be related to structural changes in the cytoskeleton of cells imposed by virus replication (e.g. syncytia forming in PEDV infected Vero cultures). Likewise, SARS-CoV-2 replication could also impose structural changes in cilia protruding from the surface of upper-airway epithelial cells (nose, trachea) and bronchi. Based on our results we cannot pinpoint a specific process in which these cilia-regulating genes act. Possible processes can either be formation of an immune cleft, a virological cleft to promote more effective infection of neighboring cells, or cytoskeleton rearrangements to support virus replication, morphogenesis and budding from cells. Interestingly, two recent studies revealed a high level of expression of the SARS-CoV-2 entry receptor ACE2 and its co-receptor TMPRSS2 in ciliated airway epithelial cells (58: Sungnak *et al.* 2020), especially in bronchial “secretory cells” (59: Lukassen *et al.* 2020). These latter cells are an intermediate population of cells in the differentiation process of club or goblet cells to terminal ciliated cells. Motile cilia on the surface of nasal and bronchial epithelial cells play an important role in mechanical clearance of hostile particles from the airways, and SARS-CoV-1 and 2 may benefit from dysfunction of this clearance system. Inhibition of cilia formation related to this mechanical clearance and inhibition of formation of immune cleft are relatively unexplored strategies of respiratory RNA viruses to evade the host defense (60: Ma *et al.* 2015). These processes deserve more attention, and may also be considered as possible target-processes for interference with drugs.

Within our set of PEDV-DEGs, and the biological processes deduced from this set, we found associations with diverse aspects of COVID-19 pathogenesis, i.e. “ARDS”, “Cardiomyopathy”, “Obesity (Diabetic)”, and “Platelets activation”. However, it is unknown whether the proteins encoded by these PEDV-DEGS indeed play a role in the biological processes underlying the symptoms and complications observed in hospitalized persons infected with SARS-CoV-2. Nevertheless, a part of these genes/processes may be starting points for further dedicated research. Research to fine tune drug treatment protocols that are already applied for COVID-19 patients, or research that provides new insights for treatments with alternative prophylactic and therapeutic approved drugs. In Table 3. we summarized the PEDV-DEGs that are, to our opinion, of interest for modulation of the biological processes underlying the pathogenesis of COVID-19.

**Table 3.** Possible target genes for COVID-19 therapy.

| Description | GENESYMBOL | regulation of expression | functional relation to COVID-19 pathogenesis |
| --- | --- | --- | --- |
| Eukaryotic Translation Elongation Factor 1 Alpha 1 | EEF1A1 | down | inhibition IFNγ production |
| Heat Shock Protein Family A (Hsp70) Member 8 | HSPA8 | down | inhibition viral replication / antigen presentation |
| Olfactory Receptor Family 2 Subfamily A Member 14 | OR2A14 | down | loss of smell and taste |
| Anoctamin 2 | ANO2 | down | loss of smell and taste |
| histamine N-Methyltransferase | HNMT | down | (methyl)-histamine metabolism, mast cell and basophils response |
| Amine Oxidase Copper Containing 1 | AOC1 | up | (methyl)-histamine metabolism, mast cell and basophils response |
| Monoamine Oxidase A | MAOA | down | (methyl)-histamine metabolism / immune signaling via cleft |
| Interleukin 4 Induced 1 | IL4I1 | down | immune signaling via cleft |
| Male Germ Cell Associated Kinase | MAK | up | viral spread or immune signaling via cleft/changes in cilia of epithelial cells. |
| Fasciculation And Elongation Protein Zeta 1 | FEZ1 | up | viral spread or immune signaling via cleft/changes in cilia of epithelial cells. |
| RAR Related Orphan Receptor C | RORC | up | T(h17) cell mediated immune response |
| Cardiotrophin 1 | CTF1 | up | cardiomyopathy |
| Endothelin 2 | EDN2 | up | cardiomyopathy and vascular dysfunction |
| Myosin Light Chain Kinase 2 | MYLK2 | down | cardiomyopathy |
| Receptor Activity Modifying Protein 1 | RAMP1 | up | cardiovascular and renal damage |
| Neuropeptide B | NPB | up | increase heart rate and blood pressure |
| Protein Phosphatase, Mg<sup>2+</sup>/Mn<sup>2+</sup> Dependent 1E | PPM1E | up | increase heart rate and blood pressure |
| Mucin 5AC, Oligomeric Mucus/Gel-Forming | MUC5AC | down | loss of mucosal barrier function |
| Polypeptide N-Acetylgalactosaminyltransferase 16 | GALNT16 | up | loss of mucosal barrier function /polyagglutination syndrome erythrocytes |
| Collagen Type II Alpha 1 Chain | COL2A1 | up | lung fibrosis/cardiomyopathy |
| Collagen Type I Alpha 1 Chain | COL1A1 | up | lung fibrosis/cardiomyopathy |
| Coagulation Factor II Thrombin Receptor Like 2 | F2RL2 | down | thrombosis/vascular dysfunction |
| Fermitin Family Member 3 (KINDLIN3) | FERMT3 | up | thrombosis/vascular dysfunction |

## Colophon

The overwhelming amount of data published recently made it impossible for us to oversee all (novel) facts about the SARS-CoV-2 virus and pathology of the COVID-19 disease. Some important genes and related processes imbedded in our set of PEDV-DEGs may have been overlooked by us. Therefore, we encourage researchers, especially medical, immunological and pharmaceutical specialists, to study this set of DEGs in detail. The users-friendly supplementary file 3 with functional information about DEGs and related biological processes can be down-loaded from the web. By publishing these PEDV data ahead of our complete study, we hope that some of the gene targets and cognate processes we have identified for the coronavirus PEDV will contribute to a better understanding how hospitalized COVID-19 patients can be treated and cured. A more condensed version of this manuscript, focusing on the original goal of our study, will be submitted to a peer-reviewed virological journal soon.

## Supporting information

material and methodes

RNAseq next generation sequencing report

RNAseq data analysis rapport

Excel file with functional information

## Authors’ contributions

WH and MH designed and arranged the *in vitro* infection experiments with Vero cells. WH performed the *in vitro* infection experiments. RNAseq analysis, mapping of reads to the reference genome of Cercopithecus aethiops and generation of RNAseq datafiles of pairwise comparisons was performed by the firm GenomeScan B.V. (Leiden, The Netherlands). MH extracted DEGs from RNAseq data files. MH and WH performed the GSEA, functional bioinformatic analysis, and datamining. MH took the lead in writing the manuscript together with WH. FL and WvdP facilitated the research project and advised about the design and execution of the experiments. All authors reviewed the manuscript and approved the final version.

## Supplementary files

*Supplementary file 1:* Materials and methods cell culture and virus propagation, infection experiment, RT-PCR detection, Immune peroxidase monolayer assay (IPMA), RT-PCR detection, Preparation of RNA pools and quality analysis of RNA. *Supplementary file 2a*: RNAseq next generation sequencing report GenomeScan (Leiden, The Netherlands).

*Supplementary file 2b*: RNAseq data analysis rapport GenomeScan (Leiden, The Netherlands).

*Supplementary file 3*: Interactive Excel file with sortable tables in separate sheets. Please, first read the sheet “read me” for an explanation and instructions for the use of the tables. Excel sheets contain tables with i) PEDV and MRV DEGs extracted from RNAseq data files, ii) functional information about the PEDV-DEGs, iii) GSEA extracted pathways (for MRV and PEDV), GO-terms (only for PEDV) and compound associations (only for PEDV), and iv) associations of PEDV-DEGs with the terms “cytokines-chemokines”, “(anti)-viral”, “Biogenic amines”, “cilia and synaptic clefts”, and the disorders “ARDS, “Cardiomyopathy”, “Obesity”, and “Platelets activation” (A-C-O-P).

## References

1) Wu C, Chen X, Cai Y, et al. Risk Factors Associated With Acute Respiratory Distress Syndrome and Death in Patients With Coronavirus Disease 2019 Pneumonia in Wuhan, China [published online ahead of print, 2020 Mar 13]. JAMA Intern Med. 2020;180(7):1–11. doi:10.1001/jamainternmed.2020.0994

2) Pensaert, M., de Bouck, P., 1978. A new coronavirus-like particle associated with diarrhea in swine. Arch Virol. 58:243–247.

3) Huang YW, Dickerman AW, Piñeyro P, et al. Origin, evolution, and genotyping of emergent porcine epidemic diarrhea virus strains in the United States. mBio. 2013;4(5):e00737–13. Published 2013 Oct 15. doi:10.1128/mBio.00737-13

4) Jung K, Saif LJ., Porcine epidemic diarrhea virus infection: Etiology, epidemiology, pathogenesis and immunoprophylaxis. Vet J. 2015;204(2):134–143. doi:10.1016/j.tvjl.2015.02.017

5) Thimmasandra Narayanappa A, Sooryanarain H, Deventhiran J, et al. A novel pathogenic Mammalian orthoreovirus from diarrheic pigs and Swine blood meal in the United States. mBio. 2015;6(3):e00593–15. Published 2015 May 19. doi:10.1128/mBio.00593-15

6) Lelli D, Beato MS, Cavicchio L, et al. Virol J. 2016;13:139. Published 2016 Aug 12. doi:10.1186/s12985-016-0593-4

7) Hulst M, Hakze-van der Honing R, Vastenhouw S, et al. Identification of a Novel Reassortant of a Mammalian Orthoreovirus in Faeces of diarrheic pigs in the Netherlands: presentation at the 11th annual meeting of EPIZONE in Paris, 2017; abstract C9 https://www.epizone-eu.net/upload_mm/3/a/0/69d0a5dc-6eae-48dd-895a-5c2e89eb2476_EPIZONE2017%20Abstract%20book.pdf.

8) Steyer A, Gutiérrez-Aguire I, Kolenc M, et al. High similarity of novel orthoreovirus detected in a child hospitalized with acute gastroenteritis to mammalian orthoreoviruses found in bats in Europe. J Clin Microbiol. 2013;51(11):3818–3825. doi:10.1128/JCM.01531-13

9) Tyler KL. Pathogenesis of reovirus infections of the central nervous system. Curr Top Microbiol Immunol. 1998;233(Pt 2):93–124. doi:10.1007/978-3-642-72095-6_6

10) Chua KB, Voon K, Crameri G, et al. Identification and characterization of a new orthoreovirus from patients with acute respiratory infections. PLoS One. 2008;3(11):e3803. doi:10.1371/journal.pone.0003803.

11) Cheng P, Lau CS, Lai A, et al. A novel reovirus isolated from a patient with acute respiratory disease. J Clin Virol. 2009;45(1):79–80. doi:10.1016/j.jcv.2009.03.001

12) Duan Q, Zhu H, Yang Y, Li WH, Zhou YS, He J. ReoV isolated from SARS patients. Chin Sci Bull. 2003;48:1293–6.

13) Zuo T, Tan H, He J, Zhu H, Zhang H, et al. (2003) Cloning and identification of reovirus isolated from specimens of SARS patients. Bull Acad Mil Med Sci 27:241–243.

14) Huang C, Liu WJ, Xu W, et al. A Bat-Derived Putative Cross-Family Recombinant Coronavirus with a Reovirus Gene. PLoS Pathog. 2016;12(9):e1005883. Published 2016 Sep 27. doi:10.1371/journal.ppat.1005883.

15) Obameso JO, Li H, Jia H, et al. The persistent prevalence and evolution of cross-family recombinant coronavirus GCCDC1 among a bat population: a two-year follow-up. Sci China Life Sci. 2017;60(12):1357–1363. doi:10.1007/s11427-017-9263-6.

16) Brian DA, Baric RS. Coronavirus genome structure and replication. Curr Top Microbiol Immunol. 2005;287:1–30. doi:10.1007/3-540-26765-4_1

17) Comparative tropism, replication kinetics, and cell damage profiling of SARS-CoV-2 and SARS-CoV with implications for clinical manifestations, transmissibility, and laboratory studies of COVID-19: an observational study. Hin Chu*, Jasper Fuk-Woo Chan*, Terrence Tsz-Tai Yuen*, Huiping Shuai*, Shuofeng Yuan, Yixin Wang, Bingjie Hu, Cyril Chik-Yan Yip, Jessica Oi-Ling Tsang, Xiner Huang, Yue Chai, Dong Yang, Yuxin Hou, Kenn Ka-Heng Chik, Xi Zhang, Agnes Yim-Fong Fung, Hoi-Wah Tsoi, Jian-Piao Cai, Wan-Mui Chan, Jonathan Daniel Ip, Allen Wing-Ho Chu, Jie Zhou, David Christopher Lung, Kin-Hang Kok, Kelvin Kai-Wang To, Owen Tak-Yin Tsang, Kwok-Hung Chan, Kwok-Yung Yuen. Lancet Microbe 2020; 1: e14–23.

18) Lamers MM, Beumer J, van der Vaart J, et al. SARS-CoV-2 productively infects human gut enterocytes. Science. 2020;369(6499):50–54. doi:10.1126/science.abc1669.

19) Osada N, Kohara A, Yamaji T, et al. The genome landscape of the african green monkey kidney-derived vero cell line. DNA Res. 2014;21(6):673–683. doi:10.1093/dnares/dsu029.

20) Chew T, Noyce R, Collins SE, Hancock MH, Mossman KL. Characterization of the interferon regulatory factor 3-mediated antiviral response in a cell line deficient for IFN production. Mol Immunol. 2009;46(3):393–399. doi:10.1016/j.molimm.2008.10.010

21) Blanco-Melo D, Nilsson-Payant BE, Liu WC, et al. Imbalanced Host Response to SARS-CoV-2 Drives Development of COVID-19. Cell. 2020;181(5):1036–1045.e9. doi:10.1016/j.cell.2020.04.026.

22) Rasmussen TB, Boniotti MB, Papetti A, et al. Full-length genome sequences of porcine epidemic diarrhoea virus strain CV777; Use of NGS to analyse genomic and sub-genomic RNAs. PLoS One. 2018;13(3):e0193682. Published 2018 Mar 1. doi:10.1371/journal.pone.0193682.

23) Shi W, Fan W, Bai J, et al. TMPRSS2 and MSPL Facilitate Trypsin-Independent Porcine Epidemic Diarrhea Virus Replication in Vero Cells. Viruses. 2017;9(5):114. Published 2017 May 18. doi:10.3390/v9050114.

24) Zmora P, Blazejewska P, Moldenhauer AS, et al. DESC1 and MSPL activate influenza A viruses and emerging coronaviruses for host cell entry. J Virol. 2014;88(20):12087–12097. doi:10.1128/JVI.01427-14.

25) Matsuyama S, Nagata N, Shirato K, Kawase M, Takeda M, Taguchi F. Efficient activation of the severe acute respiratory syndrome coronavirus spike protein by the transmembrane protease TMPRSS2. J Virol. 2010;84(24):12658–12664. doi:10.1128/JVI.01542-10.

26) Rauen T, Juang YT, Hedrich CM, Kis-Toth K, Tsokos GC. A novel isoform of the orphan receptor RORγt suppresses IL-17 production in human T cells. Genes Immun. 2012;13(4):346–350. doi:10.1038/gene.2011.85.

27) de Groot RJ. Structure, function and evolution of the hemagglutinin-esterase proteins of corona- and toroviruses. Glycoconj J. 2006;23(1-2):59–72. doi:10.1007/s10719-006-5438-8

28) Chen Y, Cai H, Pan J, et al. Functional screen reveals SARS coronavirus nonstructural protein nsp14 as a novel cap N7 methyltransferase. Proc Natl Acad Sci U S A. 2009;106(9):3484–3489. doi:10.1073/pnas.0808790106.

29) Daffis S, Szretter KJ, Schriewer J, et al. 2’-O methylation of the viral mRNA cap evades host restriction by IFIT family members. Nature. 2010;468(7322):452–456. doi:10.1038/nature09489.

30) Boulland ML, Marquet J, Molinier-Frenkel V, et al. Human IL4I1 is a secreted L-phenylalanine oxidase expressed by mature dendritic cells that inhibits T-lymphocyte proliferation. Blood. 2007;110(1):220–227. doi:10.1182/blood-2006-07-036210

31) Molinier-Frenkel V, Prévost-Blondel A, Castellano F. The IL4I1 Enzyme: A New Player in the Immunosuppressive Tumor Microenvironment. Cells. 2019;8(7):757. Published 2019 Jul 20. doi:10.3390/cells8070757.

32) Sattentau Q. Avoiding the void: cell-to-cell spread of human viruses. Nat Rev Microbiol. 2008;6(11):815–826. doi:10.1038/nrmicro1972.

33) Schneppenheim R, Castaman G, Federici AB, et al. A common 253-kb deletion involving VWF and TMEM16B in German and Italian patients with severe von Willebrand disease type 3. J Thromb Haemost. 2007;5(4):722–728. doi:10.1111/j.1538-7836.2007.02460.x

34) Xu Z, Shi L, Wang Y, et al. Pathological findings of COVID-19 associated with acute respiratory distress syndrome [published correction appears in Lancet Respir Med. 2020 Feb 25;:]. Lancet Respir Med. 2020;8(4):420–422. doi:10.1016/S2213-2600(20)30076-X.

35) Onder G, Rezza G, Brusaferro S. Case-Fatality Rate and Characteristics of Patients Dying in Relation to COVID-19 in Italy [published online ahead of print, 2020 Mar 23]. JAMA. 2020;10.1001/jama.2020.4683. doi:10.1001/jama.2020.4683.

36) Ju T, Otto VI, Cummings RD. The Tn antigen-structural simplicity and biological complexity. Angew Chem Int Ed Engl. 2011;50(8):1770–1791. doi:10.1002/anie.201002313.

37) Fung TS, Liu DX. Post-translational modifications of coronavirus proteins: roles and function. Future Virol. 2018;13(6):405–430. doi:10.2217/fvl-2018-0008.

38) Vankadari N, Wilce JA. Emerging WuHan (COVID-19) coronavirus: glycan shield and structure prediction of spike glycoprotein and its interaction with human CD26. Emerg Microbes Infect. 2020;9(1):601–604. Published 2020 Mar 17. doi:10.1080/22221751.2020.1739565.

39) Cutroneo KR, White SL, Phan SH, Ehrlich HP. Therapies for bleomycin induced lung fibrosis through regulation of TGF-beta1 induced collagen gene expression. J Cell Physiol. 2007;211(3):585–589. doi:10.1002/jcp.20972

40) Daniel Gagiannis, Julie Steinestel, Carsten Hackenbroch, Michael Hannemann, Vincent G Umathum, Niklas Gebauer, Marcel Stahl, Hanno M Witte, Konrad Steinestel. COVID-19-induced acute respiratory failure: an exacerbation of organ-specific autoimmunity? medRxiv 2020.04.27.20077180; doi:https://doi.org/10.1101/2020.04.27.20077180.

41) Wollert KC, Taga T, Saito M, et al. Cardiotrophin-1 activates a distinct form of cardiac muscle cell hypertrophy. Assembly of sarcomeric units in series VIA gp130/leukemia inhibitory factor receptor-dependent pathways. J Biol Chem. 1996;271(16):9535–9545. doi:10.1074/jbc.271.16.9535.

42) Nagai T, Ogimoto A, Okayama H, et al. A985G polymorphism of the endothelin-2 gene and atrial fibrillation in patients with hypertrophic cardiomyopathy. Circ J. 2007;71(12):1932–1936. doi:10.1253/circj.71.1932

43) Hoffmann M, Kleine-Weber H, Schroeder S, et al. SARS-CoV-2 Cell Entry Depends on ACE2 and TMPRSS2 and Is Blocked by a Clinically Proven Protease Inhibitor. Cell. 2020;181(2):271–280.e8. doi:10.1016/j.cell.2020.02.052.

44) Agapitov AV, Haynes WG. Role of endothelin in cardiovascular disease. J Renin Angiotensin Aldosterone Syst. 2002;3(1):1–15. doi:10.3317/jraas.2002.001.

45) Testai L, Barrese V, Soldovieri MV, et al. Expression and function of Kv7.4 channels in rat cardiac mitochondria: possible targets for cardioprotection. Cardiovasc Res. 2016;110(1):40–50. doi:10.1093/cvr/cvv281.

46) Terata K, Miura H, Liu Y, Loberiza F, Gutterman DD. Human coronary arteriolar dilation to adrenomedullin: role of nitric oxide and K(+) channels. Am J Physiol Heart Circ Physiol. 2000;279(6):H2620–H2626. doi:10.1152/ajpheart.2000.279.6.H2620.

47) Dobrzynski E, Montanari D, Agata J, Zhu J, Chao J, Chao L. Adrenomedullin improves cardiac function and prevents renal damage in streptozotocin-induced diabetic rats. Am J Physiol Endocrinol Metab. 2002;283(6):E1291–E1298. doi:10.1152/ajpendo.00147.

48) Tsuruda T, Kato J, Kitamura K, et al. Adrenomedullin: a possible autocrine or paracrine inhibitor of hypertrophy of cardiomyocytes. Hypertension. 1998;31(1 Pt 2):505–510. doi:10.1161/01.hyp.31.1.505.

49) Yerra VG, Advani A. Histones and heart failure in diabetes. Cell Mol Life Sci. 2018;75(17):3193–3213. doi:10.1007/s00018-018-2857-1.

50) Krishnan V, Chow MZ, Wang Z, et al. Histone H4 lysine 16 hypoacetylation is associated with defective DNA repair and premature senescence in Zmpste24-deficient mice. Proc Natl Acad Sci U S A. 2011;108(30):12325–12330. doi:10.1073/pnas.1102789108.

51) Hung CC, Luan J, Sims M, et al. Studies of the SIM1 gene in relation to human obesity and obesity-related traits. Int J Obes (Lond). 2007;31(3):429–434. doi:10.1038/sj.ijo.0803443.

52) Hiraki-Kajiyama T, Yamashita J, Yokoyama K, et al. Neuropeptide B mediates female sexual receptivity in medaka fish, acting in a female-specific but reversible manner. Elife. 2019;8:e39495. Published 2019 Aug 6. doi:10.7554/eLife.39495.

53) Cao L, Ge X, Gao Y, et al. Porcine epidemic diarrhea virus inhibits dsRNA-induced interferon-β production in porcine intestinal epithelial cells by blockade of the RIG-I-mediated pathway. Virol J. 2015;12:127. Published 2015 Aug 18. doi:10.1186/s12985-015-0345-x.

54) Kritas SK, Ronconi G, Caraffa A, Gallenga CE, Ross R, Conti P. Mast cells contribute to coronavirus-induced inflammation: new anti-inflammatory strategy [published online ahead of print, 2020 Feb 4]. J Biol Regul Homeost Agents. 2020;34(1):10.23812/20-Editorial-Kritas. doi:10.23812/20-Editorial-Kritas.

55) Raymond M, Ching-A-Sue G, Van Hecke O. Mast cell stabilisers, leukotriene antagonists and antihistamines: A rapid review of effectiveness in COVID-19. May 18^th^ 2020. https://www.cebm.net/covid-19/masTcell-stabilisers-leukotriene-antagonists-and-antihistamines-a-rapid-review-of-effectiveness-in-covid-19/.

56) Gao Y, Shang W, Zhang D, et al. Lnc-C/EBPβ Modulates Differentiation of MDSCs Through Downregulating IL4i1 With C/EBPβ LIP and WDR5. Front Immunol. 2019;10:1661. Published 2019 Jul 17. doi:10.3389/fimmu.2019.01661.

57) Movahedi K, Guilliams M, Van den Bossche J, et al. Identification of discrete tumor-induced myeloid-derived suppressor cell subpopulations with distinct T cell-suppressive activity. Blood. 2008;111(8):4233–4244. doi:10.1182/blood-2007-07-099226.

58) Sungnak W, Huang N, Bécavin C, et al. SARS-CoV-2 entry factors are highly expressed in nasal epithelial cells together with innate immune genes. Nat Med. 2020;26(5):681–687. doi:10.1038/s41591-020-0868-6.

59) Lukassen S, Chua RL, Trefzer T, et al. SARS-CoV-2 receptor ACE2 and TMPRSS2 are primarily expressed in bronchial transient secretory cells. EMBO J. 2020;39(10):e105114. doi:10.15252/embj.20105114.

60) Ma DY, Suthar MS. Mechanisms of innate immune evasion in re-emerging RNA viruses. Curr Opin Virol. 2015;12:26–37. doi:10.1016/j.coviro.2015.02.005.

