## Supplementary material for "Rapid host response to an infection with Coronavirus. Study of transcriptional responses with Porcine Epidemic Diarrhea Virus": material and methodes

**Supplementary file 1 of manuscript** "Rapid host response to an infection with Coronavirus. Study of transcriptional responses with Porcine Epidemic Diarrhea Virus".

### Materials and methods

*Cell culture and virus propagation.*

African green monkey kidney epithelial cells (Vero cells) were obtained from the American 109 Type Culture Collection (ATCC® CCL-81). Vero cells were cultured and maintained in Eagle's Minimum Essential Medium (EMEM) with 10% v/v fetal bovine serum (FBS; Bodinco BV, The Netherlands) and 1% v/v Antibiotic-Antimycotic mixture (anti-anti, Gibco®), 1% L-Glutamine (Gibco®), 1% non-essential amino acids (Gibco®) and 1% Sodium bicarbonate (Gibco®)(hereafter denoted as "complete medium"). PEDV strain CV777 was obtained from Pensaert and De Bouck in the late 1970s (add1, ref. 1) and induced diarrhea in experimental pigs. For preparation of PEDV virus stocks, Vero cells were grown in serum-free medium, i.e. 1:1 mixture of EMEM and UltraMCDK (Lonza Group Ltd, Switzerland) supplemented with 10 µg/ml trypsin and 1% v/v anti-anti. After each day of growth, 75% of this serum-free medium was replaced with fresh medium containing 10 µg/ml trypsin). To prepare a virus stocks for the infection experiment nearly confluent monolayers of Vero cells were infected with a multiplicity of infection of about 0.1 and grown until 75% of the cells showed cytopathogenic effect (CPE) induced by PEDV. PEDV-infected cultures were freeze-thawed twice at -70°C and the suspension was centrifuged at 4000xg for 10 min to remove cell debris. The supernatant was harvested and concentrated about 10-fold using a 30 K Amicon Ultracel centrifugal filter (Amicon, Sigma-Aldrich Chemie N.V. Zwijndrecht, the Netherlands). This concentrated high titer virus stock was used as inoculum in the infection experiment described below.
The MRV virus used was isolated in our lab from a faecal sample of diarrheic pigs diagnosed PCR-positive for the coronavirus (CoV) porcine epidemic diarrhea virus (PEDV) using Vero cells (add1, ref.2). A passage #2 virus stocks was prepared in a similar manner as described for PEDV, except that the serum-free medium was not supplemented with trypsin and was not replaced with 75% fresh medium every day of growth. Next generation sequencing (NGS) of passage #2 virus in Vero cells and blastn analysis of NGS-reads showed that more than 95% of the viral reads in this passage #2 virus stock mapped to MRV sequences posted in GenBank (add1, ref. 3). Nine of the 10 genomic RNA segments of this novel MRV were classified as serotype 3 (MRV3). The tenth segment, coding for the cell-attachment protein S1, diverged significantly from MRV3 S1 sequences posted in the databases. The deduced amino acid sequence of this S1 segment showed most homology to the S1 segment of a recently isolated MRV reassortant strain isolated from a common vole in Hungary (add1, ref. 3). A concentrate with high titer of this passage #2 virus stock was prepared in a similar manner as described above as for PEDV and used as inoculum for the infection experiment.

*Infection experiment.*

Overnight cultured Vero cells grown in plates with 2 $cm^2$ wells to nearly confluent monolayers were mock-infected, infected with the MRV3 or PEDV concentrates in quadruplicate with a multiplicity of infection of >1 for 30 min at 4°C. For PEDV and corresponding mock-infected cells, 10 µg/ml of Trypsin in serum-free medium was used to facilitate infection of Vero cells during the whole experiment. All virus and mock-infected timepoints were performed in quadruplicate. After incubation for 30 min at 4°C, virus was discarded and cells were washed twice and supplied with fresh culture medium. The cells were grown for 0, 2, 4, 6, and 16h at 37°C and 5% $CO_2$. After incubation for the indicated times, cells were placed on ice and from 3 of the 4 wells the medium was removed and 300 µl Trizol was added to these wells. Total RNA was isolated from triplicate wells using the Direct-zol™ RNA miniprep Kit (BaseClear Lab Products, Leiden, The Netherlands). RNA was eluted from the silica-based columns in 25 µl of RNAse-free water and stored at -80°C.The remaining quadruplicate well of mock-infected, infected with MRV3 or PEDV, grown for 16h, was used for staining with antibodies directed against PEDV and MRV3 (see below).

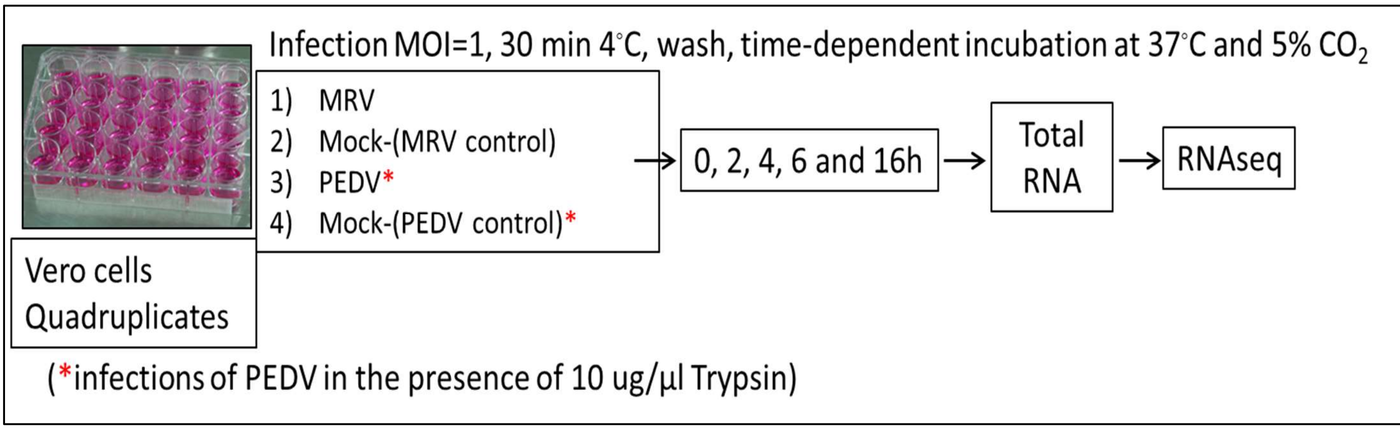


**Figure 1. add.1. Schematic presentation of the infection experiment with MRV3 and PEDV.**

*RT-PCR detection.*

The relative quantity of viral genomes of PEDV and MRV in total RNA preparations was determined by using reverse transcription (RT) -qPCR. Five µl of total RNA was used as template for reverse transcription (RT) and quantification using the Taqman FastVirus-1-step master mix for PEDV and MRV. Quantitative PCR (qPCR) tests were carried out with a LightCycler instrument (LC 480, Roche Applied Science, Mannheim, Germany) and used fluorescent labeled internal primers (probes) for detection of the amplicons. The sequence and the concentration of the forward, reverse primer and probe in PCR reactions are listed in Table add.1. In each qPCR run dilutions of virus stocks with a known titer (qPCR positive control) and culture medium of uninfected cells (qPCR negative control) were isolated and analyzed along with the samples.

**Table 1 add.1.** Primer and probe sequences used for RT-PCR detection.

| virus | Primer sequence (5'⟶ 3') | Concentration |
|---|---|---|
| PEDV | Forward: CGCAAAGACTGAACCCACTAATTT | 10 µM |
| | Reverse: TTGCCTCTGTTGTTACTTGGAGAT | 10 µM |
| | Probe : TGTTGCCATTGCCACGACTCCTGC | 10 µM |
| MRV | Forward: GGAATACTGCTGGCAAGGT | 10 µM |
| | Reverse: TCCAACGATCGGATGACGAC | 10 µM |
| | Probe : AACGGGCGAATTC | 10 µM |

*Immune peroxidase monolayer assay (IPMA).*

The medium of Vero cells, infected for 16h with PEDV, MRV3 or mock-infected was removed and cells were washed with PBS (Gibco®) and fixed in ice-cold 4% w/v paraformaldehyde (PFA) for 10 min. The PFA solution was discarded and cells were washed with PBS to remove excess PFA . Cells were incubated with a dilution of 1:200 of a mouse monoclonal antibody directed against the spike protein of PEDV or a 1:200 dilution of a rabbit polyclonal antibody raised against a peptide stretch of the attachment protein (S1) of MRV3 in PBS-Tween 20 buffer (0.05% v/v Tween 20 in PBS-Dubelco's pH 7.2) at 37 °C for 1 h. After incubation dilutions were discarded and cells were washed with PBS-Tween 20 buffer. Next, cells were incubated at 37 °C for 1h with a dilution of HRPO-conjugated rabbit anti-mouse IgG (horseradish peroxidase-conjugated Dako P-0260) of 1:200 in TBS-TB buffer (TBS-TB; 0.05% v/v Tween 20 in 20 mM Tris, 150mM NaCl pH7.2, supplemented with 1% bovine serum albumin) to detect bound anti-PEDV antibodies or with a dilution of HRP-conjugated goat anti-rabbit HRP (horseradish peroxidase-conjugated Dako P-0448) of 1:100 in TBS-TB to detect bound anti-MRV antibodies. After discarding the conjugate solutions, cells were washed with PBS-Tween 20 buffer and stained using a 1:20 dilution of 3-amino-9-ethyl-carbozole solution (4 mg/ml in DMSO; Sigma-Aldrich, The Netherlands) in 50mM sodium acetate pH 5.0 as substrate for HRP. The substrate solution was activated with 10 µl of 30% $H_2O_2$ per 20 ml.

*Preparation of RNA pools and quality analysis of RNA.*

The quantity of total RNA in all samples was determined using a NanoDrop device and equal amounts of RNA isolated from triplicate wells were pooled. The quality of each RNA pool was checked using a "High sensitivity RNA ScreenTape" in the 2200 TapeStation systems according to the manufacturer instruction. TapeStation software revisions A.02.01 was used for data analysis to calculate RIN values for each RNA pool. The RIN values for all RNA pools isolated from infected and mock-infected Vero cells at 0, 2, 4, and 6h, and of mock-infected at 16h, were > 9.0. The RIN values of PEDV and MRV infected Vero cells at 16h were <9.0. Pools of 0h, 4h and 6h were shipped on dry-ice to GenomeScan (Leiden the Netherlands) for RNAseq analysis. Details about the RNAseq analysis and data analysis and generation of data-files performed by GenomeScan are described in reports "additional file 2a and 2b". From generated datafiles we extracted differential expressed genes with a FC>2 and *p-value* of <0.05. In additional file 3, sheet "read me", details about pair-wise comparisons and number of DEGs identified in these comparisons are provided.

***References additional file 1***.

*add1, ref.1*: Pensaert, M., de Bouck, P., 1978. A new coronavirus-like particle associated with diarrhea in swine. Arch Virol. 58:243-247.

*add1, ref.2*: Marcel Hulst, Renate Hakze-van der Honing, Stephanie Vastenhouw, Alex Bossers, Frank Harders and Wim van der Poel. Identification of a Novel Reassortant of a Mammalian Orthoreovirus in Faeces of diarrheic pigs in the Netherlands: presentation at the 11th annual meeting of EPIZONE in Paris, 2017; abstract C9 in abstract book. https://www.epizone-eu.net/upload_mm/3/a/0/69d0a5dc-6eae-48dd-895a-5c2e89eb2476_EPIZONE2017%20Abstract%20book.pdf

*add1, ref.3*: Fehér, E., Kemenesi, G., Oldal, M. et al. 2017. Isolation and complete genome characterization of novel reassortant orthoreovirus from common vole (Microtus arvalis). Virus Genes 53, 307–311. https://doi.org/10.1007/s11262-016-1411-1.
