## Supplementary material for "Rapid host response to an infection with Coronavirus. Study of transcriptional responses with Porcine Epidemic Diarrhea Virus": RNAseq next generation sequencing report

### GenomeScan Report

#### Next Generation Sequencing

| | |
|---|---|
| Address: | Marcel Hulst<br>Wageningen Bioveterinary Research<br>Virology Houribweg 39<br>8221 RA Lelystad |
| Project reference: | 103618 |
| Project start date: | 2019-03-04 |
| Project manager: | David van der Meer |
| Service type: | Illumina Next Generation Sequencing |
| Progress: | Finished |
| Document status: | v1 Final |

#### Summary

This project report contains all information regarding submitted samples, qualitychecks, experimental procedures, and the resulting data that was generated in your project.

A total of 18 samples were sequenced using Illumina sequencing technology. Data analysis was performed and reported in an additional report

If you have any additional questions regarding this report, please do not hesitate to contact our Project Manager.

#### Materials

GenomeScan received in total 18 sample(s) in this batch. The samples were received in good condition on: 2019-04-03. To assess the quality of the samples, the concentration of the sample was determined using the Fragment Analyzer. Detailed quality metrics can be found in the Sample Summary table (Table 1) and the Appendix of this report. All sample(s) met our quality requirements.

Table 1. Sample information and quality metrics

| GS_ID | Customer ID | Material type | RQN | Species | Active | Entry QC (ng/ul) | Entry QC Passed |
|---|---|---|---|---|---|---|---|
| 103618-001-001 | 0-PEDV | total RNA | 9.1 | Other | Yes | 18.45 | Yes |
| 103618-001-002 | 0-PEDV control | total RNA | 9.4 | Other | Yes | 19.76 | Yes |
| 103618-001-003 | 0-MRV | total RNA | 8.3 | Other | Yes | 15.61 | Yes |
| 103618-001-004 | 0-MRV control | total RNA | 9.7 | Other | Yes | 21.77 | Yes |
| 103618-001-005 | 0-PEDV-MRV | total RNA | 9.2 | Other | Yes | 21.00 | Yes |
| 103618-001-006 | 0-PEDV-MRV control | total RNA | 9.4 | Other | Yes | 16.96 | Yes |

| GS_ID | Customer ID | Material type | RQN | Species | Active | Entry QC (ng/ul) | Entry QC Passed |
|---|---|---|---|---|---|---|---|
| 103618-001-007 | 4-PEDV | total RNA | 9.4 | Other | Yes | 15.50 | Yes |
| 103618-001-008 | 4-PEDV control | total RNA | 9.3 | Other | Yes | 24.11 | Yes |
| 103618-001-009 | 4-MRV | total RNA | 9.5 | Other | Yes | 15.46 | Yes |
| 103618-001-010 | 4-MRV control | total RNA | 9.6 | Other | Yes | 17.83 | Yes |
| 103618-001-011 | 4-PEDV-MRV | total RNA | 9.3 | Other | Yes | 18.21 | Yes |
| 103618-001-012 | 4-PEDV-MRV control | total RNA | 9.4 | Other | Yes | 22.70 | Yes |
| 103618-001-013 | 6-PEDV | total RNA | 9.3 | Other | Yes | 19.38 | Yes |
| 103618-001-014 | 6-PEDV control | total RNA | 9.4 | Other | Yes | 15.06 | Yes |
| 103618-001-015 | 6-MRV | total RNA | 9.8 | Other | Yes | 18.03 | Yes |
| 103618-001-016 | 6-MRV control | total RNA | 10.0 | Other | Yes | 14.26 | Yes |
| 103618-001-017 | 6-PEDV-MRV | total RNA | 9.6 | Other | Yes | 14.55 | Yes |
| 103618-001-018 | 6-PEDV-MRV control | total RNA | 9.4 | Other | Yes | 18.04 | Yes |

#### Experimental procedures

The NEBNext Ultra II Directional RNA Library Prep Kit for Illumina was used to process the sample(s). The sample preparation was performed according to the protocol "NEBNext Ultra II Directional RNA Library Prep Kit for Illumina" (NEB #E7760S/L). Briefly, mRNA was isolated from total RNA using the oligo-dT magnetic beads. After fragmentation of the mRNA, a cDNA synthesis was performed. This was used for ligation with the sequencing adapters and PCR amplicification of the resulting product. The quality and yield after sample preparation was measured with the Fragment Analyzer (see appendices). The size of the resulting products was consistent with the expected size distribution (a broad peak between 300-500 bp).

Table 2. Library construction

| GS_ID | Customer ID | Molarity (nM) | Index/Barcode | Prep QC Passed |
|---|---|---|---|---|
| 103618-001-001 | 0-PEDV | 2.59 | dNEB49 | Yes |
| 103618-001-002 | 0-PEDV control | 2.36 | dNEB50 | Yes |
| 103618-001-003 | 0-MRV | 1.57 | dNEB51 | Yes |
| 103618-001-004 | 0-MRV control | 2.31 | dNEB52 | Yes |
| 103618-001-005 | 0-PEDV-MRV | 4.49 | dNEB53 | Yes |
| 103618-001-006 | 0-PEDV-MRV control | 3.65 | dNEB54 | Yes |
| 103618-001-007 | 4-PEDV | 5.80 | dNEB55 | Yes |
| 103618-001-008 | 4-PEDV control | 4.25 | dNEB56 | Yes |
| 103618-001-009 | 4-MRV | 1.16 | dNEB57 | Yes |
| 103618-001-010 | 4-MRV control | 0.34 | dNEB58 | Yes |
| 103618-001-011 | 4-PEDV-MRV | 3.04 | dNEB59 | Yes |
| 103618-001-012 | 4-PEDV-MRV control | 1.83 | dNEB60 | Yes |
| 103618-001-013 | 6-PEDV | 2.75 | dNEB61 | Yes |

| GS_ID | Customer ID | Molarity (nM) | Index/Barcode | Prep QC Passed |
|---|---|---|---|---|
| 103618-001-014 | 6-PEDV control | 2.10 | dNEB62 | Yes |
| 103618-001-015 | 6-MRV | 4.69 | dNEB63 | Yes |
| 103618-001-016 | 6-MRV control | 7.48 | dNEB64 | Yes |
| 103618-001-017 | 6-PEDV-MRV | 5.57 | dNEB65 | Yes |
| 103618-001-018 | 6-PEDV-MRV control | 6.36 | dNEB66 | Yes |

Clustering and DNA sequencing using the NovaSeq6000 was performed according to manufacturer's protocols. A concentration of 1.1 nM of DNA was used. Detailed run information per flow cell can be found in Table 3. NovaSeq control software NCS v1.6 was used.

The experiments were performed at the following site(s): GenomeScan B.V., Plesmanlaan 1d, 2333 BZ, Leiden.

#### Primary data analysis and results

Image analysis, base calling, and quality check was performed with the Illumina data analysis pipeline RTA3.4.4 and Bcl2fastq v2.20. Quality-filtered sequence tags are placed in the 'Raw Data' folder on the accompanying hard disk.

The flow cell information and total raw yield (Mb) for each sample are summarised in Tables 3 and 4.

Table 3. Run info

| Flowcell ID | Device | Run length |
|---|---|---|
| HJLYHDSXX | Novaseq 6000 | 151-8-8-151 |
| HK3Y2DSXX | Novaseq 6000 | 151-8-8-151 |
| HJVT7DMXX | Novaseq 6000 | 151-8-8-151 |

Table 4. Run yields per sample

| GS_ID | Sample ID | Yield (Mb) | Clusters | % >=Q30 |
|---|---|---|---|---|
| 103618-001-001 | 0-PEDV | 6,482 | 21,465,636 | 90.97 |
| 103618-001-002 | 0-PEDV control | 5,740 | 19,006,413 | 91.15 |

| GS_ID | Sample ID | Yield (Mb) | Clusters | % >=Q30 |
|---|---|---|---|---|
| 103618-001-003 | 0-MRV | 9,324 | 30,874,087 | 90.96 |
| 103618-001-004 | 0-MRV control | 5,070 | 16,787,245 | 91.26 |
| 103618-001-005 | 0-PEDV-MRV | 6,597 | 21,844,585 | 92.77 |
| 103618-001-006 | 0-PEDV-MRV control | 6,626 | 21,939,961 | 91.65 |
| 103618-001-007 | 4-PEDV | 5,539 | 18,341,732 | 91.66 |
| 103618-001-008 | 4-PEDV control | 6,454 | 21,373,119 | 90.86 |
| 103618-001-009 | 4-MRV | 5,627 | 18,632,923 | 91.14 |
| 103618-001-011 | 4-PEDV-MRV | 5,822 | 19,275,279 | 91.86 |
| 103618-001-012 | 4-PEDV-MRV control | 6,990 | 23,147,037 | 91.93 |
| 103618-001-013 | 6-PEDV | 6,545 | 21,673,452 | 91.28 |
| 103618-001-014 | 6-PEDV control | 4,888 | 16,185,666 | 91.31 |
| 103618-001-015 | 6-MRV | 5,734 | 18,985,012 | 91.18 |
| 103618-001-016 | 6-MRV control | 5,657 | 18,734,320 | 89.89 |
| 103618-001-017 | 6-PEDV-MRV | 5,119 | 16,950,382 | 91.37 |
| 103618-001-018 | 6-PEDV-MRV control | 5,705 | 18,889,320 | 91.28 |

The raw data in the 'Raw data' directory are coded with the flow cell numbers and the per sample data can be found within these run folders.

#### Deviations, additions, exclusions from the test method

This project was performed in compliance with the requirements and specifications set by GenomeScan and the customer. All data are within specifications.

The following deviations occured:

After completion of sample preperation: user enzyme, PCR and double clean up steps were repeated. This did not work, so the whole sample preperation was repeated.

#### Data transfer and formats

The data files are stored on the accompanying data disk as part of the results delivery. The raw data can be found in the 'Raw data' folder on the disk. The report can be found in the 'Reports' folder. If data analysis is performed, the results can be found in the analysis folder.

All files provided are in either plain text (.txt), tab-delimited text (.tab, .txt), PDF format (.pdf), or OpenDocument format(.odt, .ods). Plain text files can be opened in any text editor although some files for the Next Generation Sequencing service may be too large and an editor optimised for large files is required. Tab-delimited files can be opened in any text editor or spreadsheet application. PDF files require Adobe Reader to open and OpenDocument files can be opened in Microsoft Office 2007 SP2 or later versions.

Tab delimited files (.tab) are formatted using a dot as decimal separator and without thousand separators. They should be opened using an English locale to avoid incorrect translation of decimal and thousand separators.

This project report was digitally signed by:

Lotte Salden
Technician

Stephanie van den Oever
Project Manager

Date: 2019-06-13

### *Fragment Analyzer* Run Summary:

**Filename and Data Path:** X:\Lopende Opdrachten\GAII\103618\EntryQC\Tray-1 11-11-18\2019 04 08 11H 11M.raw
**Created:** Monday, April 08, 2019 11:44:17 AM
**# of Capillaries:** 16
**Array Serial #:** 112118-01SFS
**Effect Length:** 33 cm
**Array Usage Count:** 85
**FA Version #:** 1.2.0.11
**Device Serial #:** 3003

**METHOD INFORMATION**
**Method Name:** DNF-471-33 - SS Total RNA 15nt.mthds
**Gel Prime:** No
**Full Conditioning:** Yes
**Gel Prime to Buffer:** Yes
**Gel Selection:** Gel 1
**Perform Prerun:** 8.0 kV, 30 sec.
**Rinse:** No
**Marker 1:** No
**Rinse:** Tray: 3, Row: A, # Dips: 2
**Sample Injection:** 5.0 kV, 4 sec.
**Separation:** 8.0 kV, 40.0 min.
**Tray Name:** Tray-1

**Analysis Mode:** RNA (Eukaryotic)

**NOTES**

 Copyright 2015 Advanced Analytical Technologies, Inc. 

### Gel Image

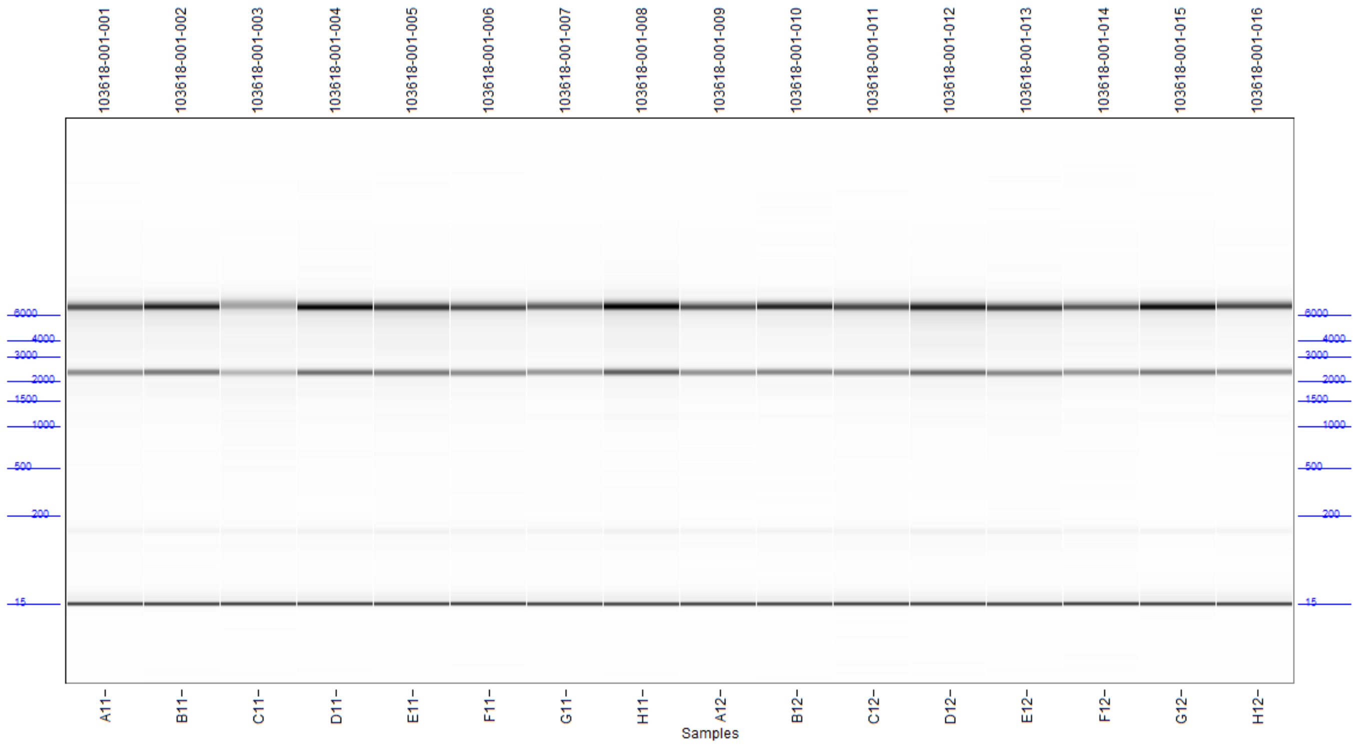


 Copyright 2015 Advanced Analytical Technologies, Inc. 

**Filename and Data Path:** X:\Lopende Opdrachten\GAII\103618\EntryQC\Tray-1 11-11-18\2019 04 08 11H 11M.raw

A11: 103618-001-001

RQN: 9.5

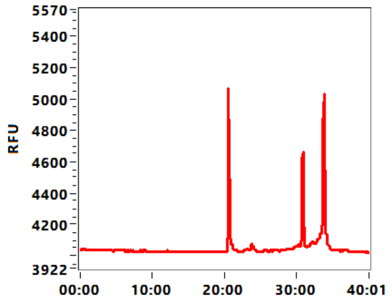


B11: 103618-001-002

RQN: 9.8

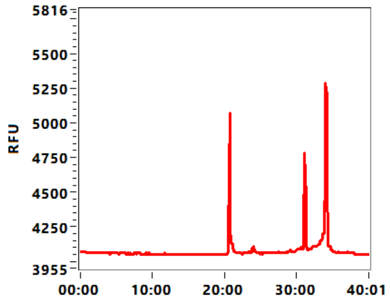


C11: 103618-001-003

RQN: 8.8

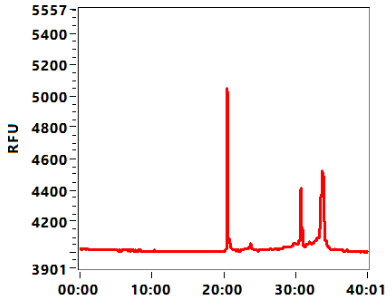


D11: 103618-001-004

RQN: 10.0

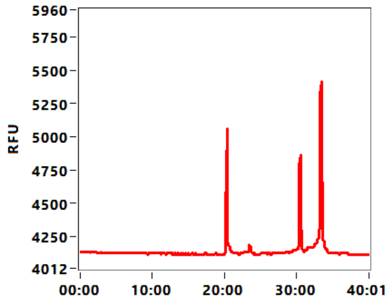


E11: 103618-001-005

RQN: 9.8

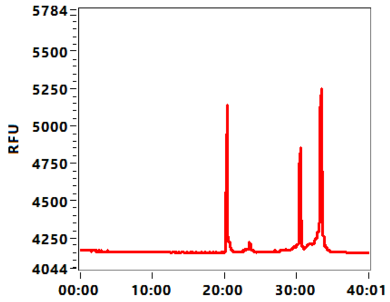


F11: 103618-001-006

RQN: 9.8

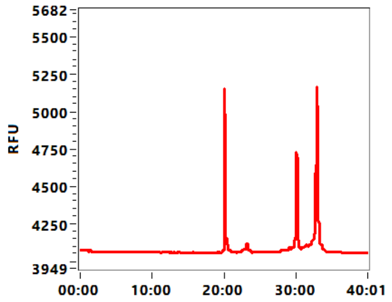


G11: 103618-001-007

RQN: 9.8

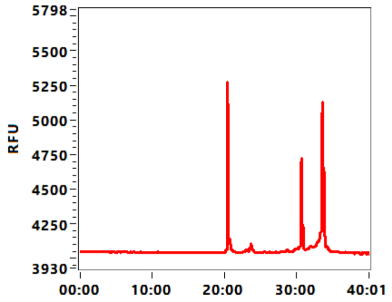


H11: 103618-001-008

RQN: 9.7

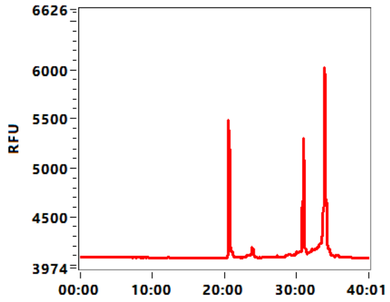


A12: 103618-001-009

RQN: 10.0

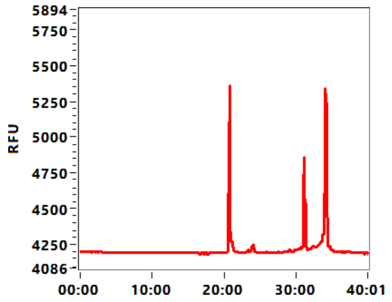


B12: 103618-001-010

RQN: 10.0

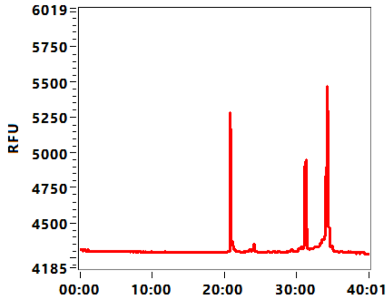


C12: 103618-001-011

RQN: 9.7

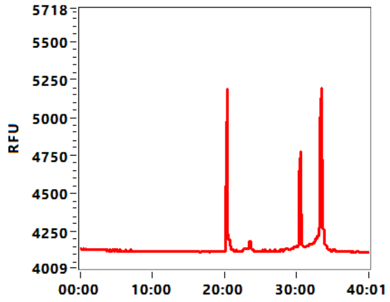


D12: 103618-001-012

RQN: 9.7

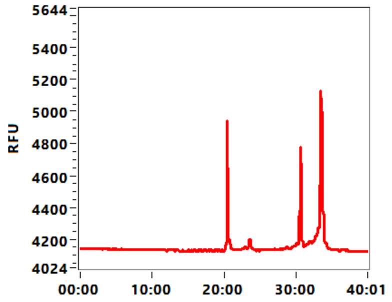


 Copyright 2015 Advanced Analytical Technologies, Inc. 

**Filename and Data Path:** X:\Lopende Opdrachten\GAII\103618\EntryQC\Tray-1 11-11-18\2019 04 08 11H 11M.raw

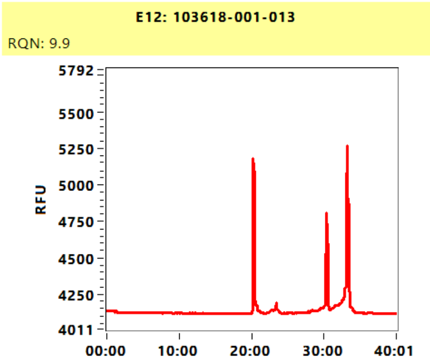


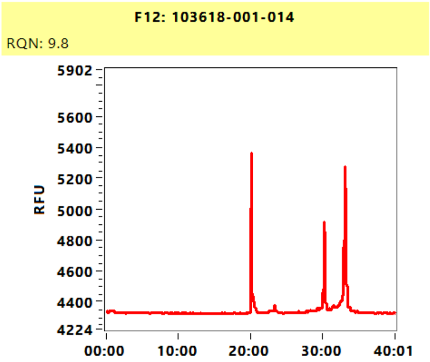


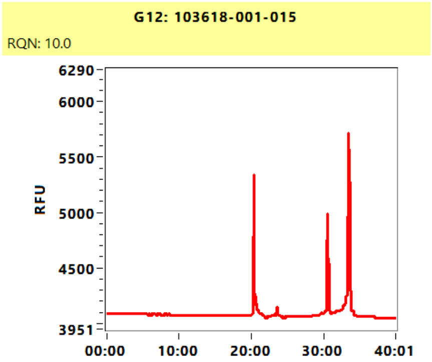


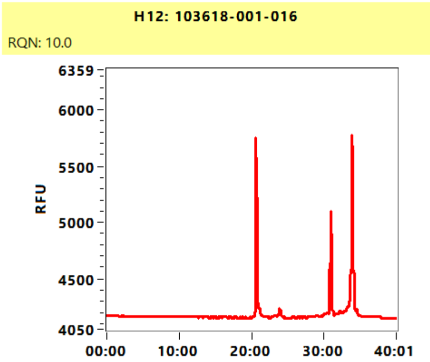


 Copyright 2015 Advanced Analytical Technologies, Inc. 

**Sample:** 103618-001-001
**Well Location:** A11
**Created:** Monday, April 08, 2019 11:44:17 AM

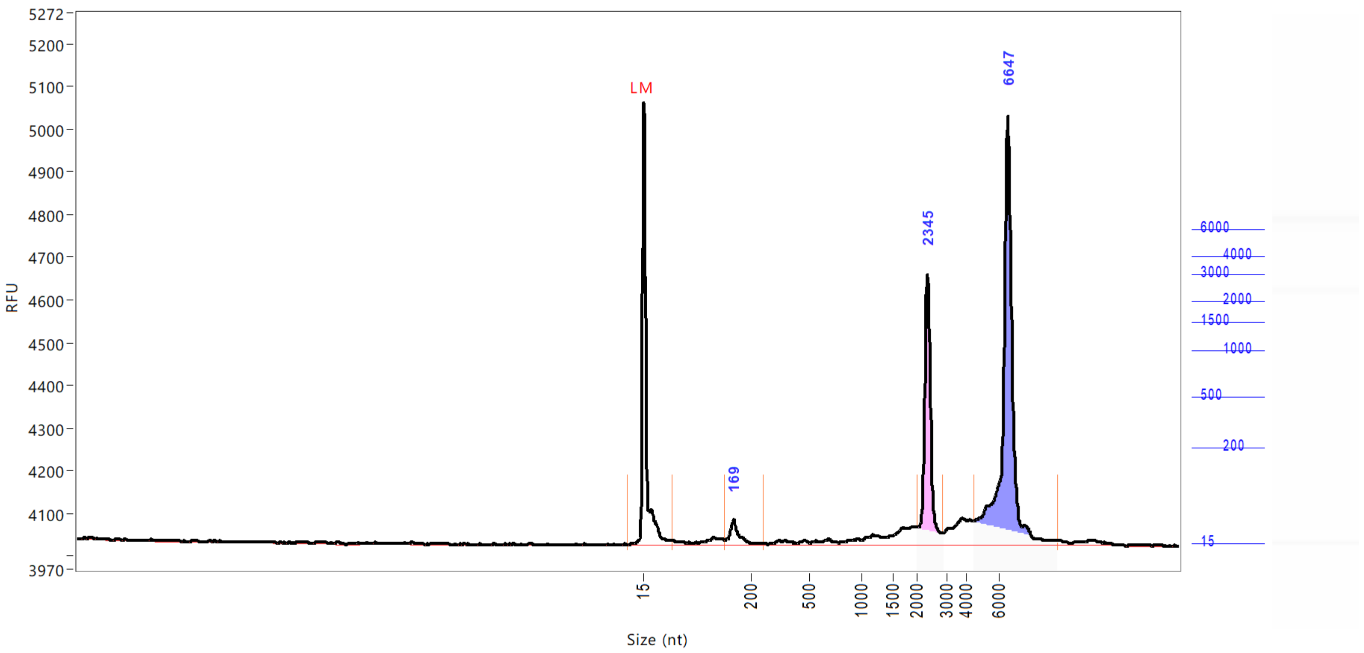


| Peak | Size (nt) | Conc. (ng/uL) | From (nt) | To (nt) | RFU |
|---|---|---|---|---|---|
| 1 | 15 (LM) | 0.5016 | 0 | 65 | 1036 |
| 2 | 169 | 0.6957 | 155 | 261 | 58 |
| 3 | 2345 | 4.2583 | 2000 | 2869 | 632 |
| 4 | 6647 | 9.5599 | 4470 | 9792 | 1003 |

| | | |
|---|---|---|
| TIC: | 14.5139 | ng/uL |
| TIM: | 22.6038 | nmole/L |
| Total Conc.: | 18.4465 | ng/uL |
| 28S/18S: | 2.2 | |
| RQN | 9.5 | |

Sample Peak Width (sec): 6 Sample Min Peak Height: 50 Sample Baseline V to V?: Y Sample Baseline V to V pts: 3
Sample Filter: Binomial # of Pts for Filter: 9 Sample Start Region (min): 0 Sample End Region (min): 40
Manual Baseline Start (min): 18 Manual Baseline End (min): 38
Marker Peak Width (sec): 6 Marker Min Peak Height: 100 Marker Baseline V to V?: Y Marker Baseline V to V pts: 3
Lower Marker Selection: First Peak > 100 RFU Upper Marker Selection: Last Peak > 100 RFU
Ladder Size (nt): 15, 200, 500, 1000, 1500, 2000, 3000, 4000, 6000
Quantification Using: Ladder Final Concentration (ng/uL): 8.0000 Dilution Factor: 12.0
Min. RFU for Data Processing: 2

 Copyright 2015 Advanced Analytical Technologies, Inc. 

**Sample:** 103618-001-002
**Well Location:** B11
**Created:** Monday, April 08, 2019 11:44:17 AM

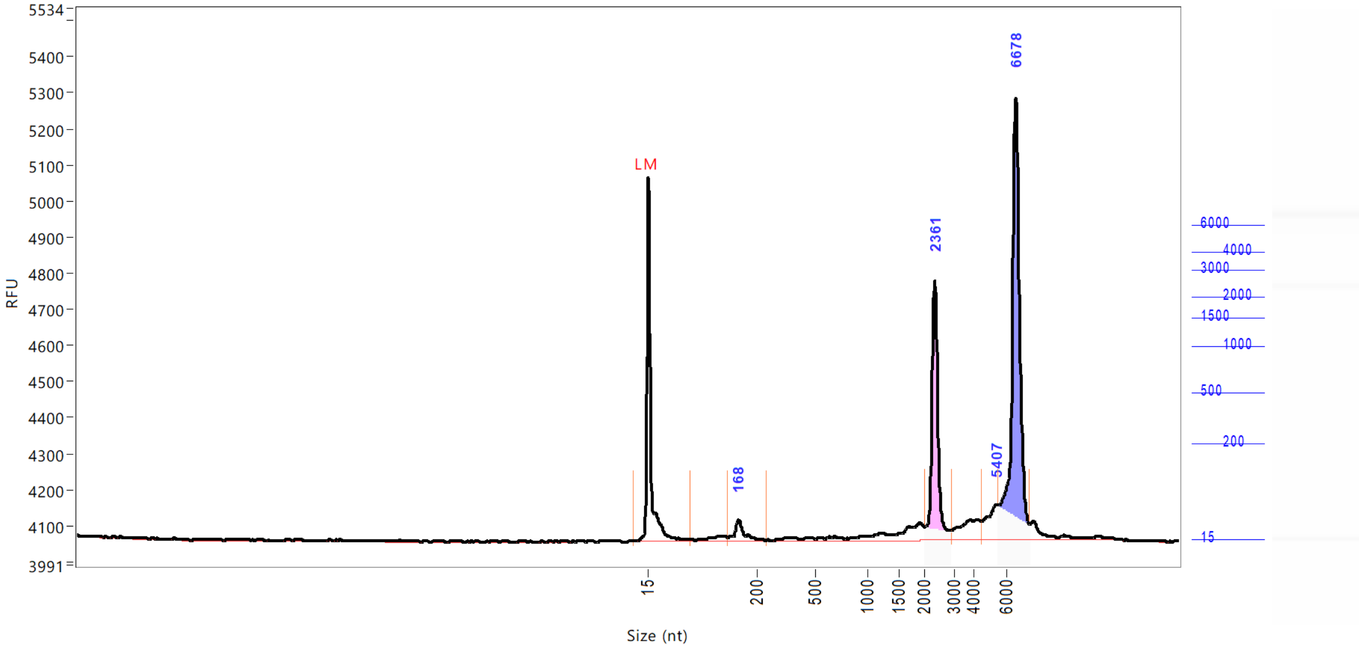


| Peak | Size (nt) | Conc. (ng/uL) | From (nt) | To (nt) | RFU |
|---|---|---|---|---|---|
| 1 | 15 (LM) | 0.5016 | 0 | 88 | 1006 |
| 2 | 168 | 0.6504 | 151 | 249 | 57 |
| 3 | 2361 | 4.7299 | 2017 | 2902 | 715 |
| 4 | 5407 | 1.0557 | 4501 | 5563 | 97 |
| 5 | 6678 | 9.6578 | 5563 | 7572 | 1225 |

| | | |
|---|---|---|
| TIC: | 16.0938 | ng/uL |
| TIM: | 23.2008 | nmole/L |
| Total Conc.: | 19.7623 | ng/uL |
| 28S/18S: | 2.0 | |
| RQN | 9.8 | |

Sample Peak Width (sec): 6 Sample Min Peak Height: 50 Sample Baseline V to V?: Y Sample Baseline V to V pts: 3
Sample Filter: Binomial # of Pts for Filter: 9 Sample Start Region (min): 0 Sample End Region (min): 40
Manual Baseline Start (min): 18 Manual Baseline End (min): 38
Marker Peak Width (sec): 6 Marker Min Peak Height: 100 Marker Baseline V to V?: Y Marker Baseline V to V pts: 3
Lower Marker Selection: First Peak > 100 RFU Upper Marker Selection: Last Peak > 100 RFU
Ladder Size (nt): 15, 200, 500, 1000, 1500, 2000, 3000, 4000, 6000
Quantification Using: Ladder Final Concentration (ng/uL): 8.0000 Dilution Factor: 12.0
Min. RFU for Data Processing: 2

 Copyright 2015 Advanced Analytical Technologies, Inc. 

**Sample:** 103618-001-003
**Well Location:** C11
**Created:** Monday, April 08, 2019 11:44:17 AM

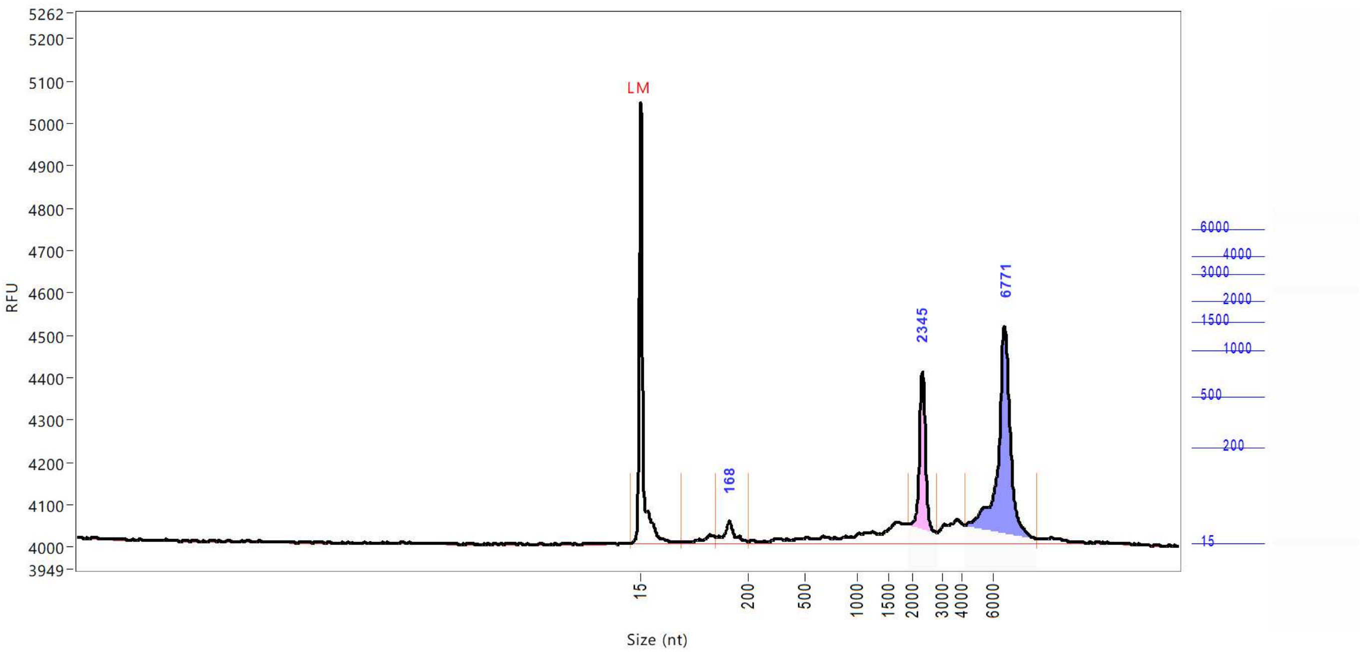


| Peak | Size (nt) | Conc. (ng/uL) | From (nt) | To (nt) | RFU |
|---|---|---|---|---|---|
| 1 | 15 (LM) | 0.5016 | 0 | 87 | 1043 |
| 2 | 168 | 0.7539 | 146 | 204 | 54 |
| 3 | 2345 | 3.2399 | 1920 | 2836 | 406 |
| 4 | 6771 | 7.2699 | 4188 | 8806 | 512 |

| | | |
|---|---|---|
| TIC: | 11.2637 | ng/uL |
| TIM: | 21.7297 | nmole/L |
| Total Conc.: | 15.6052 | ng/uL |
| 28S/18S: | 2.4 | |
| RQN | 8.8 | |

Sample Peak Width (sec): 6    Sample Min Peak Height: 50    Sample Baseline V to V?: Y    Sample Baseline V to V pts: 3
Sample Filter: Binomial    # of Pts for Filter: 9    Sample Start Region (min): 0    Sample End Region (min): 40
Manual Baseline Start (min): 18    Manual Baseline End (min): 38
Marker Peak Width (sec): 6    Marker Min Peak Height: 100    Marker Baseline V to V?: Y    Marker Baseline V to V pts: 3
Lower Marker Selection: First Peak > 100 RFU    Upper Marker Selection: Last Peak > 100 RFU
Ladder Size (nt): 15, 200, 500, 1000, 1500, 2000, 3000, 4000, 6000
Quantification Using: Ladder    Final Concentration (ng/uL): 8.0000    Dilution Factor: 12.0
Min. RFU for Data Processing: 2

 Copyright 2015 Advanced Analytical Technologies, Inc. 

**Sample:** 103618-001-004
**Well Location:** D11
**Created:** Monday, April 08, 2019 11:44:17 AM

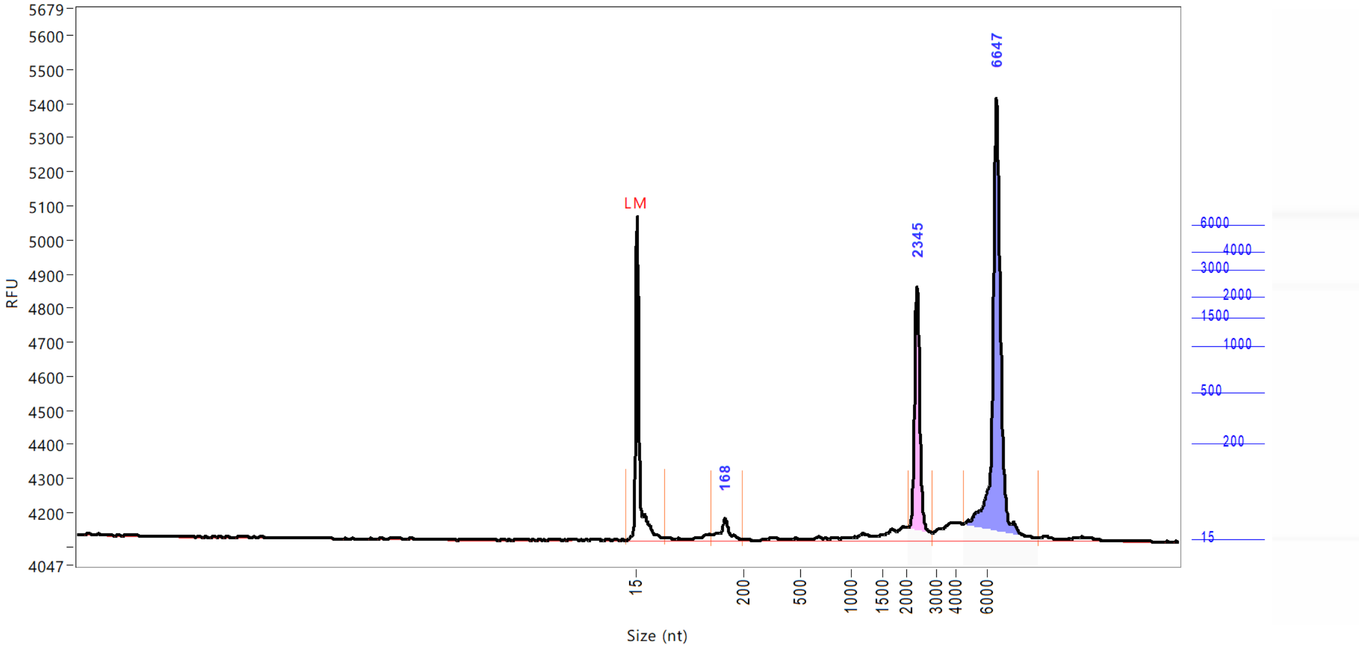


| Peak | Size (nt) | Conc. (ng/uL) | From (nt) | To (nt) | RFU |
|---|---|---|---|---|---|
| 1 | 15 (LM) | 0.5016 | 0 | 64 | 948 |
| 2 | 168 | 0.8390 | 145 | 199 | 64 |
| 3 | 2345 | 5.1327 | 2017 | 2869 | 742 |
| 4 | 6647 | 12.2906 | 4470 | 9330 | 1300 |

| | | |
|---|---|---|
| TIC: | 18.2623 | ng/uL |
| TIM: | 28.3087 | nmole/L |
| Total Conc.: | 21.7684 | ng/uL |
| 28S/18S: | 2.4 | |
| RQN | 10.0 | |

Sample Peak Width (sec): 6 Sample Min Peak Height: 50 Sample Baseline V to V?: Y Sample Baseline V to V pts: 3
Sample Filter: Binomial # of Pts for Filter: 9 Sample Start Region (min): 0 Sample End Region (min): 40
Manual Baseline Start (min): 18 Manual Baseline End (min): 38
Marker Peak Width (sec): 6 Marker Min Peak Height: 100 Marker Baseline V to V?: Y Marker Baseline V to V pts: 3
Lower Marker Selection: First Peak > 100 RFU Upper Marker Selection: Last Peak > 100 RFU
Ladder Size (nt): 15, 200, 500, 1000, 1500, 2000, 3000, 4000, 6000
Quantification Using: Ladder Final Concentration (ng/uL): 8.0000 Dilution Factor: 12.0
Min. RFU for Data Processing: 2

 Copyright 2015 Advanced Analytical Technologies, Inc. 

**Sample:** 103618-001-005
**Well Location:** E11
**Created:** Monday, April 08, 2019 11:44:17 AM

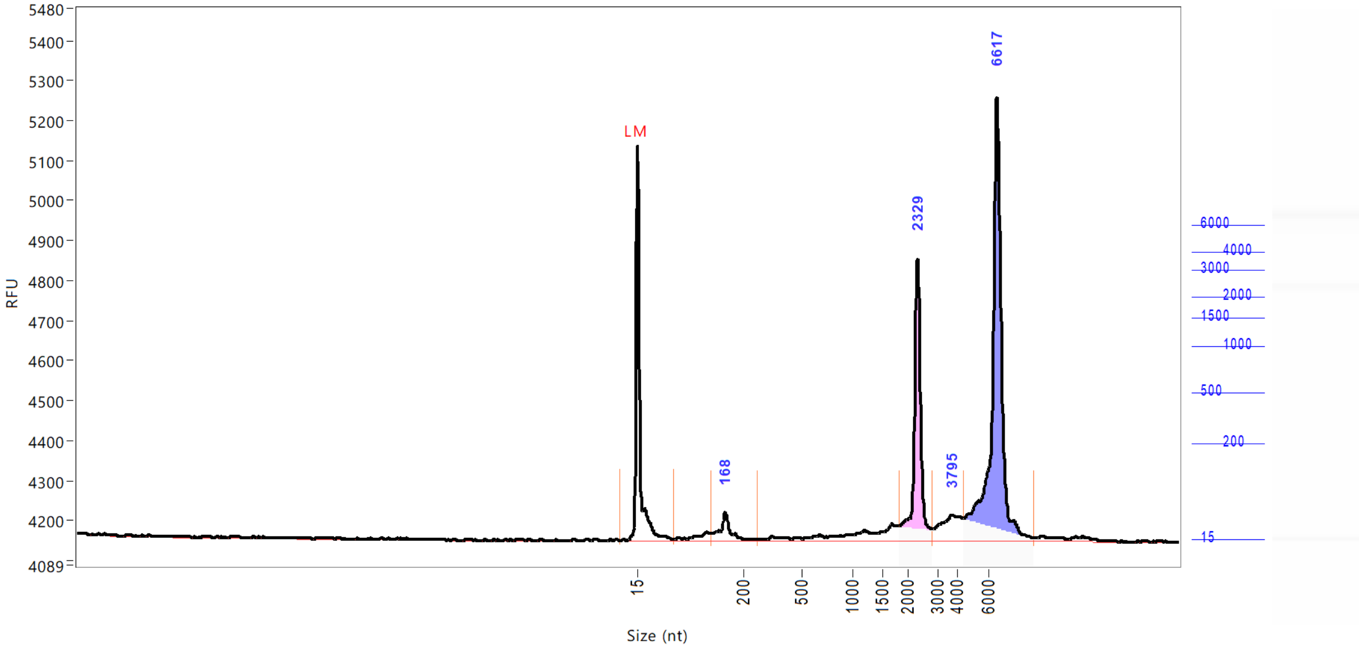


| Peak | Size (nt) | Conc. (ng/uL) | From (nt) | To (nt) | RFU |
|---|---|---|---|---|---|
| 1 | 15 (LM) | 0.5016 | 0 | 78 | 985 |
| 2 | 168 | 0.9009 | 143 | 266 | 68 |
| 3 | 2329 | 5.2340 | 1830 | 2836 | 701 |
| 4 | 3795 | 1.4328 | 2836 | 4438 | 64 |
| 5 | 6617 | 11.0159 | 4438 | 8991 | 1108 |

| | | |
|---|---|---|
| TIC: | 18.5836 | ng/uL |
| TIM: | 30.4039 | nmole/L |
| Total Conc.: | 20.9953 | ng/uL |
| 28S/18S: | 2.1 | |
| RQN | 9.8 | |

Sample Peak Width (sec): 6 Sample Min Peak Height: 50 Sample Baseline V to V?: Y Sample Baseline V to V pts: 3
Sample Filter: Binomial # of Pts for Filter: 9 Sample Start Region (min): 0 Sample End Region (min): 40
Manual Baseline Start (min): 18 Manual Baseline End (min): 38
Marker Peak Width (sec): 6 Marker Min Peak Height: 100 Marker Baseline V to V?: Y Marker Baseline V to V pts: 3
Lower Marker Selection: First Peak > 100 RFU Upper Marker Selection: Last Peak > 100 RFU
Ladder Size (nt): 15, 200, 500, 1000, 1500, 2000, 3000, 4000, 6000
Quantification Using: Ladder Final Concentration (ng/uL): 8.0000 Dilution Factor: 12.0
Min. RFU for Data Processing: 2


Copyright 2015 Advanced Analytical Technologies, Inc.

**Sample:** 103618-001-006
**Well Location:** F11
**Created:** Monday, April 08, 2019 11:44:17 AM

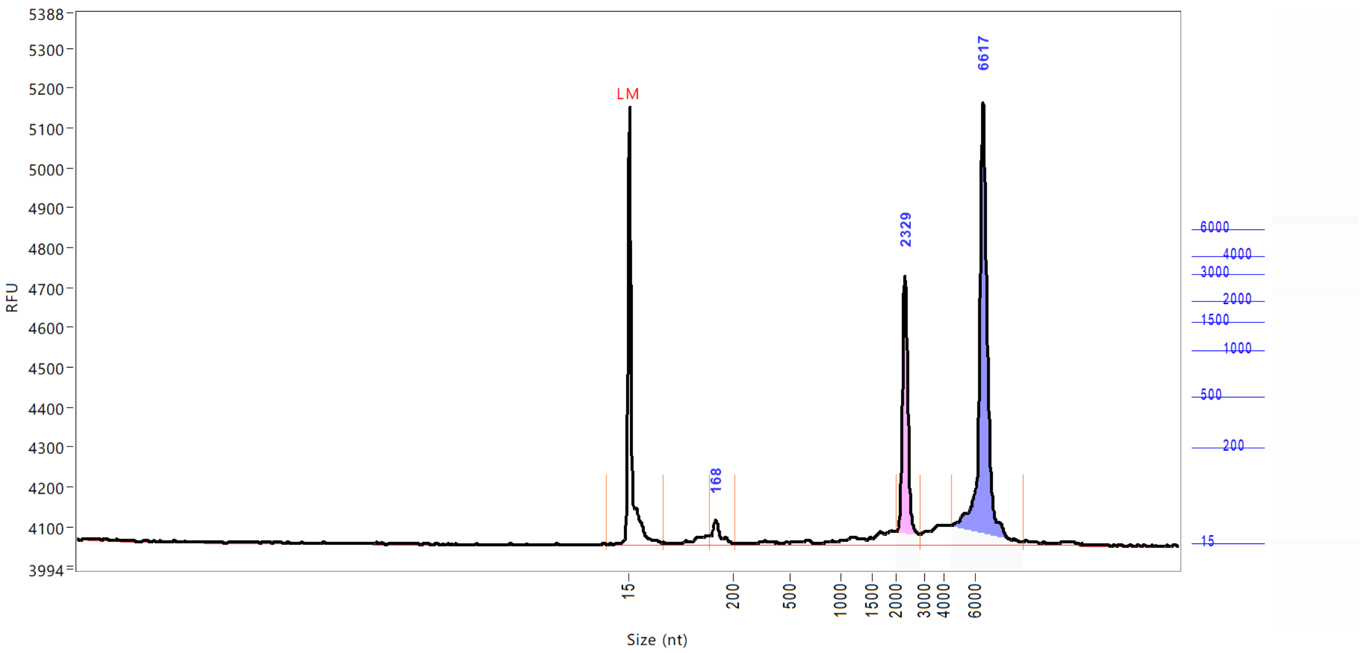


| Peak | Size (nt) | Conc. (ng/uL) | From (nt) | To (nt) | RFU |
|---|---|---|---|---|---|
| 1 | 15 (LM) | 0.5016 | 0 | 75 | 1095 |
| 2 | 168 | 0.5949 | 158 | 209 | 61 |
| 3 | 2329 | 4.0178 | 2017 | 2853 | 675 |
| 4 | 6617 | 9.1487 | 4501 | 9206 | 1111 |

| | | |
|---|---|---|
| TIC: | 13.7614 | ng/uL |
| TIM: | 20.5558 | nmole/L |
| Total Conc.: | 16.9574 | ng/uL |
| 28S/18S: | 2.2 | |
| RQN | 9.8 | |

Sample Peak Width (sec): 6 Sample Min Peak Height: 50 Sample Baseline V to V?: Y Sample Baseline V to V pts: 3
Sample Filter: Binomial # of Pts for Filter: 9 Sample Start Region (min): 0 Sample End Region (min): 40
Manual Baseline Start (min): 18 Manual Baseline End (min): 38
Marker Peak Width (sec): 6 Marker Min Peak Height: 100 Marker Baseline V to V?: Y Marker Baseline V to V pts: 3
Lower Marker Selection: First Peak > 100 RFU Upper Marker Selection: Last Peak > 100 RFU
Ladder Size (nt): 15, 200, 500, 1000, 1500, 2000, 3000, 4000, 6000
Quantification Using: Ladder Final Concentration (ng/uL): 8.0000 Dilution Factor: 12.0
Min. RFU for Data Processing: 2

 Copyright 2015 Advanced Analytical Technologies, Inc. 

**Sample:** 103618-001-007
**Well Location:** G11
**Created:** Monday, April 08, 2019 11:44:17 AM

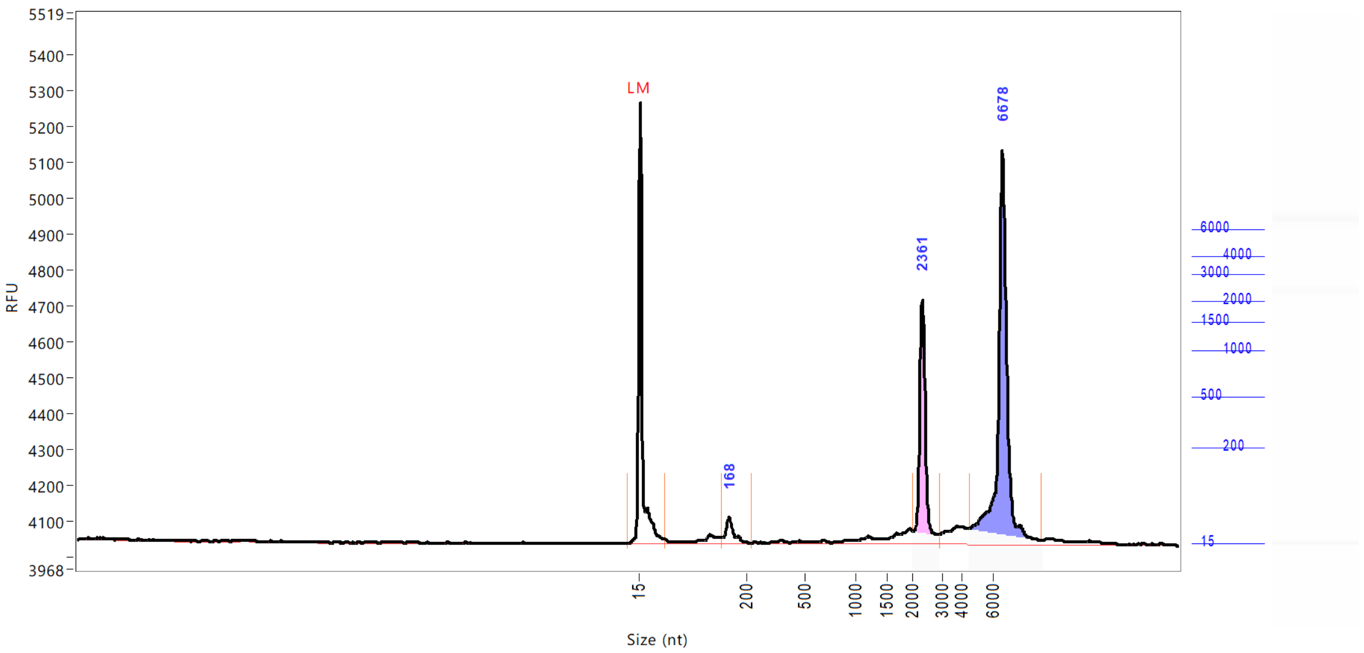


| Peak | Size (nt) | Conc. (ng/uL) | From (nt) | To (nt) | RFU |
|---|---|---|---|---|---|
| 1 | 15 (LM) | 0.5016 | 0 | 58 | 1231 |
| 2 | 168 | 0.6575 | 158 | 219 | 73 |
| 3 | 2361 | 3.6895 | 2034 | 2918 | 678 |
| 4 | 6678 | 8.3078 | 4501 | 9206 | 1095 |

| | | |
|---|---|---|
| TIC: | 12.6549 | ng/uL |
| TIM: | 20.6882 | nmole/L |
| Total Conc.: | 15.4973 | ng/uL |
| 28S/18S: | 2.3 | |
| RQN | 9.8 | |

Sample Peak Width (sec): 6 Sample Min Peak Height: 50 Sample Baseline V to V?: Y Sample Baseline V to V pts: 3
Sample Filter: Binomial # of Pts for Filter: 9 Sample Start Region (min): 0 Sample End Region (min): 40
Manual Baseline Start (min): 18 Manual Baseline End (min): 38
Marker Peak Width (sec): 6 Marker Min Peak Height: 100 Marker Baseline V to V?: Y Marker Baseline V to V pts: 3
Lower Marker Selection: First Peak > 100 RFU Upper Marker Selection: Last Peak > 100 RFU
Ladder Size (nt): 15, 200, 500, 1000, 1500, 2000, 3000, 4000, 6000
Quantification Using: Ladder Final Concentration (ng/uL): 8.0000 Dilution Factor: 12.0
Min. RFU for Data Processing: 2


Copyright 2015 Advanced Analytical Technologies, Inc.

**Sample:** 103618-001-008
**Well Location:** H11
**Created:** Monday, April 08, 2019 11:44:17 AM

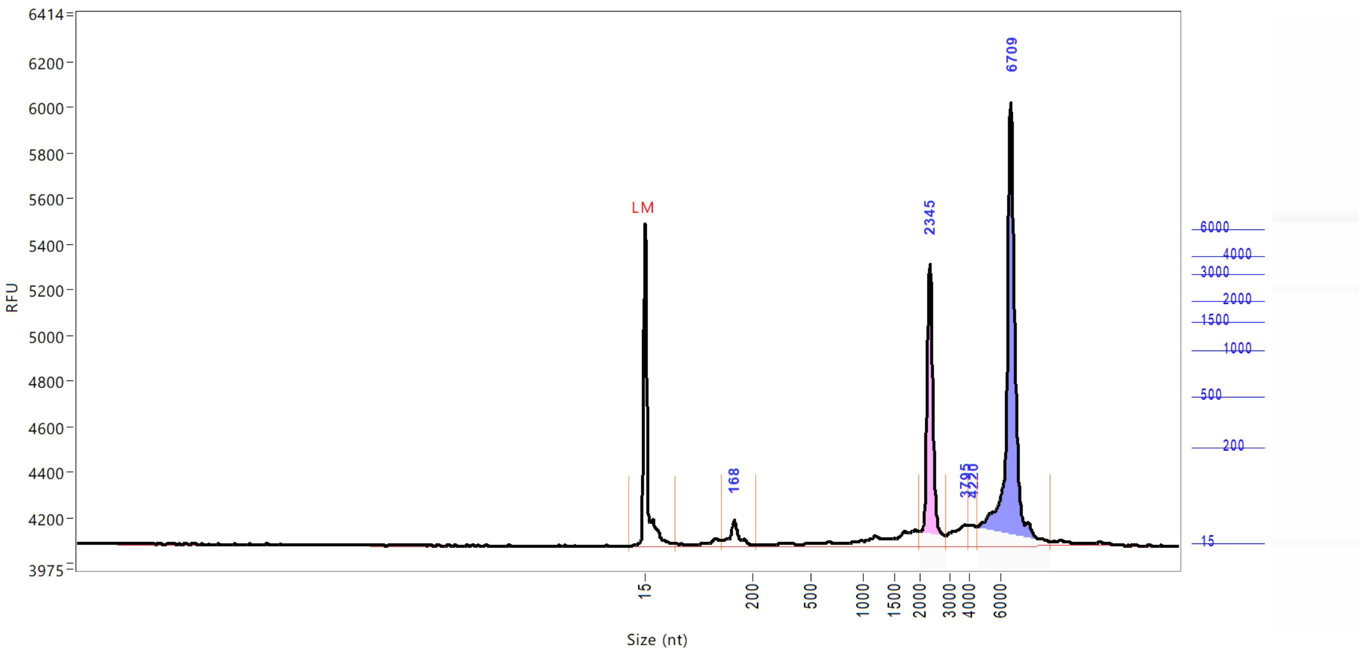


| Peak | Size (nt) | Conc. (ng/uL) | From (nt) | To (nt) | RFU |
|---|---|---|---|---|---|
| 1 | 15 (LM) | 0.5016 | 0 | 66 | 1412 |
| 2 | 168 | 0.9417 | 146 | 216 | 110 |
| 3 | 2345 | 5.9644 | 2001 | 2902 | 1234 |
| 4 | 3795 | 0.9869 | 2902 | 4000 | 90 |
| 5 | 4220 | 0.5119 | 4000 | 4563 | 88 |
| 6 | 6709 | 12.8279 | 4563 | 9206 | 1946 |

| | | |
|---|---|---|
| TIC: | 21.2328 | ng/uL |
| TIM: | 32.6292 | nmole/L |
| Total Conc.: | 24.1070 | ng/uL |
| 28S/18S: | 2.1 | |
| RQN | 9.7 | |

Sample Peak Width (sec): 6 Sample Min Peak Height: 50 Sample Baseline V to V?: Y Sample Baseline V to V pts: 3
Sample Filter: Binomial # of Pts for Filter: 9 Sample Start Region (min): 0 Sample End Region (min): 40
Manual Baseline Start (min): 18 Manual Baseline End (min): 38
Marker Peak Width (sec): 6 Marker Min Peak Height: 100 Marker Baseline V to V?: Y Marker Baseline V to V pts: 3
Lower Marker Selection: First Peak > 100 RFU Upper Marker Selection: Last Peak > 100 RFU
Ladder Size (nt): 15, 200, 500, 1000, 1500, 2000, 3000, 4000, 6000
Quantification Using: Ladder Final Concentration (ng/uL): 8.0000 Dilution Factor: 12.0
Min. RFU for Data Processing: 2

 Copyright 2015 Advanced Analytical Technologies, Inc. 

**Sample:** 103618-001-009
**Well Location:** A12
**Created:** Monday, April 08, 2019 11:44:17 AM

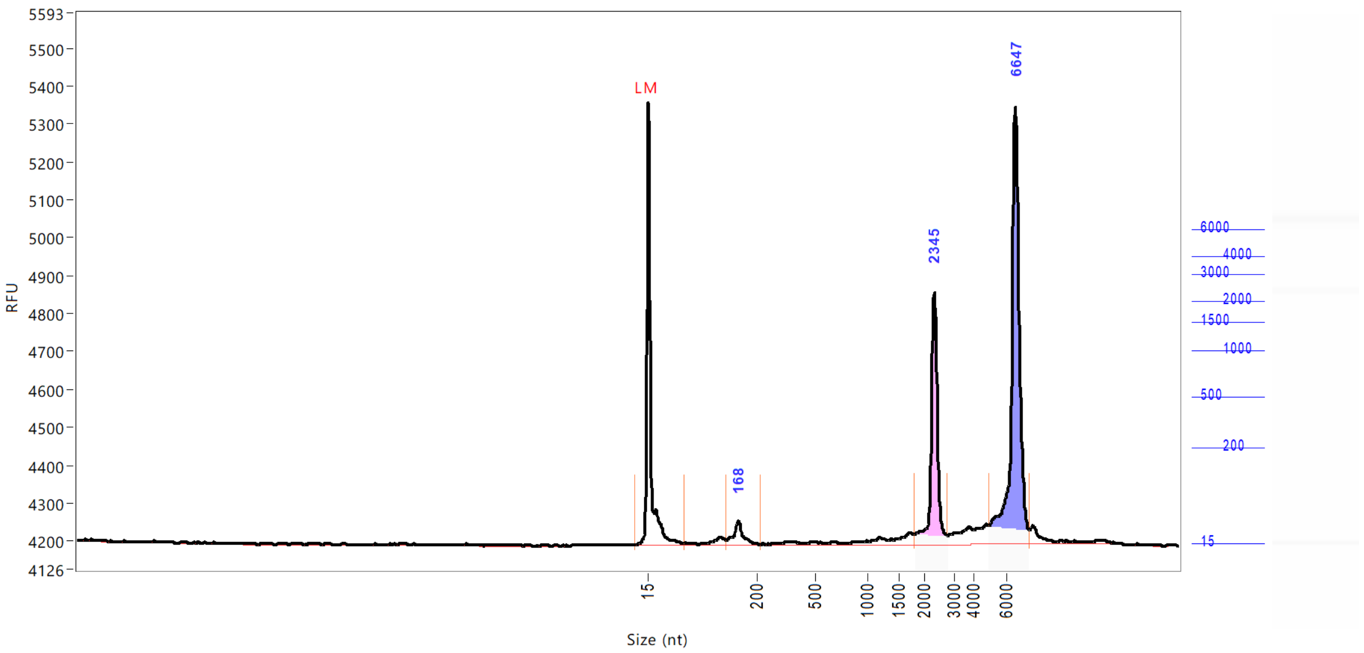


| Peak | Size (nt) | Conc. (ng/uL) | From (nt) | To (nt) | RFU |
|---|---|---|---|---|---|
| 1 | 15 (LM) | 0.5016 | 0 | 76 | 1167 |
| 2 | 168 | 0.6359 | 148 | 216 | 60 |
| 3 | 2345 | 3.9802 | 1820 | 2787 | 664 |
| 4 | 6647 | 8.1234 | 4969 | 7542 | 1151 |

| | | |
|---|---|---|
| TIC: | 12.7396 | ng/uL |
| TIM: | 21.0096 | nmole/L |
| Total Conc.: | 15.4606 | ng/uL |
| 28S/18S: | 2.1 | |
| RQN | 10.0 | |

Sample Peak Width (sec): 6    Sample Min Peak Height: 50    Sample Baseline V to V?: Y    Sample Baseline V to V pts: 3
Sample Filter: Binomial    # of Pts for Filter: 9    Sample Start Region (min): 0    Sample End Region (min): 40
Manual Baseline Start (min): 18    Manual Baseline End (min): 38
Marker Peak Width (sec): 6    Marker Min Peak Height: 100    Marker Baseline V to V?: Y    Marker Baseline V to V pts: 3
Lower Marker Selection: First Peak > 100 RFU    Upper Marker Selection: Last Peak > 100 RFU
Ladder Size (nt): 15, 200, 500, 1000, 1500, 2000, 3000, 4000, 6000
Quantification Using: Ladder    Final Concentration (ng/uL): 8.0000    Dilution Factor: 12.0
Min. RFU for Data Processing: 2


Copyright 2015 Advanced Analytical Technologies, Inc.

**Sample:** 103618-001-010
**Well Location:** B12
**Created:** Monday, April 08, 2019 11:44:17 AM

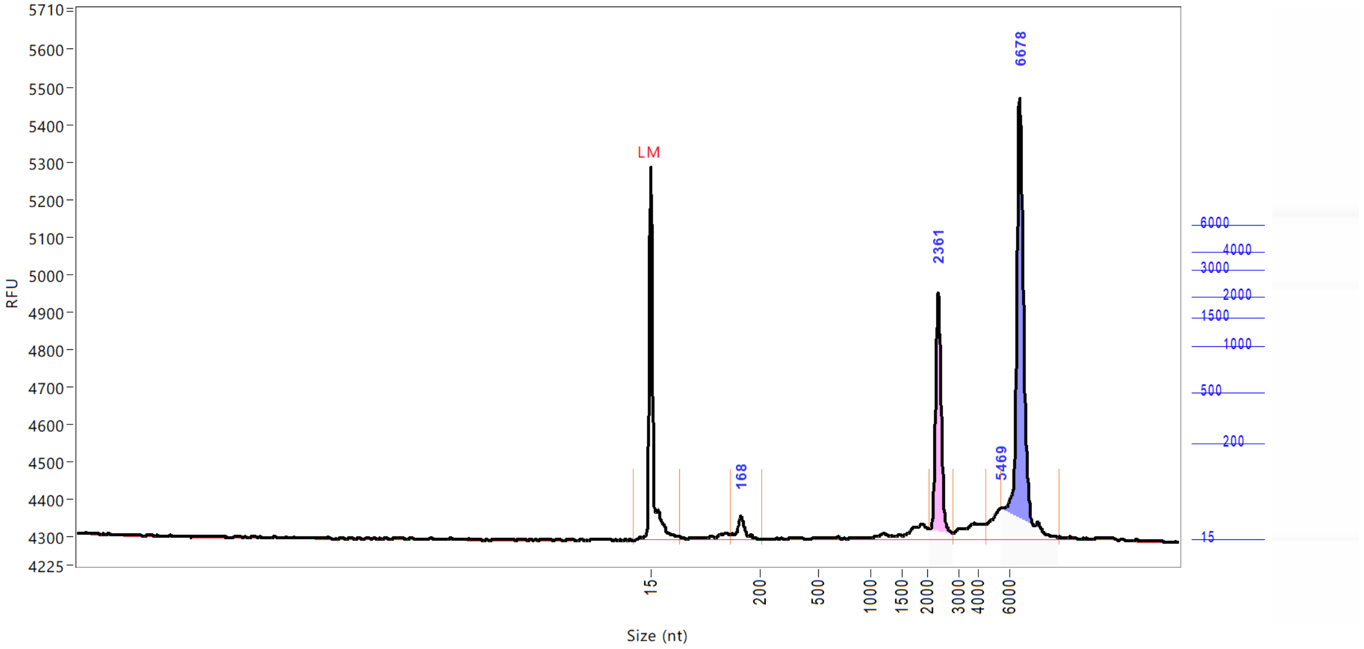


| Peak | Size (nt) | Conc. (ng/uL) | From (nt) | To (nt) | RFU |
|---|---|---|---|---|---|
| 1 | 15 (LM) | 0.5016 | 0 | 63 | 993 |
| 2 | 168 | 0.6599 | 150 | 214 | 61 |
| 3 | 2361 | 4.3038 | 2067 | 2853 | 659 |
| 4 | 5469 | 0.8221 | 4501 | 5500 | 82 |
| 5 | 6678 | 9.5387 | 5500 | 9145 | 1178 |

| | | |
|---|---|---|
| TIC: | 15.3245 | ng/uL |
| TIM: | 22.8408 | nmole/L |
| Total Conc.: | 17.8316 | ng/uL |
| 28S/18S: | 1.9 | |
| RQN | 10.0 | |

Sample Peak Width (sec): 6 Sample Min Peak Height: 50 Sample Baseline V to V?: Y Sample Baseline V to V pts: 3
Sample Filter: Binomial # of Pts for Filter: 9 Sample Start Region (min): 0 Sample End Region (min): 40
Manual Baseline Start (min): 18 Manual Baseline End (min): 38
Marker Peak Width (sec): 6 Marker Min Peak Height: 100 Marker Baseline V to V?: Y Marker Baseline V to V pts: 3
Lower Marker Selection: First Peak > 100 RFU Upper Marker Selection: Last Peak > 100 RFU
Ladder Size (nt): 15, 200, 500, 1000, 1500, 2000, 3000, 4000, 6000
Quantification Using: Ladder Final Concentration (ng/uL): 8.0000 Dilution Factor: 12.0
Min. RFU for Data Processing: 2

 Copyright 2015 Advanced Analytical Technologies, Inc. 

**Sample:** 103618-001-011
**Well Location:** C12
**Created:** Monday, April 08, 2019 11:44:17 AM

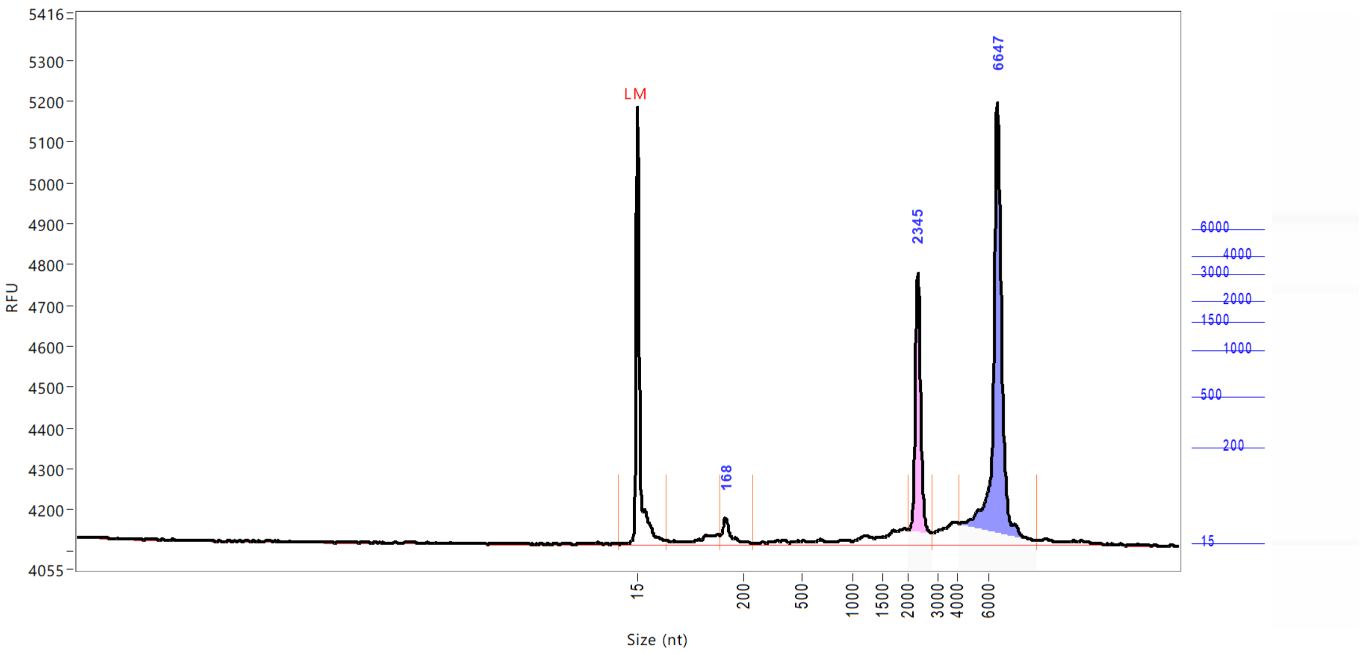


| Peak | Size (nt) | Conc. (ng/uL) | From (nt) | To (nt) | RFU |
|---|---|---|---|---|---|
| 1 | 15 (LM) | 0.5016 | 0 | 65 | 1067 |
| 2 | 168 | 0.6936 | 159 | 249 | 64 |
| 3 | 2345 | 4.1821 | 2001 | 2836 | 663 |
| 4 | 6647 | 9.7939 | 4157 | 9145 | 1082 |

| | | |
|---|---|---|
| TIC: | 14.6697 | ng/uL |
| TIM: | 22.6592 | nmole/L |
| Total Conc.: | 18.2083 | ng/uL |
| 28S/18S: | 2.3 | |
| RQN | 9.7 | |

Sample Peak Width (sec): 6 Sample Min Peak Height: 50 Sample Baseline V to V?: Y Sample Baseline V to V pts: 3
Sample Filter: Binomial # of Pts for Filter: 9 Sample Start Region (min): 0 Sample End Region (min): 40
Manual Baseline Start (min): 18 Manual Baseline End (min): 38
Marker Peak Width (sec): 6 Marker Min Peak Height: 100 Marker Baseline V to V?: Y Marker Baseline V to V pts: 3
Lower Marker Selection: First Peak > 100 RFU Upper Marker Selection: Last Peak > 100 RFU
Ladder Size (nt): 15, 200, 500, 1000, 1500, 2000, 3000, 4000, 6000
Quantification Using: Ladder Final Concentration (ng/uL): 8.0000 Dilution Factor: 12.0
Min. RFU for Data Processing: 2

 Copyright 2015 Advanced Analytical Technologies, Inc. 

**Sample:** 103618-001-012
**Well Location:** D12
**Created:** Monday, April 08, 2019 11:44:17 AM

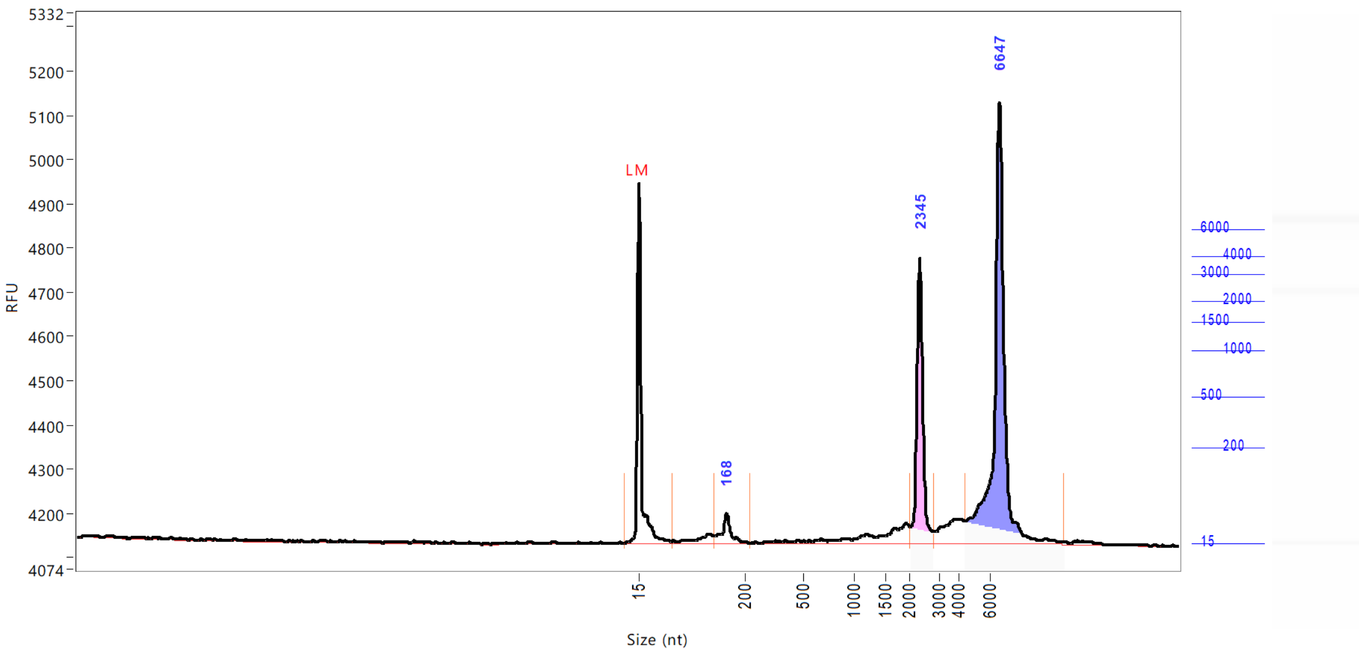


| Peak | Size (nt) | Conc. (ng/uL) | From (nt) | To (nt) | RFU |
|---|---|---|---|---|---|
| 1 | 15 (LM) | 0.5016 | 0 | 73 | 811 |
| 2 | 168 | 0.9662 | 145 | 221 | 66 |
| 3 | 2345 | 5.3180 | 2017 | 2803 | 645 |
| 4 | 6647 | 12.3892 | 4407 | 10810 | 998 |

| | | |
|---|---|---|
| TIC: | 18.6734 | ng/uL |
| TIM: | 31.1182 | nmole/L |
| Total Conc.: | 22.7015 | ng/uL |
| 28S/18S: | 2.2 | |
| RQN | 9.7 | |

Sample Peak Width (sec): 6    Sample Min Peak Height: 50    Sample Baseline V to V?: Y    Sample Baseline V to V pts: 3
Sample Filter: Binomial    # of Pts for Filter: 9    Sample Start Region (min): 0    Sample End Region (min): 40
Manual Baseline Start (min): 18    Manual Baseline End (min): 38
Marker Peak Width (sec): 6    Marker Min Peak Height: 100    Marker Baseline V to V?: Y    Marker Baseline V to V pts: 3
Lower Marker Selection: First Peak > 100 RFU    Upper Marker Selection: Last Peak > 100 RFU
Ladder Size (nt): 15, 200, 500, 1000, 1500, 2000, 3000, 4000, 6000
Quantification Using: Ladder    Final Concentration (ng/uL): 8.0000    Dilution Factor: 12.0
Min. RFU for Data Processing: 2

 Copyright 2015 Advanced Analytical Technologies, Inc. 

**Sample:** 103618-001-013
**Well Location:** E12
**Created:** Monday, April 08, 2019 11:44:17 AM

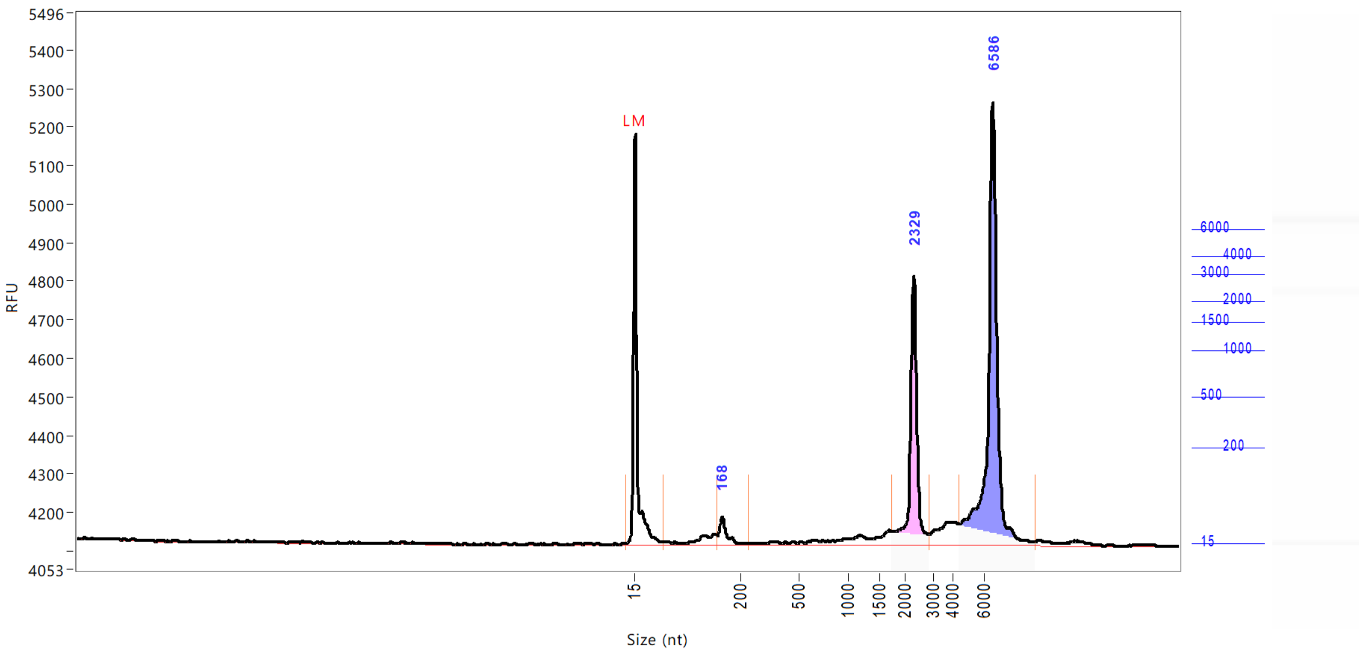


| Peak | Size (nt) | Conc. (ng/uL) | From (nt) | To (nt) | RFU |
|---|---|---|---|---|---|
| 1 | 15 (LM) | 0.5016 | 0 | 66 | 1062 |
| 2 | 168 | 0.6681 | 158 | 239 | 70 |
| 3 | 2329 | 4.9251 | 1750 | 2836 | 699 |
| 4 | 6586 | 10.2427 | 4407 | 9330 | 1150 |

| | | |
|---|---|---|
| TIC: | 15.8360 | ng/uL |
| TIM: | 23.8252 | nmole/L |
| Total Conc.: | 19.3776 | ng/uL |
| 28S/18S: | 2.1 | |
| RQN | 9.9 | |

Sample Peak Width (sec): 6 Sample Min Peak Height: 50 Sample Baseline V to V?: Y Sample Baseline V to V pts: 3
Sample Filter: Binomial # of Pts for Filter: 9 Sample Start Region (min): 0 Sample End Region (min): 40
Manual Baseline Start (min): 18 Manual Baseline End (min): 38
Marker Peak Width (sec): 6 Marker Min Peak Height: 100 Marker Baseline V to V?: Y Marker Baseline V to V pts: 3
Lower Marker Selection: First Peak > 100 RFU Upper Marker Selection: Last Peak > 100 RFU
Ladder Size (nt): 15, 200, 500, 1000, 1500, 2000, 3000, 4000, 6000
Quantification Using: Ladder Final Concentration (ng/uL): 8.0000 Dilution Factor: 12.0
Min. RFU for Data Processing: 2

 Copyright 2015 Advanced Analytical Technologies, Inc. 

**Sample:** 103618-001-014
**Well Location:** F12
**Created:** Monday, April 08, 2019 11:44:17 AM

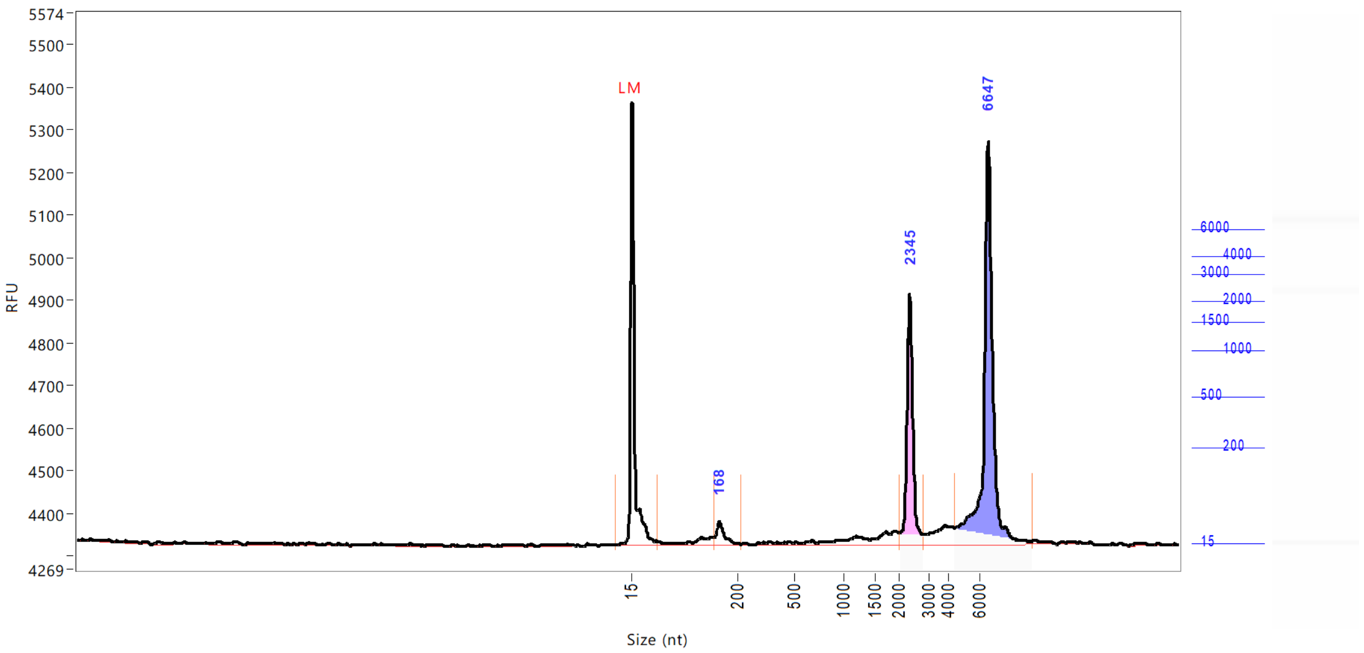


| Peak | Size (nt) | Conc. (ng/uL) | From (nt) | To (nt) | RFU |
|---|---|---|---|---|---|
| 1 | 15 (LM) | 0.5016 | 0 | 60 | 1038 |
| 2 | 168 | 0.5258 | 159 | 221 | 52 |
| 3 | 2345 | 3.6855 | 2017 | 2803 | 584 |
| 4 | 6647 | 8.2427 | 4438 | 9484 | 943 |

| | | |
|---|---|---|
| TIC: | 12.4540 | ng/uL |
| TIM: | 18.3882 | nmole/L |
| Total Conc.: | 15.0553 | ng/uL |

| | |
|---|---|
| 28S/18S: | 2.2 |
| RQN | 9.8 |

```
Sample Peak Width (sec): 6    Sample Min Peak Height: 50    Sample Baseline V to V?: Y    Sample Baseline V to V pts: 3
Sample Filter: Binomial       # of Pts for Filter: 9        Sample Start Region (min): 0  Sample End Region (min): 40
Manual Baseline Start (min): 18     Manual Baseline End (min): 38
Marker Peak Width (sec): 6    Marker Min Peak Height: 100   Marker Baseline V to V?: Y    Marker Baseline V to V pts: 3
Lower Marker Selection: First Peak > 100 RFU                Upper Marker Selection: Last Peak > 100 RFU
Ladder Size (nt): 15, 200, 500, 1000, 1500, 2000, 3000, 4000, 6000
Quantification Using: Ladder          Final Concentration (ng/uL): 8.0000          Dilution Factor: 12.0
Min. RFU for Data Processing: 2
```

 Copyright 2015 Advanced Analytical Technologies, Inc. 

**Sample:** 103618-001-015
**Well Location:** G12
**Created:** Monday, April 08, 2019 11:44:17 AM

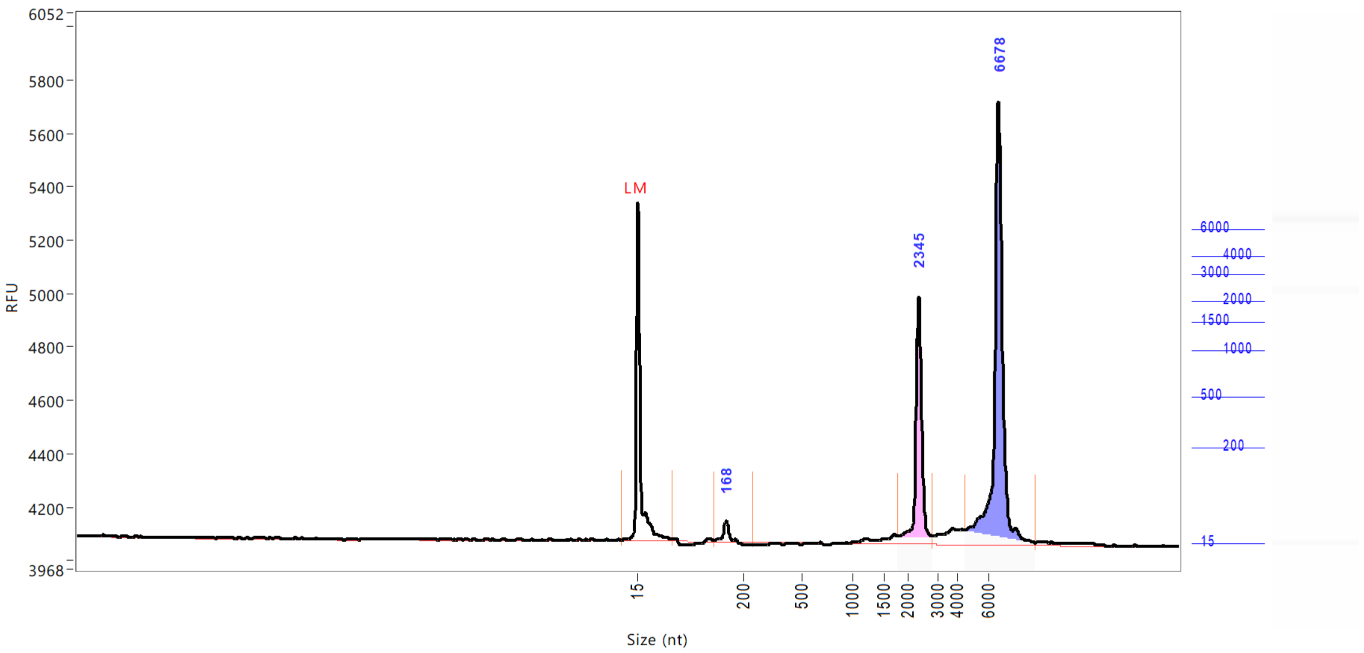


| Peak | Size (nt) | Conc. (ng/uL) | From (nt) | To (nt) | RFU |
|---|---|---|---|---|---|
| 1 | 15 (LM) | 0.5016 | 0 | 76 | 1264 |
| 2 | 168 | 0.4737 | 148 | 246 | 76 |
| 3 | 2345 | 4.8799 | 1790 | 2803 | 923 |
| 4 | 6678 | 10.8455 | 4501 | 9052 | 1659 |

| | | |
|---|---|---|
| TIC: | 16.1991 | ng/uL |
| TIM: | 20.5140 | nmole/L |
| Total Conc.: | 18.0342 | ng/uL |
| 28S/18S: | 2.2 | |
| RQN | 10.0 | |

Sample Peak Width (sec): 6  Sample Min Peak Height: 50  Sample Baseline V to V?: Y  Sample Baseline V to V pts: 3
Sample Filter: Binomial  # of Pts for Filter: 9  Sample Start Region (min): 0  Sample End Region (min): 40
Manual Baseline Start (min): 18  Manual Baseline End (min): 38
Marker Peak Width (sec): 6  Marker Min Peak Height: 100  Marker Baseline V to V?: Y  Marker Baseline V to V pts: 3
Lower Marker Selection: First Peak > 100 RFU  Upper Marker Selection: Last Peak > 100 RFU
Ladder Size (nt): 15, 200, 500, 1000, 1500, 2000, 3000, 4000, 6000
Quantification Using: Ladder  Final Concentration (ng/uL): 8.0000  Dilution Factor: 12.0
Min. RFU for Data Processing: 2

 Copyright 2015 Advanced Analytical Technologies, Inc. 

**Sample:** 103618-001-016
**Well Location:** H12
**Created:** Monday, April 08, 2019 11:44:17 AM

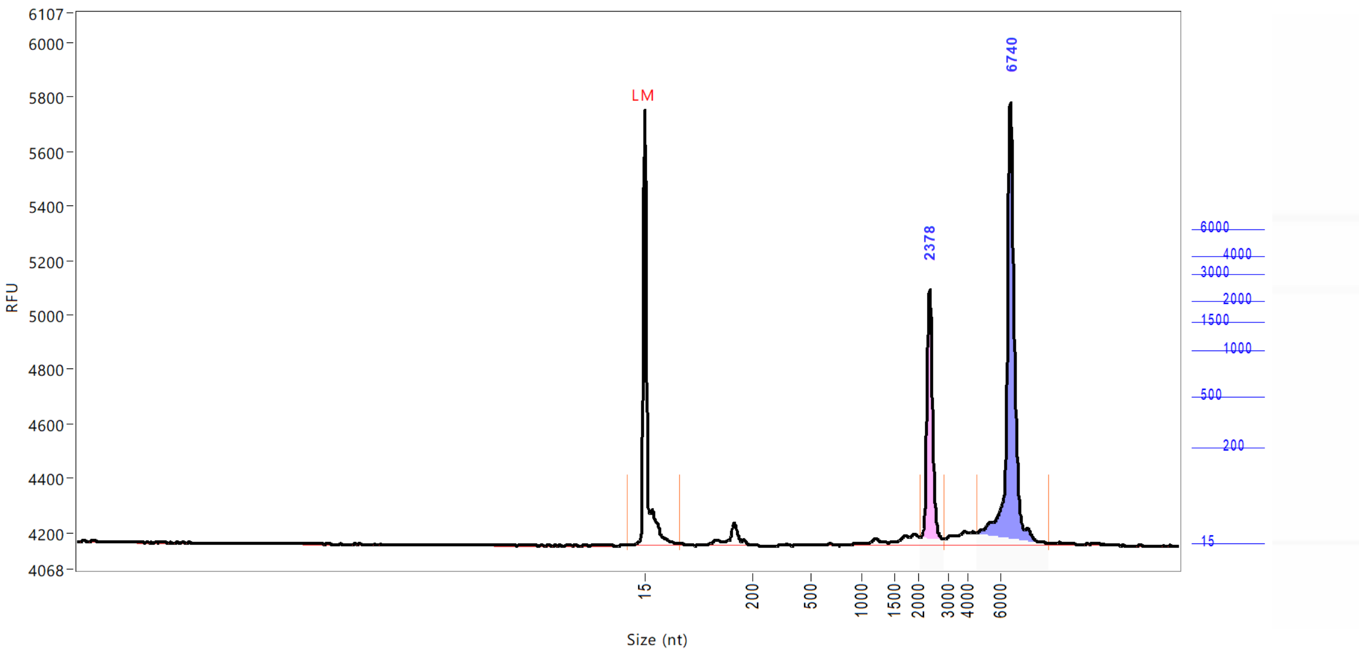


| Peak | Size (nt) | Conc. (ng/uL) | From (nt) | To (nt) | RFU |
|---|---|---|---|---|---|
| 1 | 15 (LM) | 0.5016 | 0 | 75 | 1596 |
| 2 | 2378 | 3.7010 | 2050 | 2836 | 938 |
| 3 | 6740 | 8.3015 | 4563 | 9176 | 1625 |

| | | |
|---|---|---|
| TIC: | 12.0024 | ng/uL |
| TIM: | 8.7648 | nmole/L |
| Total Conc.: | 14.2560 | ng/uL |
| 28S/18S: | 2.2 | |
| RQN | 10.0 | |

Sample Peak Width (sec): 6 Sample Min Peak Height: 200 Sample Baseline V to V?: Y Sample Baseline V to V pts: 3
Sample Filter: Binomial # of Pts for Filter: 9 Sample Start Region (min): 0 Sample End Region (min): 40
Manual Baseline Start (min): 18 Manual Baseline End (min): 38
Marker Peak Width (sec): 6 Marker Min Peak Height: 100 Marker Baseline V to V?: Y Marker Baseline V to V pts: 3
Lower Marker Selection: First Peak > 100 RFU Upper Marker Selection: Last Peak > 100 RFU
Ladder Size (nt): 15, 200, 500, 1000, 1500, 2000, 3000, 4000, 6000
Quantification Using: Ladder Final Concentration (ng/uL): 8.0000 Dilution Factor: 12.0
Min. RFU for Data Processing: 2

 Copyright 2015 Advanced Analytical Technologies, Inc. 

**Sample:** DNA Size Ladder
**Well Location:** H12
**Created:** Monday, April 08, 2019 11:44:17 AM
**Import From:** C:\Users\r.hennevelt\Desktop\FA ladders\RNA_SS_ladder.SCAL
**Fit Type:** Point to Point

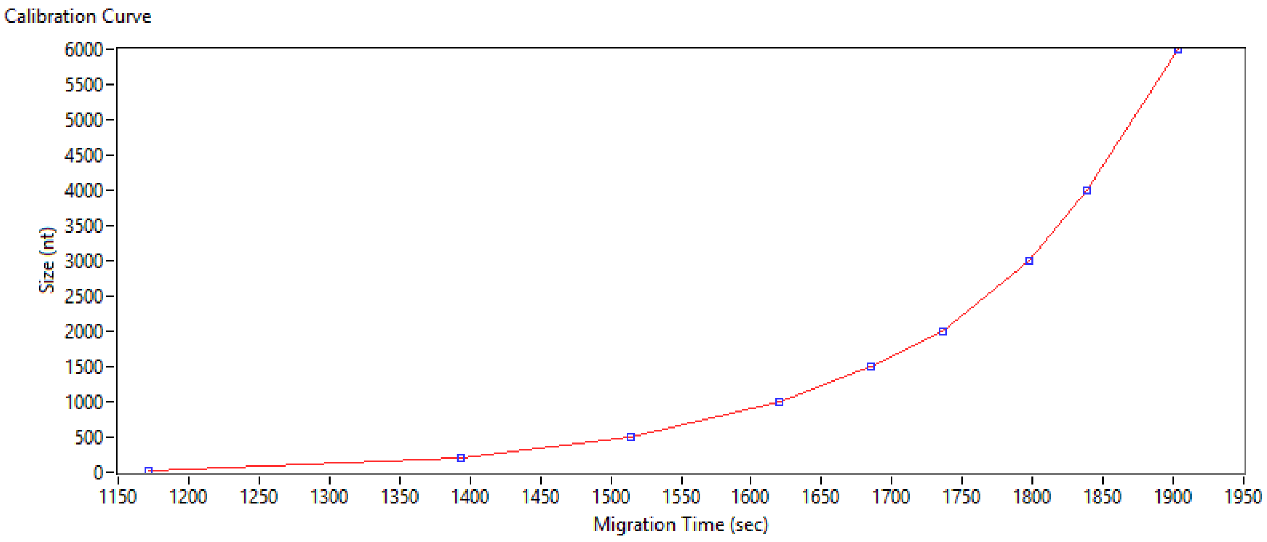


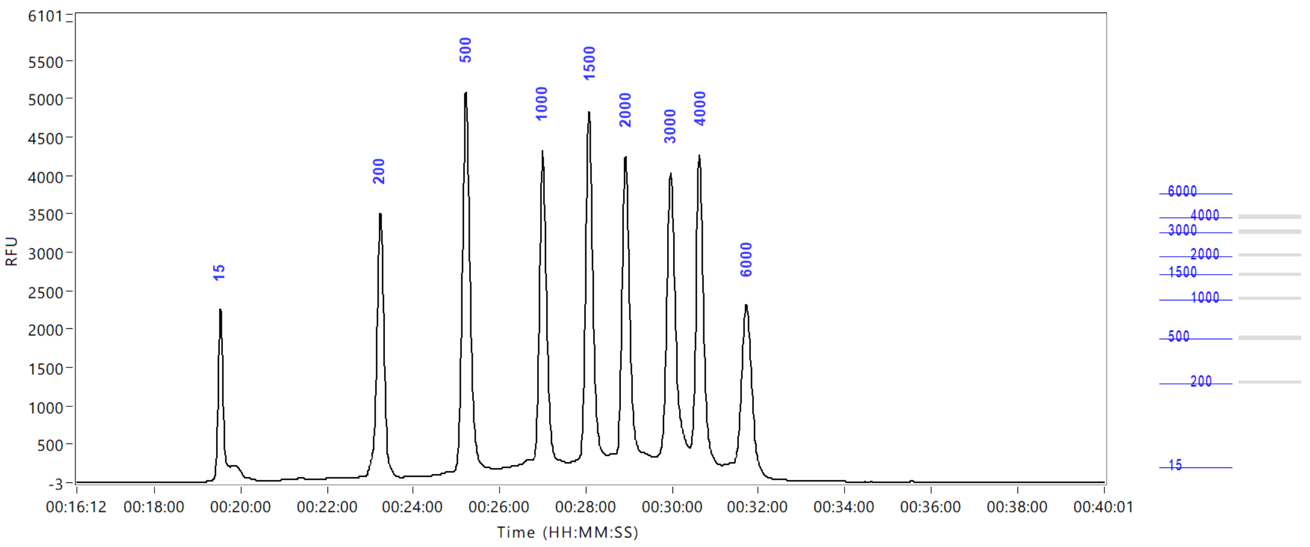


 Copyright 2015 Advanced Analytical Technologies, Inc. 

### *Fragment Analyzer* Run Summary:

**Filename and Data Path:** X:\Lopende Opdrachten\GAII\103618\EntryQC\EntryQC_103627-001_103618_RNA-SS 12-24-36\2019 04 08 12H 24M.raw
**Created:** Monday, April 08, 2019 12:57:12 PM
**# of Capillaries:** 2
**Array Serial #:** 112118-01SFS
**Effect Length:** 33 cm
**Array Usage Count:** 86
**FA Version #:** 1.2.0.11
**Device Serial #:** 3003

**METHOD INFORMATION**
**Method Name:** DNF-471-33 - SS Total RNA 15nt.mthds
**Gel Prime:** No
**Full Conditioning:** Yes
**Gel Prime to Buffer:** Yes
**Gel Selection:** Gel 1
**Perform Prerun:** 8.0 kV, 30 sec.
**Rinse:** No
**Marker 1:** No
**Rinse:** Tray: 3, Row: A, # Dips: 2
**Sample Injection:** 5.0 kV, 4 sec.
**Separation:** 8.0 kV, 40.0 min.
**Tray Name:** EntryQC_103627-001_103618_RNA-SS

**Analysis Mode:** RNA (Eukaryotic)

**NOTES**

 Copyright 2015 Advanced Analytical Technologies, Inc. 

### Gel Image

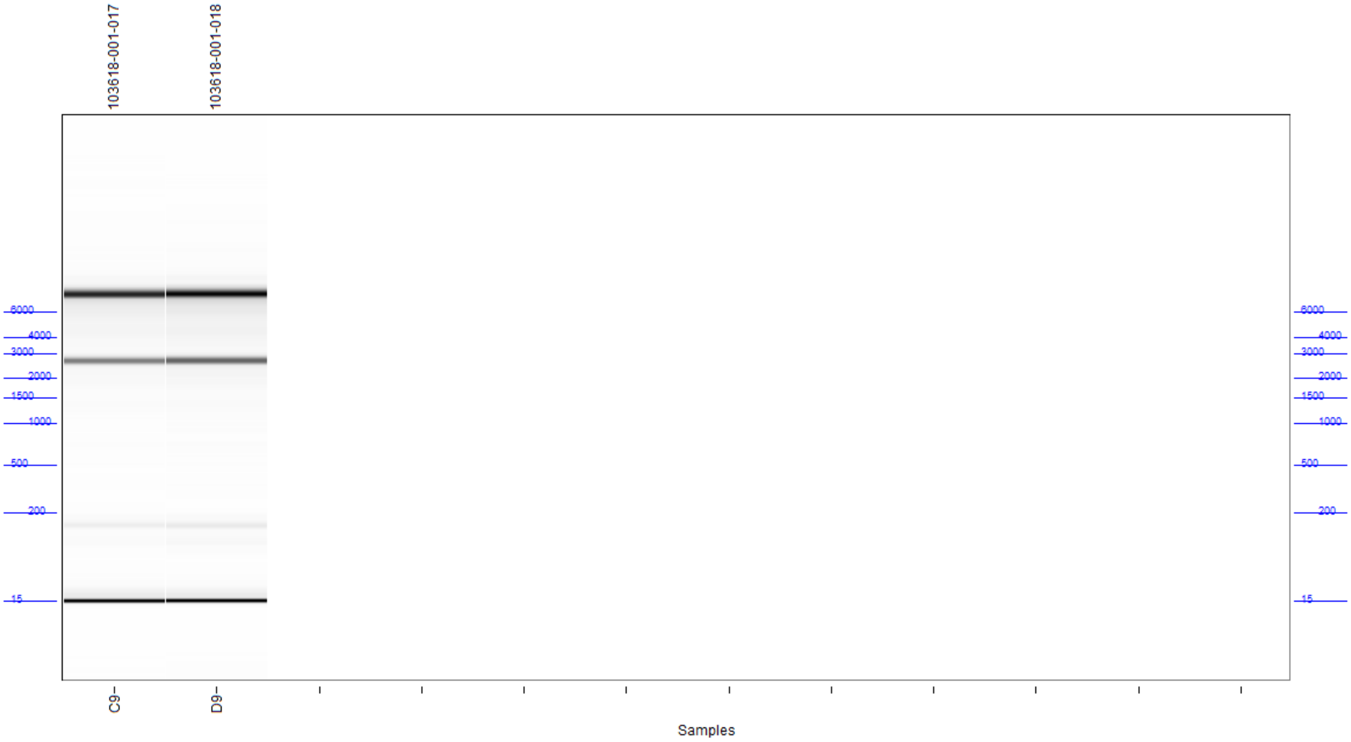



Copyright 2015 Advanced Analytical Technologies, Inc.

**Filename and Data Path:** X:\Lopende Opdrachten\GAII\103618\EntryQC\EntryQC_103627-001_103618_RNA-SS 12-24-36\ 2019 04 08 12H 24M.raw

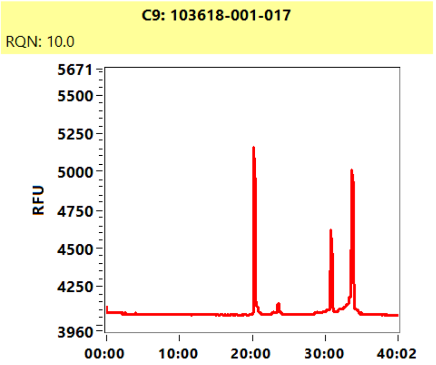


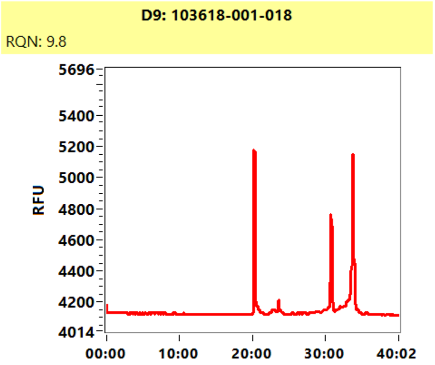

**Sample:** 103618-001-017
**Well Location:** C9
**Created:** Monday, April 08, 2019 12:57:12 PM

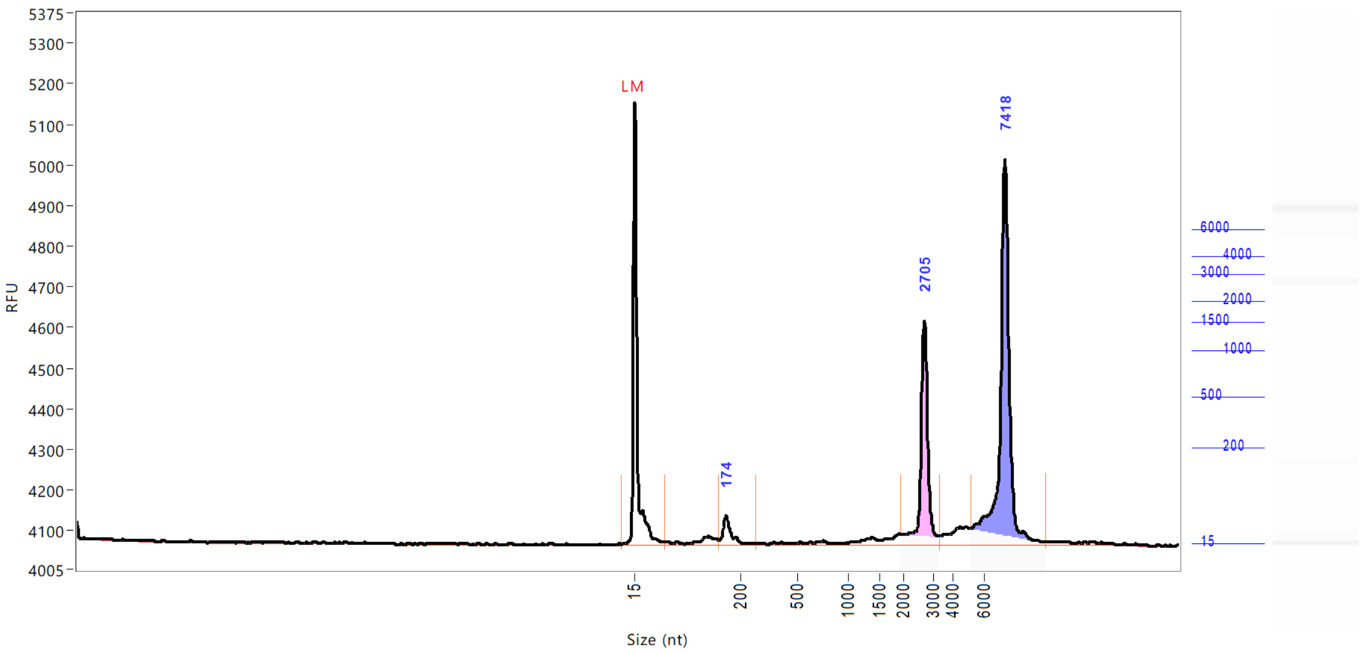


| Peak | Size (nt) | Conc. (ng/uL) | From (nt) | To (nt) | RFU |
|---|---|---|---|---|---|
| 1 | 15 (LM) | 0.5016 | 0 | 68 | 1089 |
| 2 | 174 | 0.6939 | 162 | 279 | 72 |
| 3 | 2705 | 3.8678 | 1950 | 3334 | 551 |
| 4 | 7418 | 8.0254 | 5219 | 10008 | 945 |

| | | |
|---|---|---|
| TIC: | 12.5871 | ng/uL |
| TIM: | 20.1734 | nmole/L |
| Total Conc.: | 14.5476 | ng/uL |
| 28S/18S: | 2.1 | |
| RQN | 10.0 | |

Sample Peak Width (sec): 6 Sample Min Peak Height: 50 Sample Baseline V to V?: Y Sample Baseline V to V pts: 3
Sample Filter: Binomial # of Pts for Filter: 9 Sample Start Region (min): 0 Sample End Region (min): 40
Manual Baseline Start (min): 18 Manual Baseline End (min): 38
Marker Peak Width (sec): 6 Marker Min Peak Height: 100 Marker Baseline V to V?: Y Marker Baseline V to V pts: 3
Lower Marker Selection: First Peak > 100 RFU Upper Marker Selection: Last Peak > 100 RFU
Ladder Size (nt): 15, 200, 500, 1000, 1500, 2000, 3000, 4000, 6000
Quantification Using: Ladder Final Concentration (ng/uL): 8.0000 Dilution Factor: 12.0
Min. RFU for Data Processing: 2

 Copyright 2015 Advanced Analytical Technologies, Inc. 

**Sample:** 103618-001-018
**Well Location:** D9
**Created:** Monday, April 08, 2019 12:57:12 PM

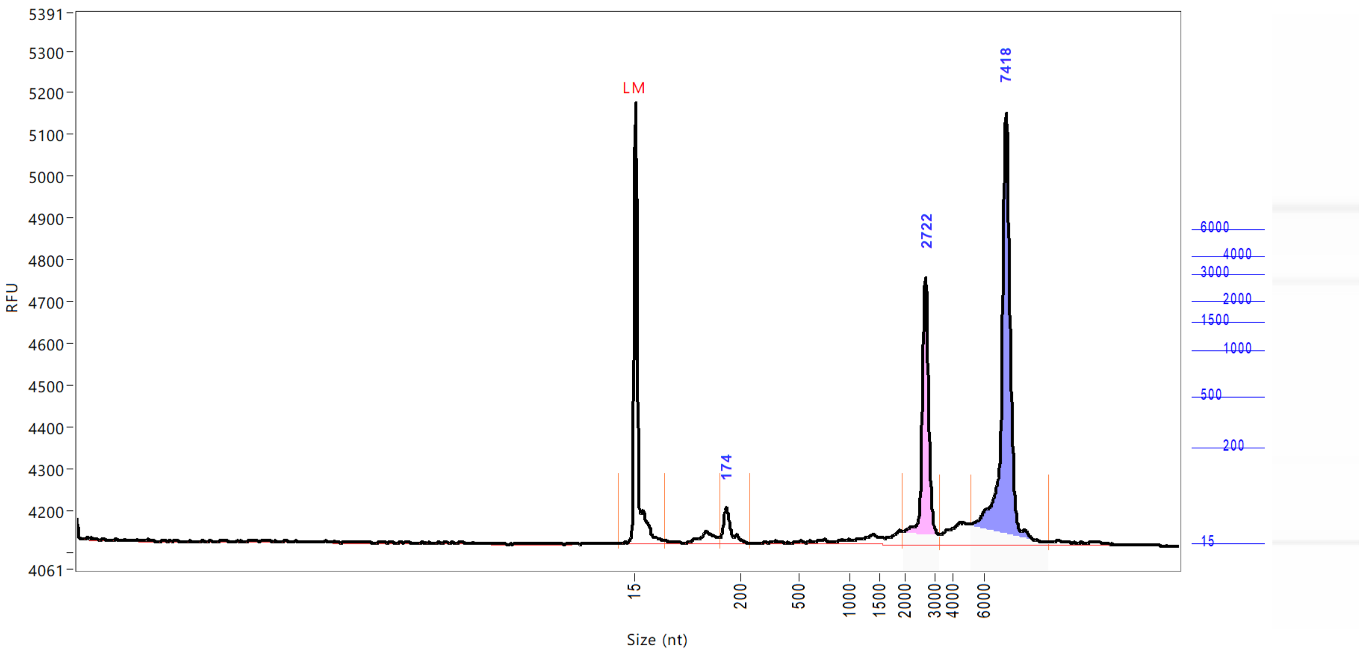


| Peak | Size (nt) | Conc. (ng/uL) | From (nt) | To (nt) | RFU |
|---|---|---|---|---|---|
| 1 | 15 (LM) | 0.5016 | 0 | 66 | 1054 |
| 2 | 174 | 0.8555 | 163 | 249 | 85 |
| 3 | 2722 | 4.7868 | 1970 | 3308 | 636 |
| 4 | 7418 | 9.5344 | 5094 | 10193 | 1031 |

| | | |
|---|---|---|
| TIC: | 15.1767 | ng/uL |
| TIM: | 24.7312 | nmole/L |
| Total Conc.: | 18.0436 | ng/uL |
| 28S/18S: | 2.0 | |
| RQN | 9.8 | |

Sample Peak Width (sec): 6 Sample Min Peak Height: 50 Sample Baseline V to V?: Y Sample Baseline V to V pts: 3
Sample Filter: Binomial # of Pts for Filter: 9 Sample Start Region (min): 0 Sample End Region (min): 40
Manual Baseline Start (min): 18 Manual Baseline End (min): 38
Marker Peak Width (sec): 6 Marker Min Peak Height: 100 Marker Baseline V to V?: Y Marker Baseline V to V pts: 3
Lower Marker Selection: First Peak > 100 RFU Upper Marker Selection: Last Peak > 100 RFU
Ladder Size (nt): 15, 200, 500, 1000, 1500, 2000, 3000, 4000, 6000
Quantification Using: Ladder Final Concentration (ng/uL): 8.0000 Dilution Factor: 12.0
Min. RFU for Data Processing: 2

 Copyright 2015 Advanced Analytical Technologies, Inc. 

**Sample:** DNA Size Ladder
**Well Location:** H12
**Created:** Monday, April 08, 2019 12:57:12 PM
**Import From:** C:\Users\r.hennevelt\Desktop\FA ladders\RNA_SS_ladder.SCAL
**Fit Type:** Point to Point

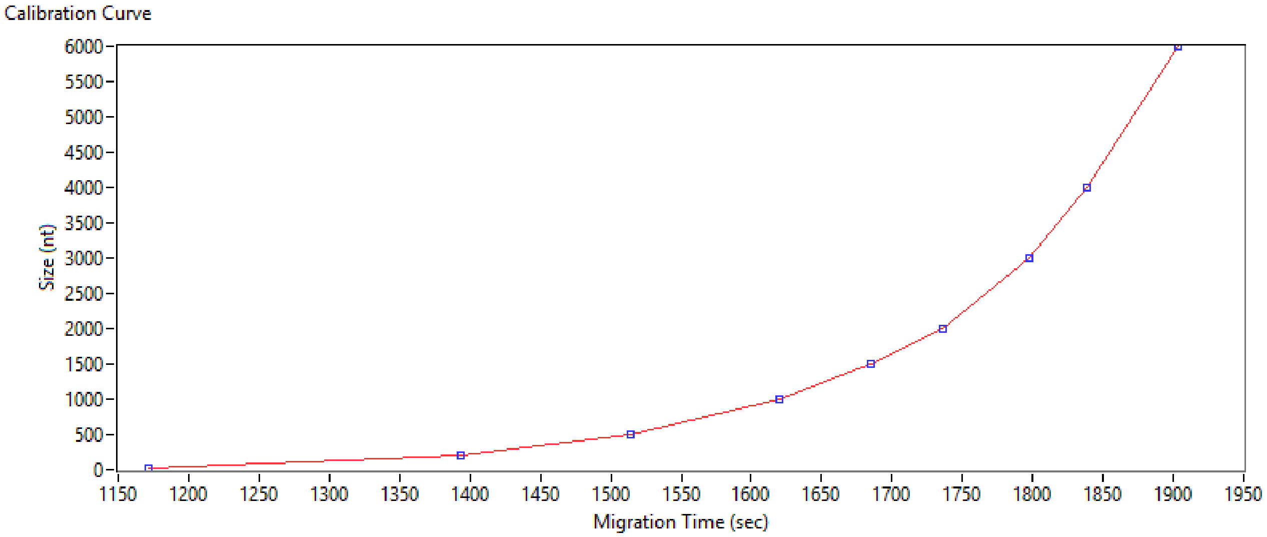


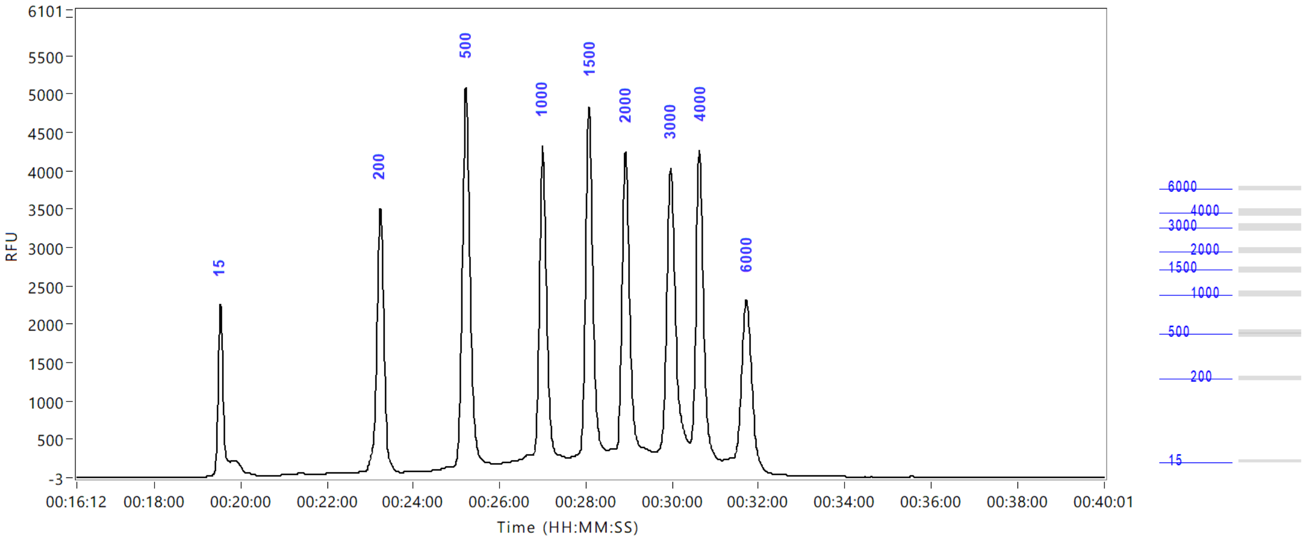


 Copyright 2015 Advanced Analytical Technologies, Inc. 

### *Fragment Analyzer* Run Summary:

**Filename and Data Path:** X:\Lopende Opdrachten\GAII\103639\PrepQC\PrepQC_103639_103655_7075-
Filename and Data Path: 05_103626_103618_103650_7060-02 11-00-09\2019 05 02 11H 00M.raw
**Created:** Thursday, May 02, 2019 11:28:53 AM
**# of Capillaries:** 18
**Array Serial #:** 112118-01SFS
**Effect Length:** 33 cm
**Array Usage Count:** 142
**FA Version #:** 1.2.0.11
**Device Serial #:** 3003

**METHOD INFORMATION**
**Method Name:** DNF-474-33 - HS NGS Fragment 1-6000bp.mthds
**Gel Prime:** No
**Full Conditioning:** Yes
**Gel Prime to Buffer:** No
**Gel Selection:** Gel 1
**Perform Prerun:** 6.0 kV, 30 sec.
**Rinse:** No
**Marker 1:** No
**Rinse:** Tray: 3, Row: A, # Dips: 1
**Sample Injection:** 5.0 kV, 30 sec.
**Separation:** 6.0 kV, 50.0 min.
**Tray Name:** PrepQC_103639_103655_7075-05_103626_103618_103650_7060-02

**Analysis Mode:** NGS

**NOTES**

 Copyright 2015 Advanced Analytical Technologies, Inc. 

### Gel Image

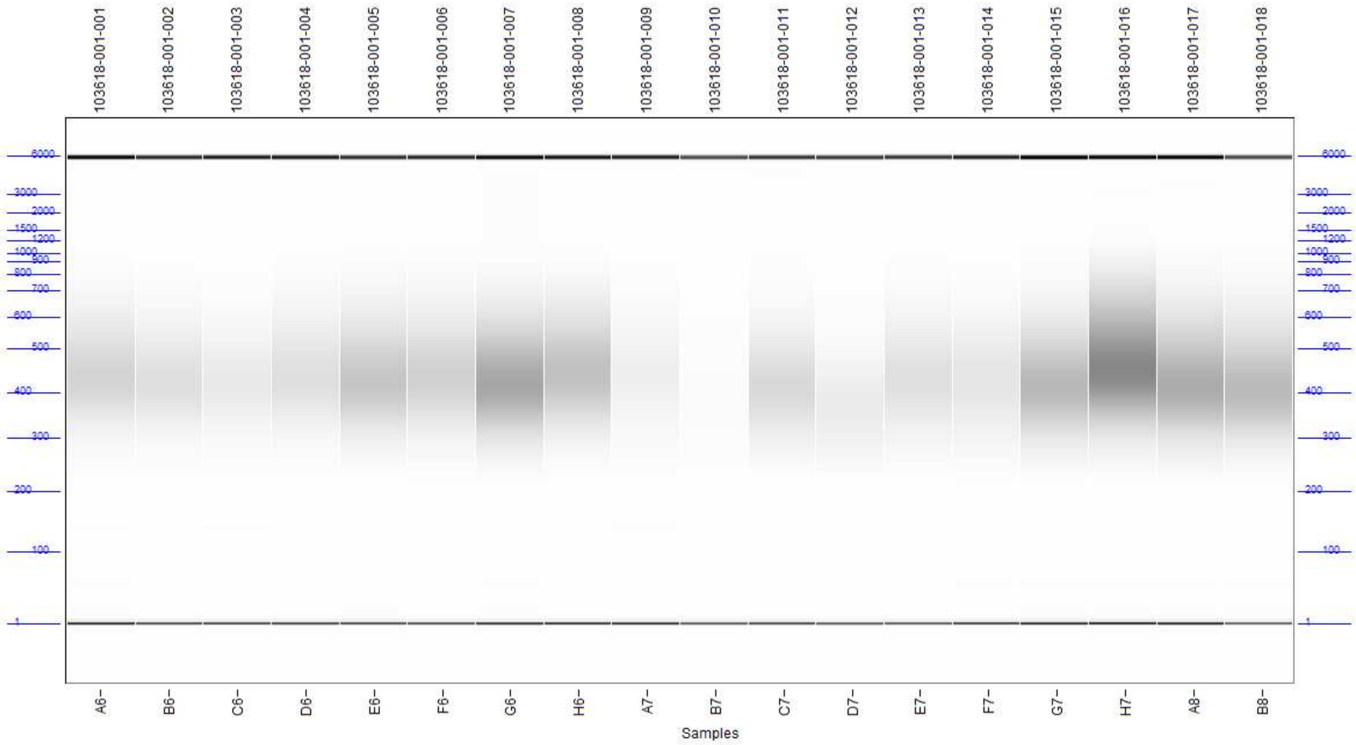



Copyright 2015 Advanced Analytical Technologies, Inc.

**Filename and Data Path:** X:\Lopende Opdrachten\GAII\103639\PrepQC\PrepQC_103639_103655_7075-05_103626_103618_103650_7060-02 11-00-09\2019 05 02 11H 00M.raw

A6: 103618-001-001

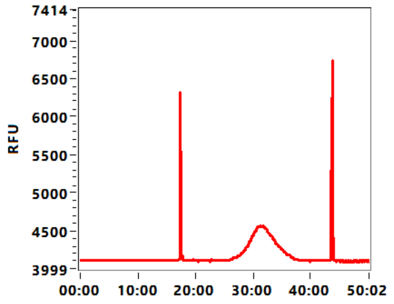


B6: 103618-001-002

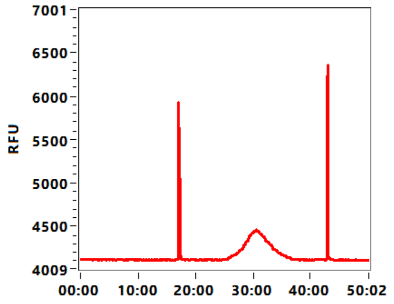


C6: 103618-001-003

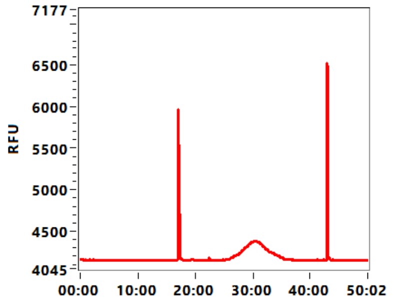


D6: 103618-001-004

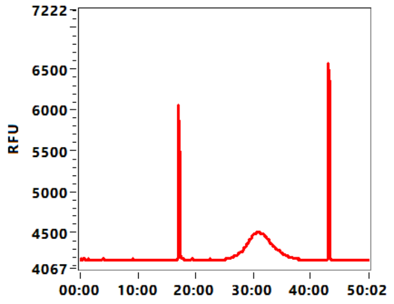


E6: 103618-001-005

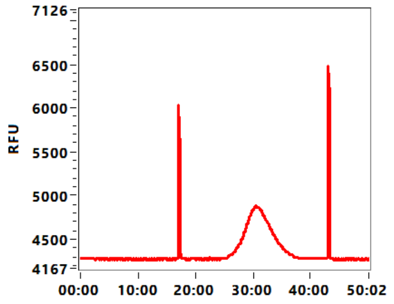


F6: 103618-001-006

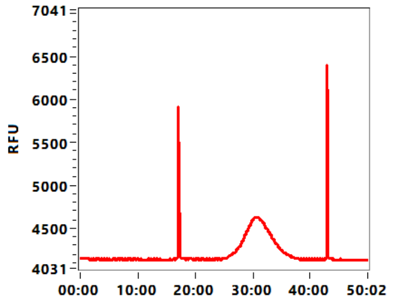


G6: 103618-001-007

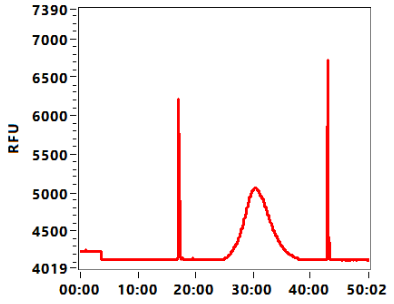


H6: 103618-001-008

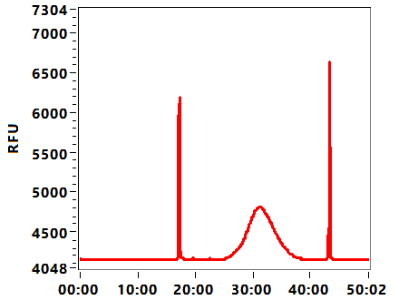


A7: 103618-001-009

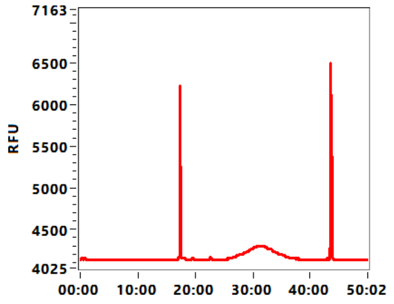


B7: 103618-001-010

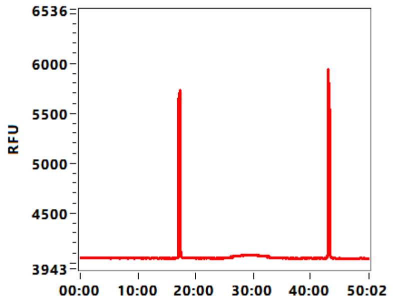


C7: 103618-001-011

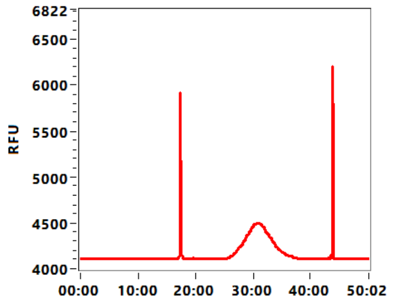


D7: 103618-001-012

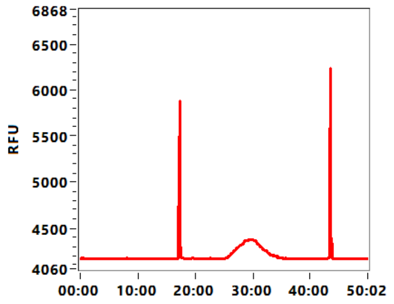



Copyright 2015 Advanced Analytical Technologies, Inc.

**Filename and Data Path:** X:\Lopende Opdrachten\GAII\103639\PrepQC\PrepQC_103639_103655_7075-05_103626_103618_103650_7060-02 11-00-09\2019 05 02 11H 00M.raw

E7: 103618-001-013

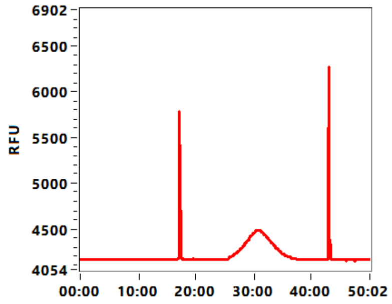


F7: 103618-001-014

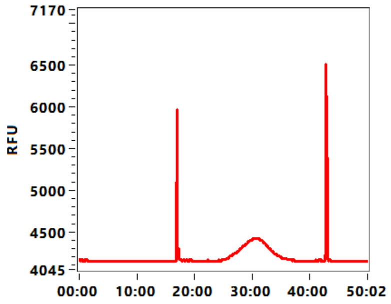


G7: 103618-001-015

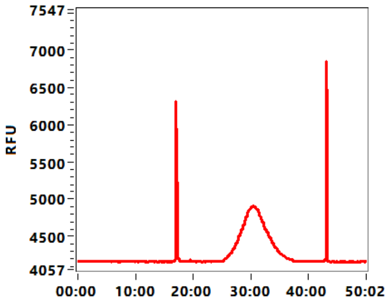


H7: 103618-001-016

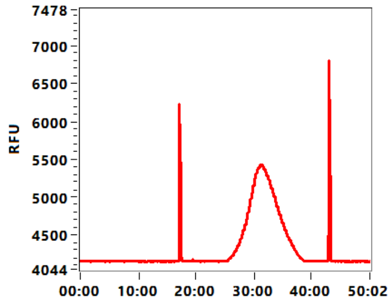


A8: 103618-001-017

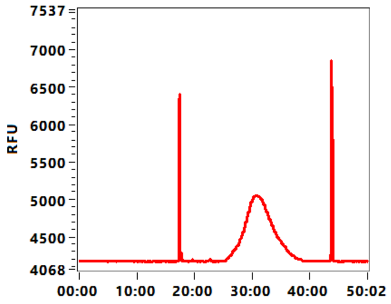


B8: 103618-001-018

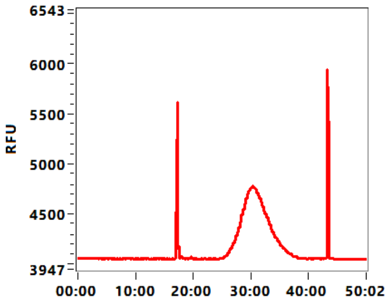



Copyright 2015 Advanced Analytical Technologies, Inc.

**Sample:** 103618-001-001
**Well Location:** A6
**Created:** Thursday, May 02, 2019 11:28:53 AM

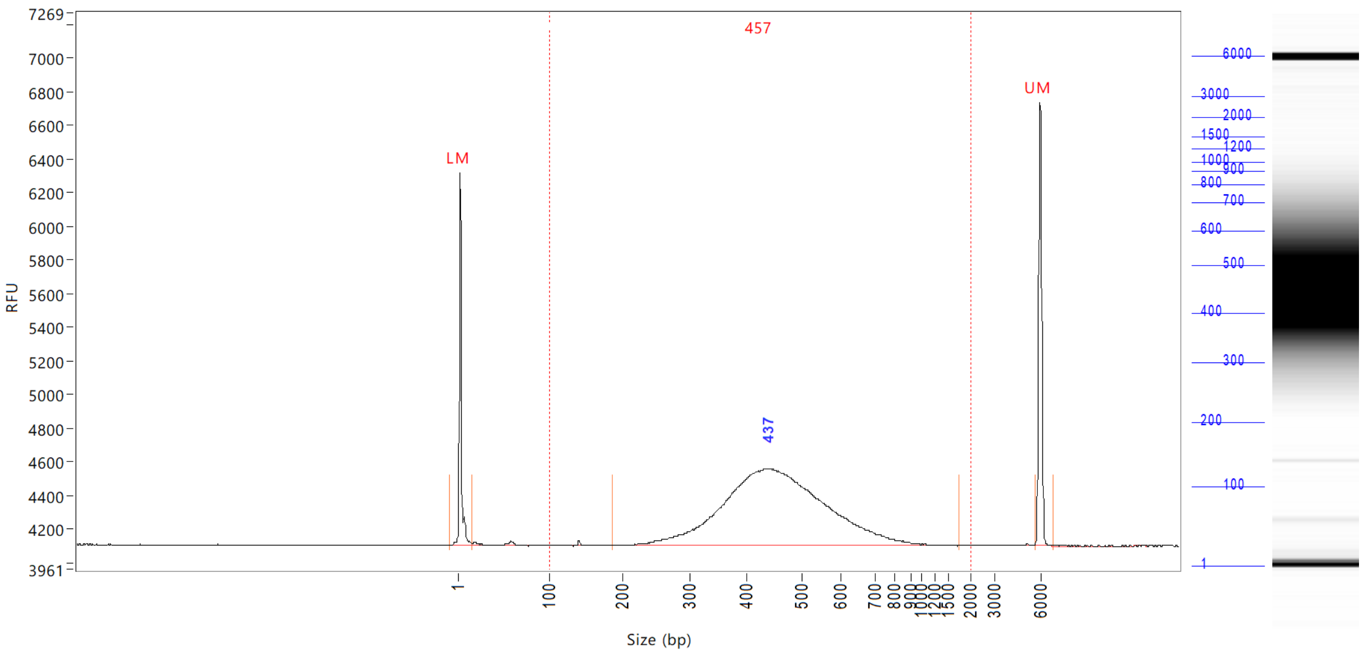


| Peak | Size (bp) | Conc. (ng/uL) | From (bp) | To (bp) | RFU |
|---|---|---|---|---|---|
| 1 | 1 (LM) | 0.0104 | 0 | 15 | 2214 |
| 2 | 437 | 0.7180 | 185 | 1759 | 455 |
| 3 | 6000 (UM) | 0.0076 | 5640 | 6883 | 2641 |

| | | |
|---|---|---|
| TIC: | 0.7180 | ng/uL |
| TIM: | 2.5853 | nmole/L |
| Total Conc.: | 0.7288 | ng/uL |

Smear Analysis 100 bp to 2000 bp 0.7198 ng/ul 98.8 %Total 2.5905 nmole/L 457 Avg. Size (b.p.) 25.47 %CV

Sample Peak Width (sec): 50 Sample Min Peak Height: 25 Sample Baseline V to V?: Y Sample Baseline V to V pts: 3
Sample Filter: Binomial # of Pts for Filter: 3 Sample Start Region (min): 0 Sample End Region (min): 50
Manual Baseline Start (min): 10 Manual Baseline End (min): 48
Marker Peak Width (sec): 5 Marker Min Peak Height: 200 Marker Baseline V to V?: Y Marker Baseline V to V pts: 3
Lower Marker Selection: First Peak > 200 RFU Upper Marker Selection: Last Peak > 200 RFU
Ladder Size (bp): 1, 100, 200, 300, 400, 500, 600, 700, 800, 900, 1000, 1200, 1500, 2000, 3000, 6000
Quantification Using: Ladder Final Concentration (ng/uL): 0.0830 Dilution Factor: 12.0


Copyright 2015 Advanced Analytical Technologies, Inc.

**Sample:** 103618-001-002
**Well Location:** B6
**Created:** Thursday, May 02, 2019 11:28:53 AM

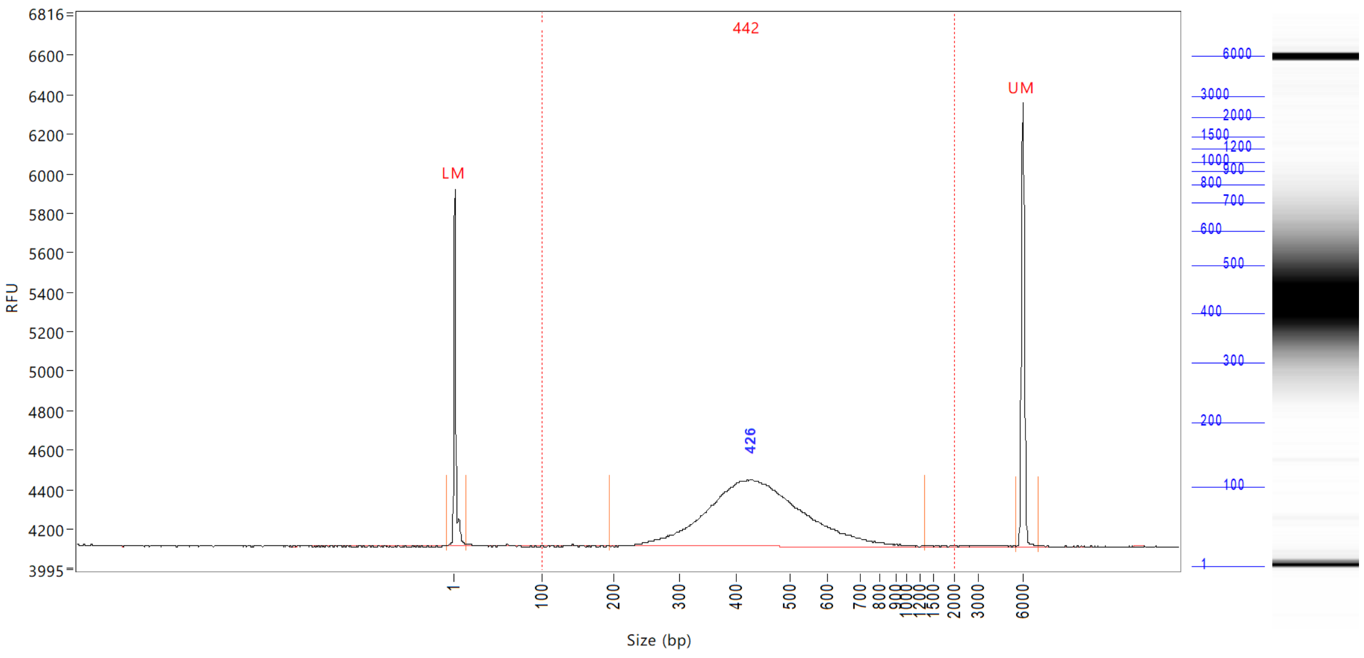


| Peak | Size (bp) | Conc. (ng/uL) | From (bp) | To (bp) | RFU |
|---|---|---|---|---|---|
| 1 | 1 (LM) | 0.0104 | 0 | 15 | 1808 |
| 2 | 426 | 0.6325 | 193 | 1324 | 337 |
| 3 | 6000 (UM) | 0.0083 | 5520 | 7026 | 2253 |

| | | |
|---|---|---|
| TIC: | 0.6325 | ng/uL |
| TIM: | 2.3601 | nmole/L |
| Total Conc.: | 0.6408 | ng/uL |

Smear Analysis 100 bp to 2000 bp 0.6341 ng/ul 99.0 %Total 2.3591 nmole/L 442 Avg. Size (b.p.) 25.22 %CV

Sample Peak Width (sec): 50 Sample Min Peak Height: 25 Sample Baseline V to V?: Y Sample Baseline V to V pts: 3
Sample Filter: Binomial # of Pts for Filter: 3 Sample Start Region (min): 0 Sample End Region (min): 50
Manual Baseline Start (min): 10 Manual Baseline End (min): 48
Marker Peak Width (sec): 5 Marker Min Peak Height: 200 Marker Baseline V to V?: Y Marker Baseline V to V pts: 3
Lower Marker Selection: First Peak > 200 RFU Upper Marker Selection: Last Peak > 200 RFU
Ladder Size (bp): 1, 100, 200, 300, 400, 500, 600, 700, 800, 900, 1000, 1200, 1500, 2000, 3000, 6000
Quantification Using: Ladder Final Concentration (ng/uL): 0.0830 Dilution Factor: 12.0

 Copyright 2015 Advanced Analytical Technologies, Inc. 

**Sample:** 103618-001-003
**Well Location:** C6
**Created:** Thursday, May 02, 2019 11:28:53 AM

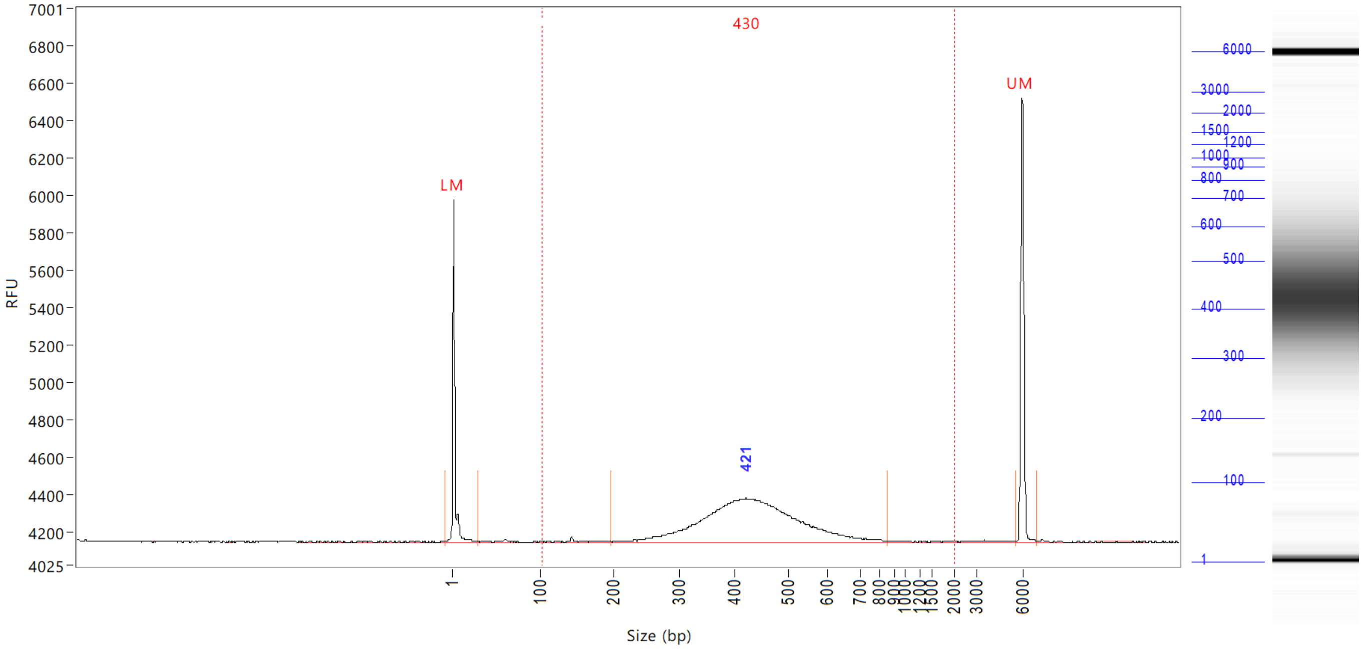


| Peak | Size (bp) | Conc. (ng/uL) | From (bp) | To (bp) | RFU |
|---|---|---|---|---|---|
| 1 | 1 (LM) | 0.0104 | 0 | 30 | 1828 |
| 2 | 421 | 0.4042 | 197 | 852 | 232 |
| 3 | 6000 (UM) | 0.0082 | 5616 | 6979 | 2376 |

| | | |
|---|---|---|
| TIC: | 0.4042 | ng/uL |
| TIM: | 1.5467 | nmole/L |
| Total Conc.: | 0.4163 | ng/uL |

| Smear Analysis | 100 bp to 2000 bp | 0.4104 ng/ul | 98.6 %Total | 1.5707 nmole/L | 430 Avg. Size (b.p.) | 26.30 %CV |
|---|---|---|---|---|---|---|

Sample Peak Width (sec): 50 Sample Min Peak Height: 25 Sample Baseline V to V?: Y Sample Baseline V to V pts: 3
Sample Filter: Binomial # of Pts for Filter: 3 Sample Start Region (min): 0 Sample End Region (min): 50
Manual Baseline Start (min): 10 Manual Baseline End (min): 48
Marker Peak Width (sec): 5 Marker Min Peak Height: 200 Marker Baseline V to V?: Y Marker Baseline V to V pts: 3
Lower Marker Selection: First Peak > 200 RFU Upper Marker Selection: Last Peak > 200 RFU
Ladder Size (bp): 1, 100, 200, 300, 400, 500, 600, 700, 800, 900, 1000, 1200, 1500, 2000, 3000, 6000
Quantification Using: Ladder Final Concentration (ng/uL): 0.0830 Dilution Factor: 12.0

 Copyright 2015 Advanced Analytical Technologies, Inc. 

**Sample:** 103618-001-004
**Well Location:** D6
**Created:** Thursday, May 02, 2019 11:28:53 AM

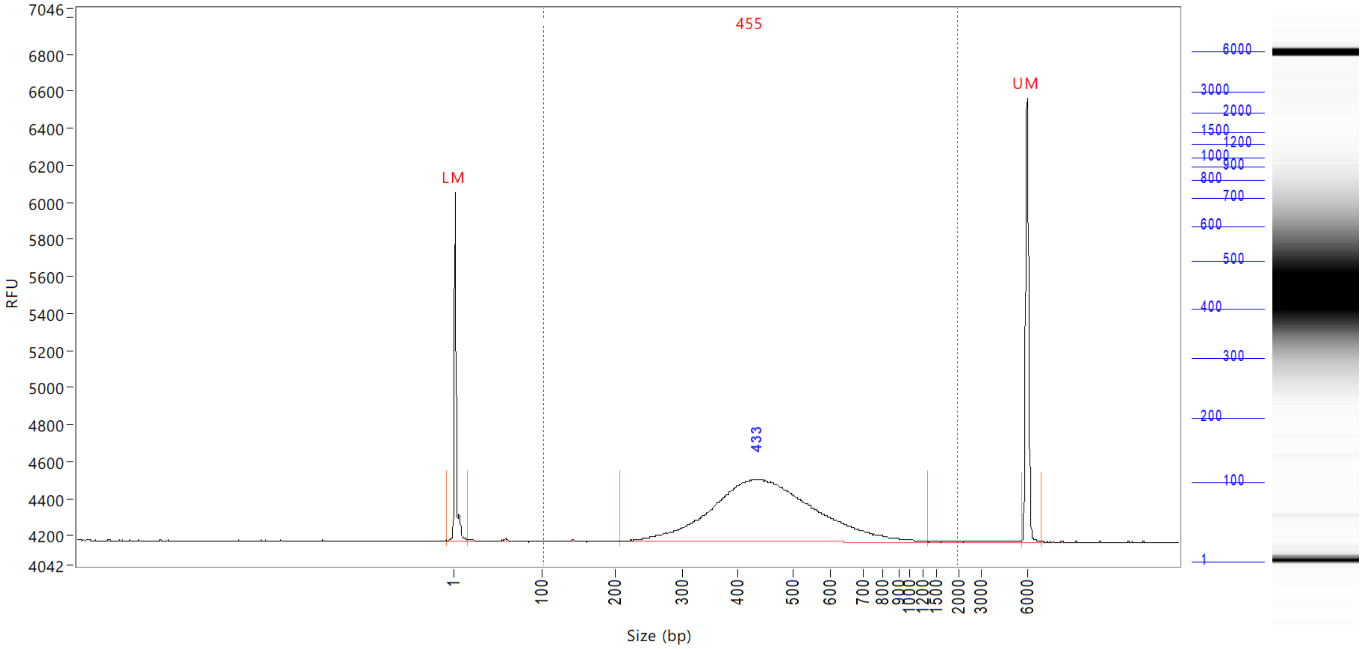


| Peak | Size (bp) | Conc. (ng/uL) | From (bp) | To (bp) | RFU |
|---|---|---|---|---|---|
| 1 | 1 (LM) | 0.0104 | 0 | 16 | 1884 |
| 2 | 433 | 0.6358 | 205 | 1315 | 337 |
| 3 | 6000 (UM) | 0.0084 | 5664 | 6883 | 2400 |

| | | |
|---|---|---|
| TIC: | 0.6358 | ng/uL |
| TIM: | 2.2991 | nmole/L |
| Total Conc.: | 0.6493 | ng/uL |

Smear Analysis 100 bp to 2000 bp 0.6396 ng/ul 98.5 %Total 2.3109 nmole/L 455 Avg. Size (b.p.) 27.35 %CV

Sample Peak Width (sec): 50 Sample Min Peak Height: 25 Sample Baseline V to V?: Y Sample Baseline V to V pts: 3
Sample Filter: Binomial # of Pts for Filter: 3 Sample Start Region (min): 0 Sample End Region (min): 50
Manual Baseline Start (min): 10 Manual Baseline End (min): 48
Marker Peak Width (sec): 5 Marker Min Peak Height: 200 Marker Baseline V to V?: Y Marker Baseline V to V pts: 3
Lower Marker Selection: First Peak > 200 RFU Upper Marker Selection: Last Peak > 200 RFU
Ladder Size (bp): 1, 100, 200, 300, 400, 500, 600, 700, 800, 900, 1000, 1200, 1500, 2000, 3000, 6000
Quantification Using: Ladder Final Concentration (ng/uL): 0.0830 Dilution Factor: 12.0

 Copyright 2015 Advanced Analytical Technologies, Inc. 

**Sample:** 103618-001-005
**Well Location:** E6
**Created:** Thursday, May 02, 2019 11:28:53 AM

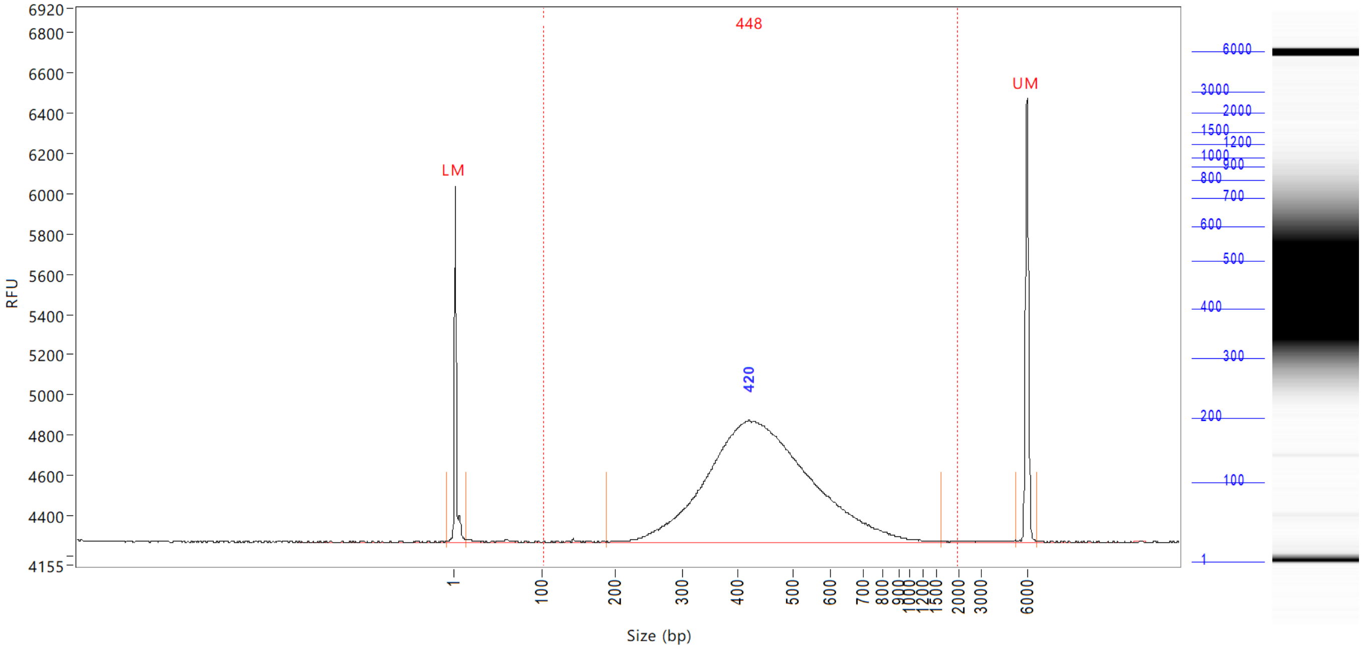


| Peak | Size (bp) | Conc. (ng/uL) | From (bp) | To (bp) | RFU |
|---|---|---|---|---|---|
| 1 | 1 (LM) | 0.0104 | 0 | 14 | 1763 |
| 2 | 420 | 1.2201 | 188 | 1626 | 605 |
| 3 | 6000 (UM) | 0.0083 | 5280 | 6597 | 2209 |

| | | |
|---|---|---|
| TIC: | 1.2201 | ng/uL |
| TIM: | 4.4811 | nmole/L |
| Total Conc.: | 1.2363 | ng/uL |

Smear Analysis 100 bp to 2000 bp 1.2244 ng/ul 99.0 %Total 4.4932 nmole/L 448 Avg. Size (b.p.) 27.19 %CV

Sample Peak Width (sec): 50 Sample Min Peak Height: 25 Sample Baseline V to V?: Y Sample Baseline V to V pts: 3
Sample Filter: Binomial # of Pts for Filter: 3 Sample Start Region (min): 0 Sample End Region (min): 50
Manual Baseline Start (min): 10 Manual Baseline End (min): 48
Marker Peak Width (sec): 5 Marker Min Peak Height: 200 Marker Baseline V to V?: Y Marker Baseline V to V pts: 3
Lower Marker Selection: First Peak > 200 RFU Upper Marker Selection: Last Peak > 200 RFU
Ladder Size (bp): 1, 100, 200, 300, 400, 500, 600, 700, 800, 900, 1000, 1200, 1500, 2000, 3000, 6000
Quantification Using: Ladder Final Concentration (ng/uL): 0.0830 Dilution Factor: 12.0


Copyright 2015 Advanced Analytical Technologies, Inc.

**Sample:** 103618-001-006
**Well Location:** F6
**Created:** Thursday, May 02, 2019 11:28:53 AM

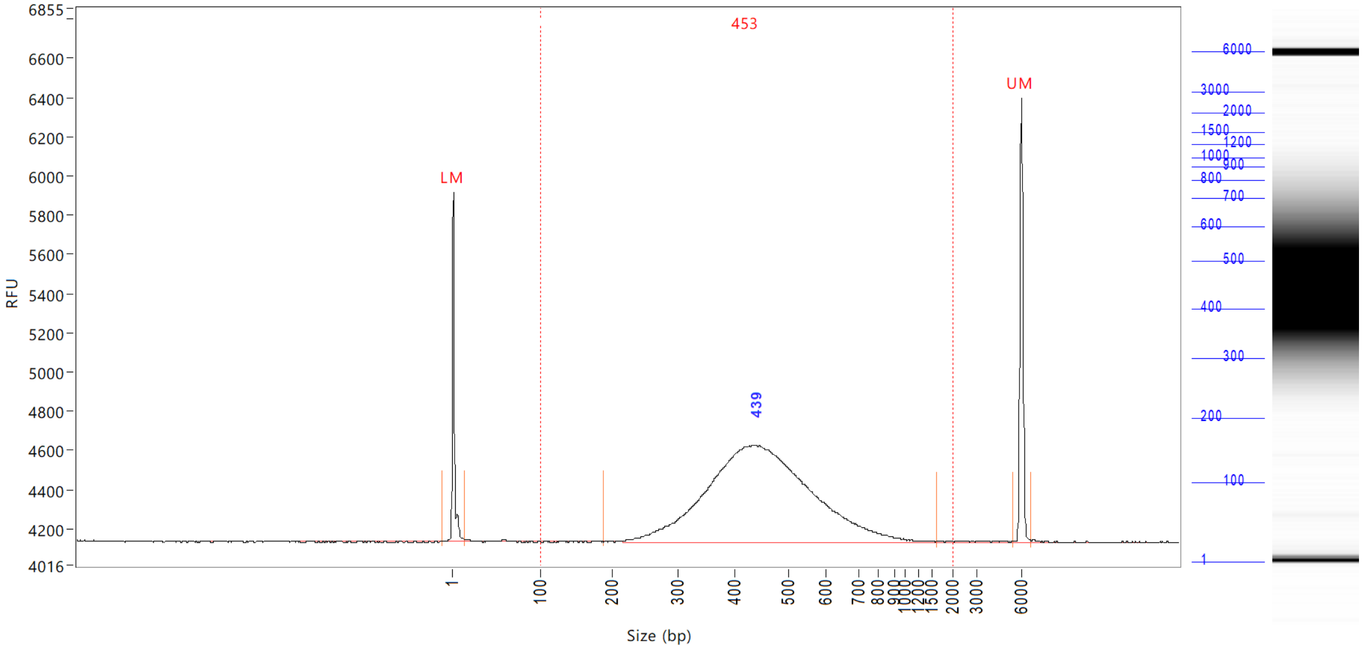


| Peak | Size (bp) | Conc. (ng/uL) | From (bp) | To (bp) | RFU |
|---|---|---|---|---|---|
| 1 | 1 (LM) | 0.0104 | 0 | 15 | 1783 |
| 2 | 439 | 1.0040 | 186 | 1609 | 496 |
| 3 | 6000 (UM) | 0.0084 | 5448 | 6644 | 2268 |

| | | |
|---|---|---|
| TIC: | 1.0040 | ng/uL |
| TIM: | 3.6466 | nmole/L |
| Total Conc.: | 1.0140 | ng/uL |

Smear Analysis 100 bp to 2000 bp 1.0060 ng/ul 99.2 %Total 3.6516 nmole/L 453 Avg. Size (b.p.) 26.44 %CV

Sample Peak Width (sec): 50 Sample Min Peak Height: 25 Sample Baseline V to V?: Y Sample Baseline V to V pts: 3
Sample Filter: Binomial # of Pts for Filter: 3 Sample Start Region (min): 0 Sample End Region (min): 50
Manual Baseline Start (min): 10 Manual Baseline End (min): 48
Marker Peak Width (sec): 5 Marker Min Peak Height: 200 Marker Baseline V to V?: Y Marker Baseline V to V pts: 3
Lower Marker Selection: First Peak > 200 RFU Upper Marker Selection: Last Peak > 200 RFU
Ladder Size (bp): 1, 100, 200, 300, 400, 500, 600, 700, 800, 900, 1000, 1200, 1500, 2000, 3000, 6000
Quantification Using: Ladder Final Concentration (ng/uL): 0.0830 Dilution Factor: 12.0

 Copyright 2015 Advanced Analytical Technologies, Inc. 

**Sample:** 103618-001-007
**Well Location:** G6
**Created:** Thursday, May 02, 2019 11:28:53 AM

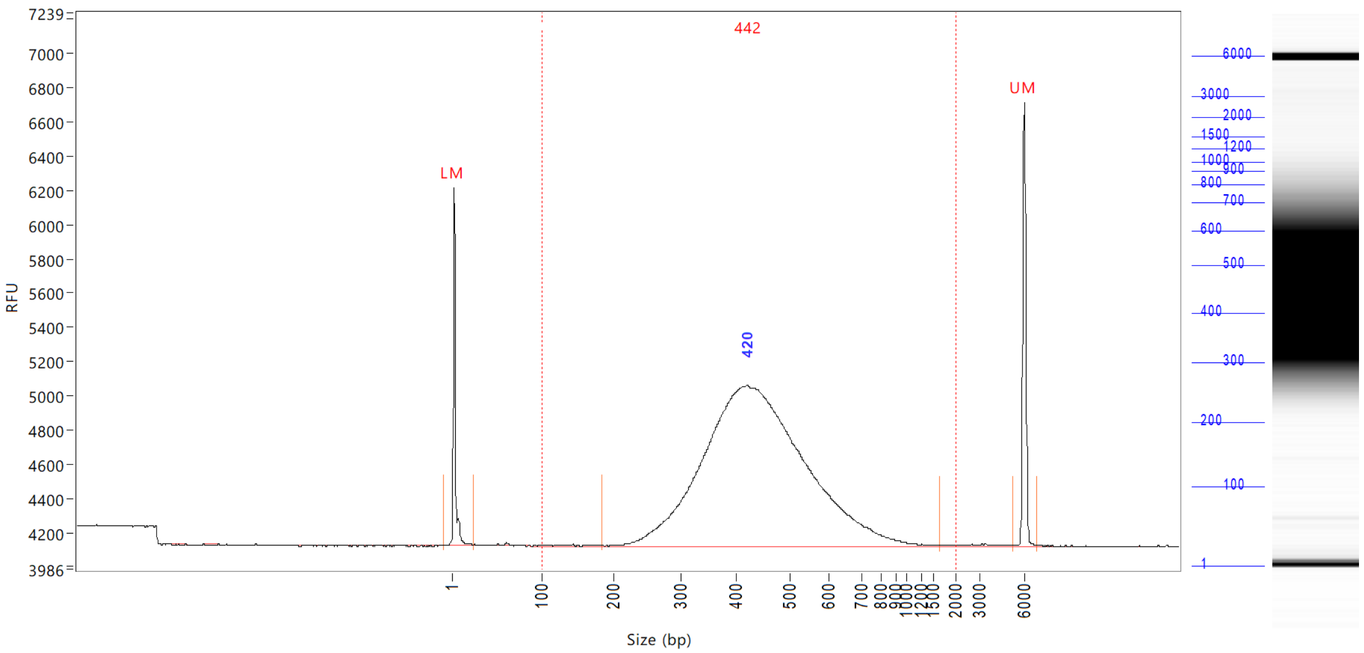


| Peak | Size (bp) | Conc. (ng/uL) | From (bp) | To (bp) | RFU |
|---|---|---|---|---|---|
| 1 | 1 (LM) | 0.0104 | 0 | 23 | 2091 |
| 2 | 420 | 1.5537 | 182 | 1651 | 939 |
| 3 | 6000 (UM) | 0.0081 | 5280 | 6788 | 2599 |

| | | |
|---|---|---|
| TIC: | 1.5537 | ng/uL |
| TIM: | 5.7968 | nmole/L |
| Total Conc.: | 1.5681 | ng/uL |

Smear Analysis 100 bp to 2000 bp 1.5571 ng/ul 99.3 %Total 5.7999 nmole/L 442 Avg. Size (b.p.) 26.89 %CV

```
Sample Peak Width (sec): 50   Sample Min Peak Height: 25    Sample Baseline V to V?: Y    Sample Baseline V to V pts: 3
Sample Filter: Binomial       # of Pts for Filter: 3        Sample Start Region (min): 0  Sample End Region (min): 50
Manual Baseline Start (min): 10     Manual Baseline End (min): 48
Marker Peak Width (sec): 5    Marker Min Peak Height: 200   Marker Baseline V to V?: Y    Marker Baseline V to V pts: 3
Lower Marker Selection: First Peak > 200 RFU                Upper Marker Selection: Last Peak > 200 RFU
Ladder Size (bp): 1, 100, 200, 300, 400, 500, 600, 700, 800, 900, 1000, 1200, 1500, 2000, 3000, 6000
Quantification Using: Ladder          Final Concentration (ng/uL): 0.0830          Dilution Factor: 12.0
```


Copyright 2015 Advanced Analytical Technologies, Inc.

**Sample:** 103618-001-008
**Well Location:** H6
**Created:** Thursday, May 02, 2019 11:28:53 AM

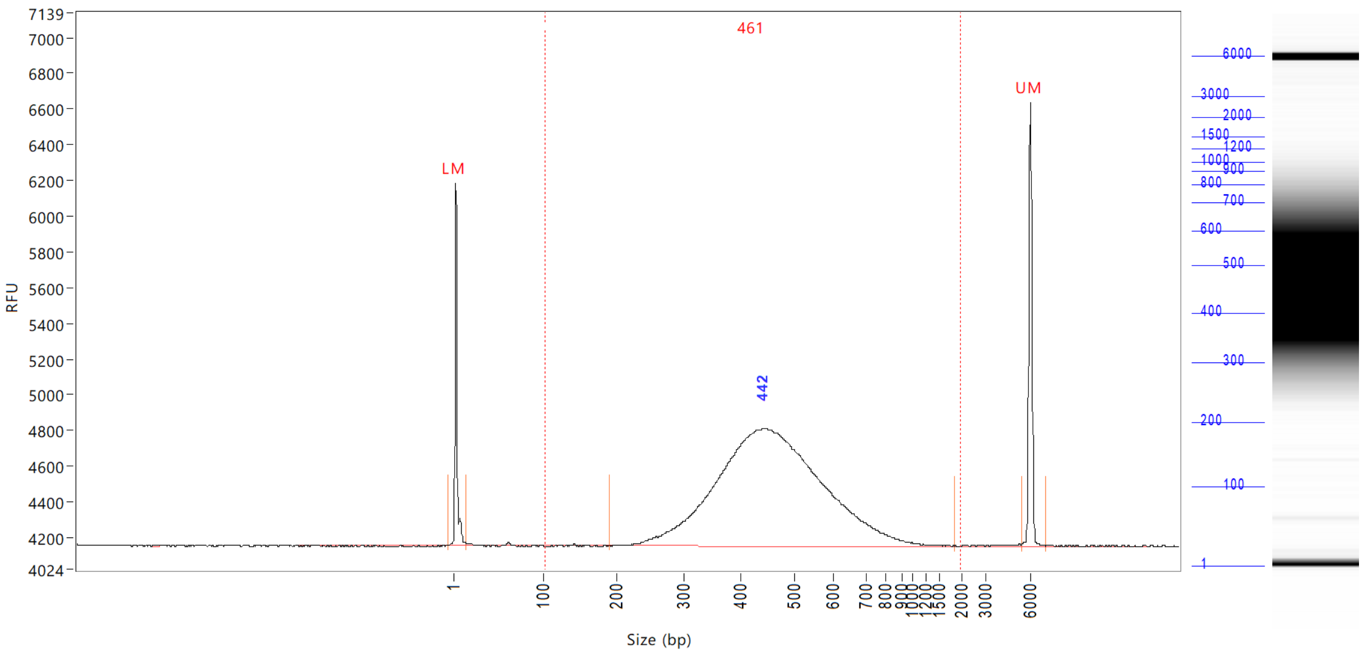


| Peak | Size (bp) | Conc. (ng/uL) | From (bp) | To (bp) | RFU |
|---|---|---|---|---|---|
| 1 | 1 (LM) | 0.0104 | 0 | 14 | 2027 |
| 2 | 442 | 1.1915 | 190 | 1859 | 659 |
| 3 | 6000 (UM) | 0.0083 | 5424 | 7003 | 2488 |

| | | |
|---|---|---|
| TIC: | 1.1915 | ng/uL |
| TIM: | 4.2529 | nmole/L |
| Total Conc.: | 1.1994 | ng/uL |

Smear Analysis 100 bp to 2000 bp 1.1925 ng/ul 99.4 %Total 4.2547 nmole/L 461 Avg. Size (b.p.) 25.16 %CV

Sample Peak Width (sec): 50 Sample Min Peak Height: 25 Sample Baseline V to V?: Y Sample Baseline V to V pts: 3
Sample Filter: Binomial # of Pts for Filter: 3 Sample Start Region (min): 0 Sample End Region (min): 50
Manual Baseline Start (min): 10 Manual Baseline End (min): 48
Marker Peak Width (sec): 5 Marker Min Peak Height: 200 Marker Baseline V to V?: Y Marker Baseline V to V pts: 3
Lower Marker Selection: First Peak > 200 RFU Upper Marker Selection: Last Peak > 200 RFU
Ladder Size (bp): 1, 100, 200, 300, 400, 500, 600, 700, 800, 900, 1000, 1200, 1500, 2000, 3000, 6000
Quantification Using: Ladder Final Concentration (ng/uL): 0.0830 Dilution Factor: 12.0

**Sample:** 103618-001-009
**Well Location:** A7
**Created:** Thursday, May 02, 2019 11:28:53 AM

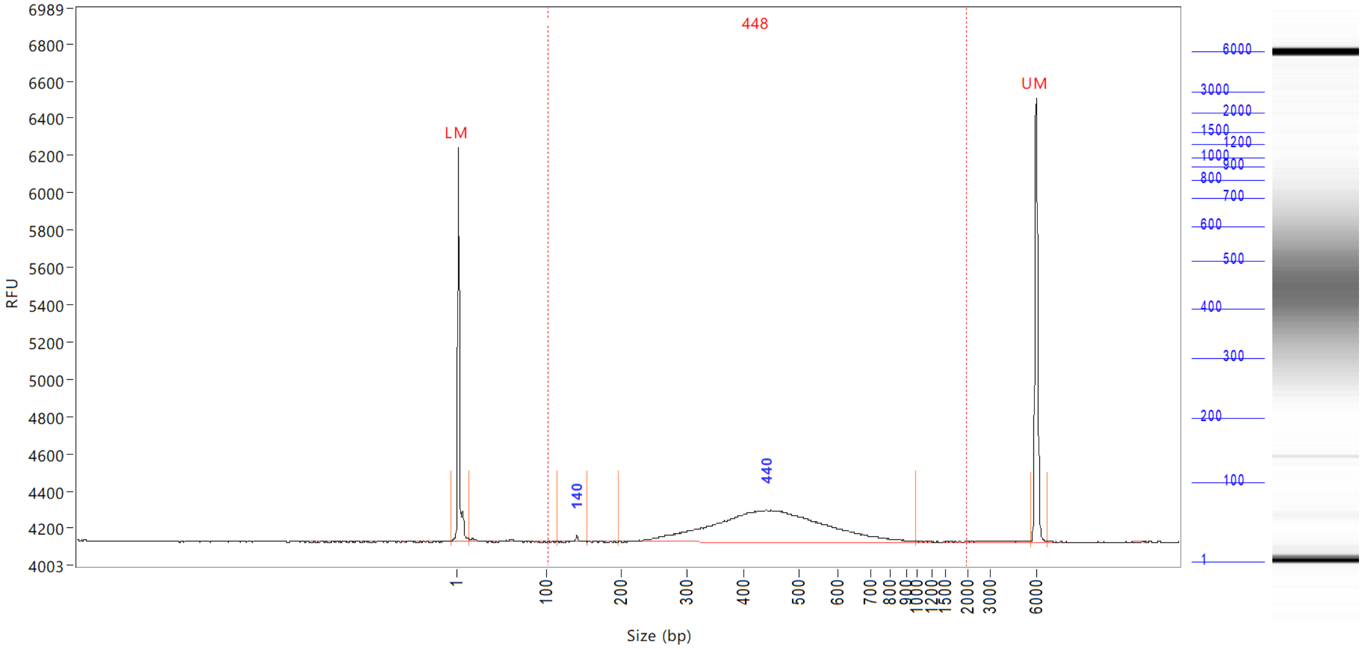


| Peak | Size (bp) | Conc. (ng/uL) | From (bp) | To (bp) | RFU |
|---|---|---|---|---|---|
| 1 | 1 (LM) | 0.0104 | 0 | 15 | 2113 |
| 2 | 140 | 0.0019 | 113 | 154 | 32 |
| 3 | 440 | 0.3119 | 196 | 996 | 170 |
| 4 | 6000 (UM) | 0.0074 | 5640 | 6764 | 2386 |

| | | |
|---|---|---|
| TIC: | 0.3138 | ng/uL |
| TIM: | 1.1706 | nmole/L |
| Total Conc.: | 0.3231 | ng/uL |

Smear Analysis 100 bp to 2000 bp 0.3150 ng/ul 97.5 %Total 1.1561 nmole/L 448 Avg. Size (b.p.) 29.13 %CV

Sample Peak Width (sec): 50 Sample Min Peak Height: 25 Sample Baseline V to V?: Y Sample Baseline V to V pts: 3
Sample Filter: Binomial # of Pts for Filter: 3 Sample Start Region (min): 0 Sample End Region (min): 50
Manual Baseline Start (min): 10 Manual Baseline End (min): 48
Marker Peak Width (sec): 5 Marker Min Peak Height: 200 Marker Baseline V to V?: Y Marker Baseline V to V pts: 3
Lower Marker Selection: First Peak > 200 RFU Upper Marker Selection: Last Peak > 200 RFU
Ladder Size (bp): 1, 100, 200, 300, 400, 500, 600, 700, 800, 900, 1000, 1200, 1500, 2000, 3000, 6000
Quantification Using: Ladder Final Concentration (ng/uL): 0.0830 Dilution Factor: 12.0

**Sample:** 103618-001-010
**Well Location:** B7
**Created:** Thursday, May 02, 2019 11:28:53 AM

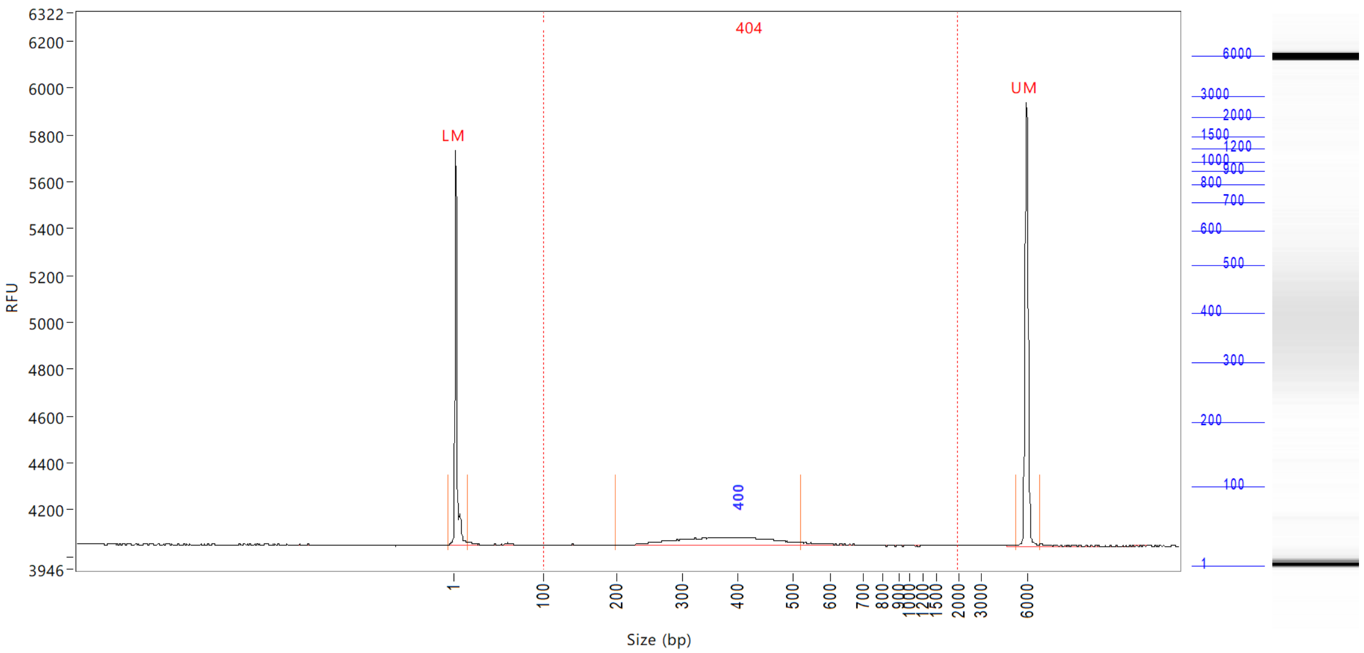


| Peak | Size (bp) | Conc. (ng/uL) | From (bp) | To (bp) | RFU |
|---|---|---|---|---|---|
| 1 | 1 (LM) | 0.0104 | 0 | 15 | 1685 |
| 2 | 400 | 0.0749 | 199 | 522 | 36 |
| 3 | 6000 (UM) | 0.0076 | 5328 | 6812 | 1897 |

| | | |
|---|---|---|
| TIC: | 0.0749 | ng/uL |
| TIM: | 0.3333 | nmole/L |
| Total Conc.: | 0.0914 | ng/uL |

Smear Analysis 100 bp to 2000 bp 0.0846 ng/ul 92.6 %Total 0.3448 nmole/L 404 Avg. Size (b.p.) 44.09 %CV

Sample Peak Width (sec): 50 Sample Min Peak Height: 25 Sample Baseline V to V?: Y Sample Baseline V to V pts: 3
Sample Filter: Binomial # of Pts for Filter: 3 Sample Start Region (min): 0 Sample End Region (min): 50
Manual Baseline Start (min): 10 Manual Baseline End (min): 48
Marker Peak Width (sec): 5 Marker Min Peak Height: 200 Marker Baseline V to V?: Y Marker Baseline V to V pts: 3
Lower Marker Selection: First Peak > 200 RFU Upper Marker Selection: Last Peak > 200 RFU
Ladder Size (bp): 1, 100, 200, 300, 400, 500, 600, 700, 800, 900, 1000, 1200, 1500, 2000, 3000, 6000
Quantification Using: Ladder Final Concentration (ng/uL): 0.0830 Dilution Factor: 12.0


Copyright 2015 Advanced Analytical Technologies, Inc.

**Sample:** 103618-001-011
**Well Location:** C7
**Created:** Thursday, May 02, 2019 11:28:53 AM

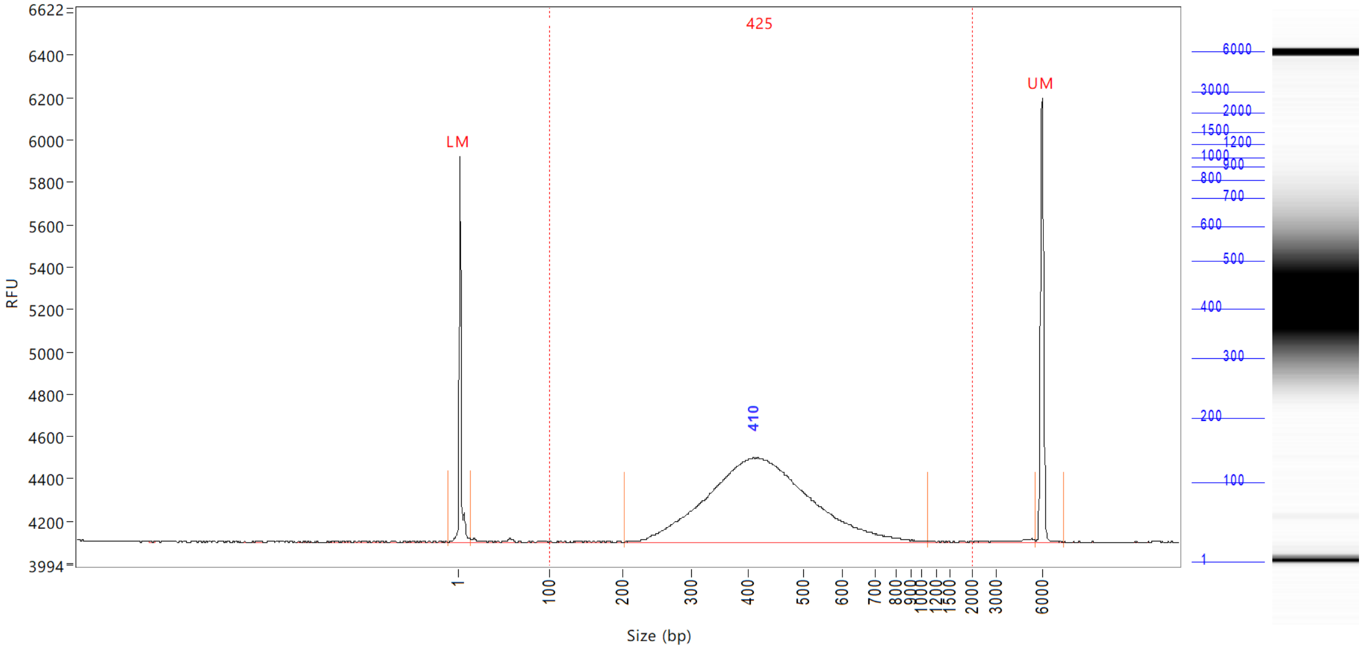


| Peak | Size (bp) | Conc. (ng/uL) | From (bp) | To (bp) | RFU |
|---|---|---|---|---|---|
| 1 | 1 (LM) | 0.0104 | 0 | 14 | 1819 |
| 2 | 410 | 0.7809 | 201 | 1088 | 398 |
| 3 | 6000 (UM) | 0.0076 | 5592 | 7432 | 2100 |

| | | |
|---|---|---|
| TIC: | 0.7809 | ng/uL |
| TIM: | 3.0304 | nmole/L |
| Total Conc.: | 0.7960 | ng/uL |

| | | | | | | |
|---|---|---|---|---|---|---|
| Smear Analysis | 100 bp to 2000 bp | 0.7845 ng/ul | 98.6 %Total | 3.0361 nmole/L | 425 Avg. Size (b.p.) | 25.95 %CV |

```
Sample Peak Width (sec): 50   Sample Min Peak Height: 25    Sample Baseline V to V?: Y    Sample Baseline V to V pts: 3
Sample Filter: Binomial       # of Pts for Filter: 3        Sample Start Region (min): 0  Sample End Region (min): 50
Manual Baseline Start (min): 10     Manual Baseline End (min): 48
Marker Peak Width (sec): 5    Marker Min Peak Height: 200   Marker Baseline V to V?: Y    Marker Baseline V to V pts: 3
Lower Marker Selection: First Peak > 200 RFU                Upper Marker Selection: Last Peak > 200 RFU
Ladder Size (bp): 1, 100, 200, 300, 400, 500, 600, 700, 800, 900, 1000, 1200, 1500, 2000, 3000, 6000
Quantification Using: Ladder          Final Concentration (ng/uL): 0.0830          Dilution Factor: 12.0
```


Copyright 2015 Advanced Analytical Technologies, Inc.

**Sample:** 103618-001-012
**Well Location:** D7
**Created:** Thursday, May 02, 2019 11:28:53 AM

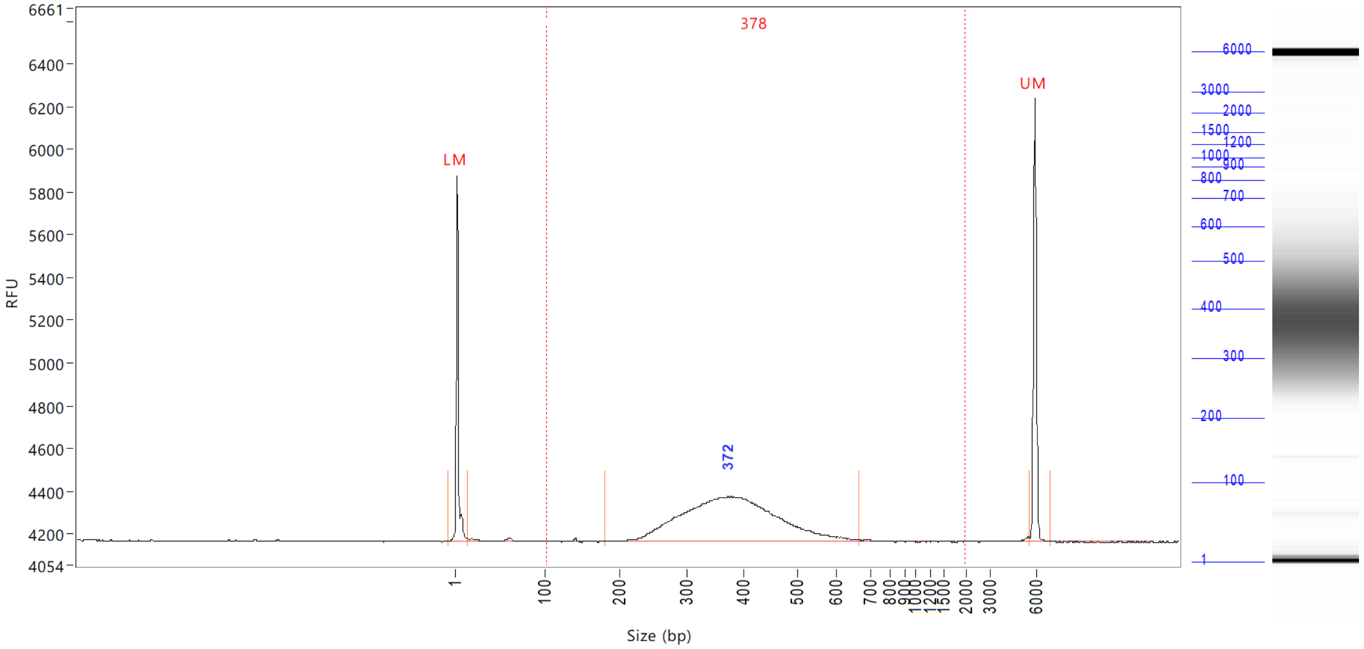


| Peak | Size (bp) | Conc. (ng/uL) | From (bp) | To (bp) | RFU |
|---|---|---|---|---|---|
| 1 | 1 (LM) | 0.0104 | 0 | 14 | 1709 |
| 2 | 372 | 0.4193 | 180 | 668 | 210 |
| 3 | 6000 (UM) | 0.0080 | 5616 | 7003 | 2081 |

| | | |
|---|---|---|
| TIC: | 0.4193 | ng/uL |
| TIM: | 1.8297 | nmole/L |
| Total Conc.: | 0.4290 | ng/uL |

Smear Analysis 100 bp to 2000 bp 0.4219 ng/ul 98.3 %Total 1.8341 nmole/L 378 Avg. Size (b.p.) 22.69 %CV

Sample Peak Width (sec): 50 Sample Min Peak Height: 25 Sample Baseline V to V?: Y Sample Baseline V to V pts: 3
Sample Filter: Binomial # of Pts for Filter: 3 Sample Start Region (min): 0 Sample End Region (min): 50
Manual Baseline Start (min): 10 Manual Baseline End (min): 48
Marker Peak Width (sec): 5 Marker Min Peak Height: 200 Marker Baseline V to V?: Y Marker Baseline V to V pts: 3
Lower Marker Selection: First Peak > 200 RFU Upper Marker Selection: Last Peak > 200 RFU
Ladder Size (bp): 1, 100, 200, 300, 400, 500, 600, 700, 800, 900, 1000, 1200, 1500, 2000, 3000, 6000
Quantification Using: Ladder Final Concentration (ng/uL): 0.0830 Dilution Factor: 12.0

 Copyright 2015 Advanced Analytical Technologies, Inc. 

**Sample:** 103618-001-013
**Well Location:** E7
**Created:** Thursday, May 02, 2019 11:28:53 AM

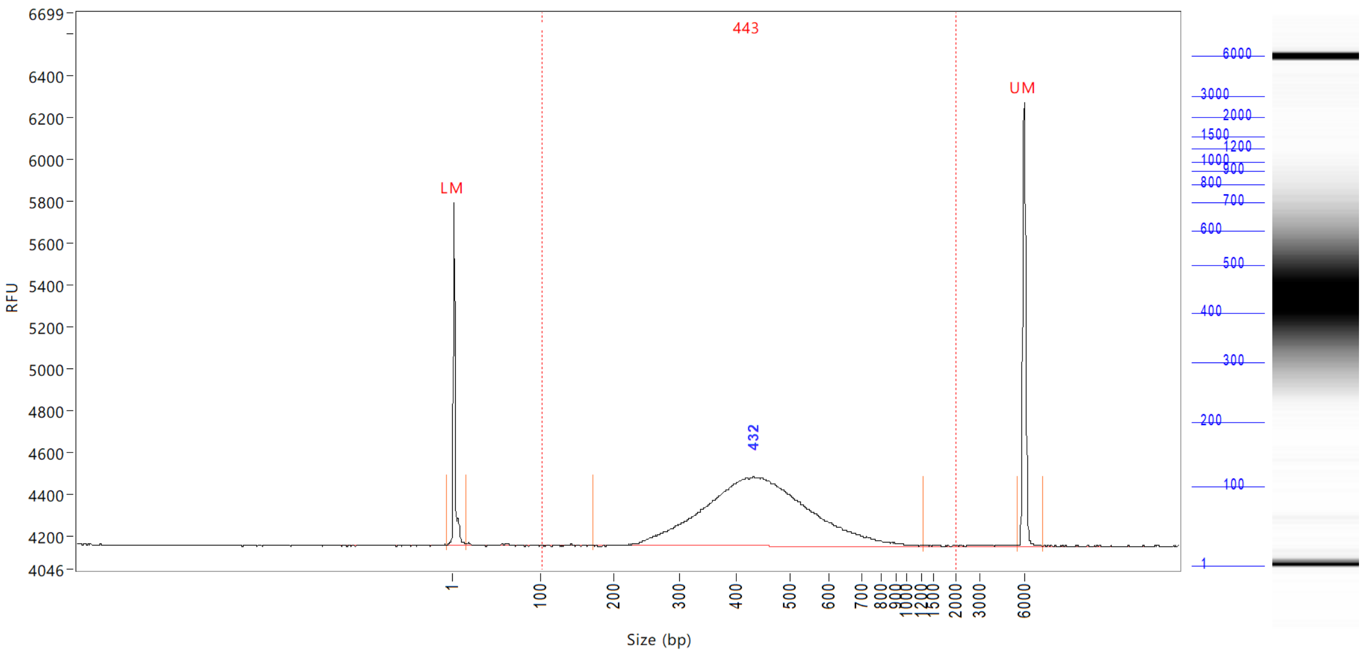


| Peak | Size (bp) | Conc. (ng/uL) | From (bp) | To (bp) | RFU |
|---|---|---|---|---|---|
| 1 | 1 (LM) | 0.0104 | 0 | 16 | 1638 |
| 2 | 432 | 0.7384 | 171 | 1254 | 329 |
| 3 | 6000 (UM) | 0.0086 | 5592 | 7241 | 2120 |

| | | |
|---|---|---|
| TIC: | 0.7384 | ng/uL |
| TIM: | 2.7488 | nmole/L |
| Total Conc.: | 0.7486 | ng/uL |

Smear Analysis | 100 bp to 2000 bp | 0.7406 ng/ul | 98.9 %Total | 2.7484 nmole/L | 443 Avg. Size (b.p.) | 26.79 %CV

Sample Peak Width (sec): 50 Sample Min Peak Height: 25 Sample Baseline V to V?: Y Sample Baseline V to V pts: 3
Sample Filter: Binomial # of Pts for Filter: 3 Sample Start Region (min): 0 Sample End Region (min): 50
Manual Baseline Start (min): 10 Manual Baseline End (min): 48
Marker Peak Width (sec): 5 Marker Min Peak Height: 200 Marker Baseline V to V?: Y Marker Baseline V to V pts: 3
Lower Marker Selection: First Peak > 200 RFU Upper Marker Selection: Last Peak > 200 RFU
Ladder Size (bp): 1, 100, 200, 300, 400, 500, 600, 700, 800, 900, 1000, 1200, 1500, 2000, 3000, 6000
Quantification Using: Ladder Final Concentration (ng/uL): 0.0830 Dilution Factor: 12.0


Copyright 2015 Advanced Analytical Technologies, Inc.

**Sample:** 103618-001-014
**Well Location:** F7
**Created:** Thursday, May 02, 2019 11:28:53 AM

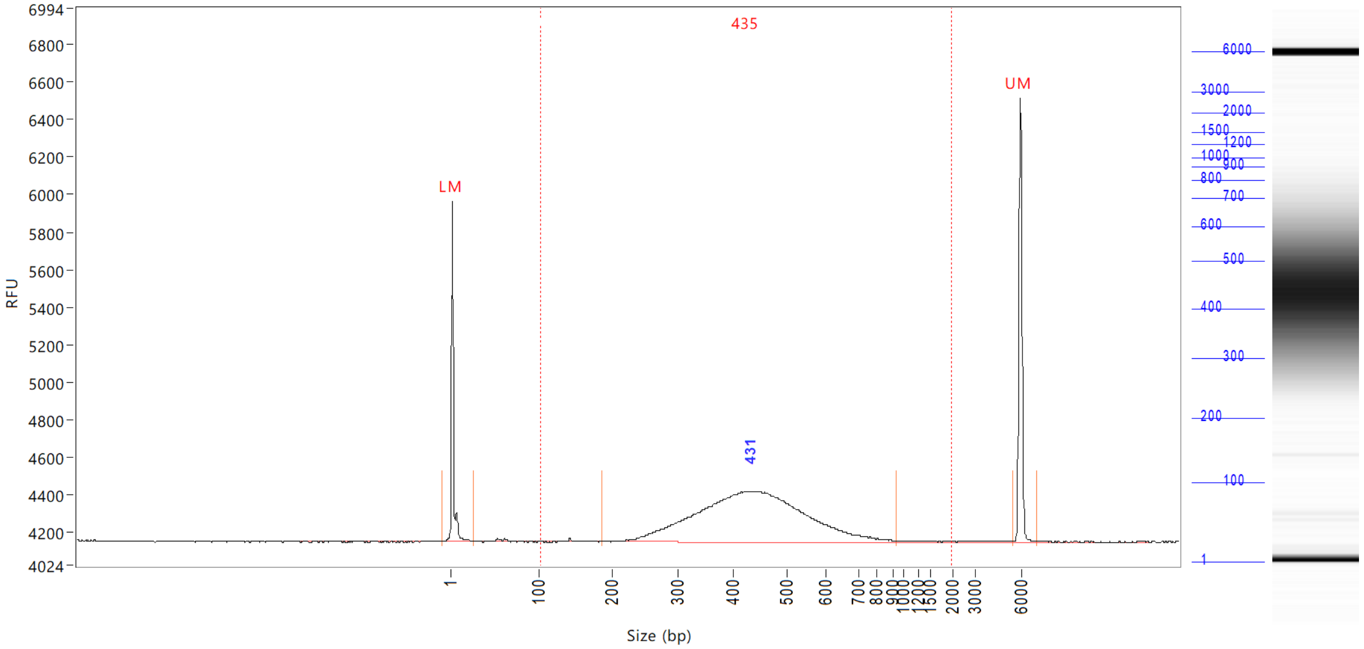


| Peak | Size (bp) | Conc. (ng/uL) | From (bp) | To (bp) | RFU |
|---|---|---|---|---|---|
| 1 | 1 (LM) | 0.0104 | 0 | 26 | 1816 |
| 2 | 431 | 0.5480 | 186 | 941 | 271 |
| 3 | 6000 (UM) | 0.0083 | 5496 | 7122 | 2371 |

| | | |
|---|---|---|
| TIC: | 0.5480 | ng/uL |
| TIM: | 2.0822 | nmole/L |
| Total Conc.: | 0.5629 | ng/uL |

Smear Analysis 100 bp to 2000 bp 0.5537 ng/ul 98.4 %Total 2.0965 nmole/L 435 Avg. Size (b.p.) 28.48 %CV

Sample Peak Width (sec): 50 Sample Min Peak Height: 25 Sample Baseline V to V?: Y Sample Baseline V to V pts: 3
Sample Filter: Binomial # of Pts for Filter: 3 Sample Start Region (min): 0 Sample End Region (min): 50
Manual Baseline Start (min): 10 Manual Baseline End (min): 48
Marker Peak Width (sec): 5 Marker Min Peak Height: 200 Marker Baseline V to V?: Y Marker Baseline V to V pts: 3
Lower Marker Selection: First Peak > 200 RFU Upper Marker Selection: Last Peak > 200 RFU
Ladder Size (bp): 1, 100, 200, 300, 400, 500, 600, 700, 800, 900, 1000, 1200, 1500, 2000, 3000, 6000
Quantification Using: Ladder Final Concentration (ng/uL): 0.0830 Dilution Factor: 12.0

 Copyright 2015 Advanced Analytical Technologies, Inc. 

**Sample:** 103618-001-015
**Well Location:** G7
**Created:** Thursday, May 02, 2019 11:28:53 AM

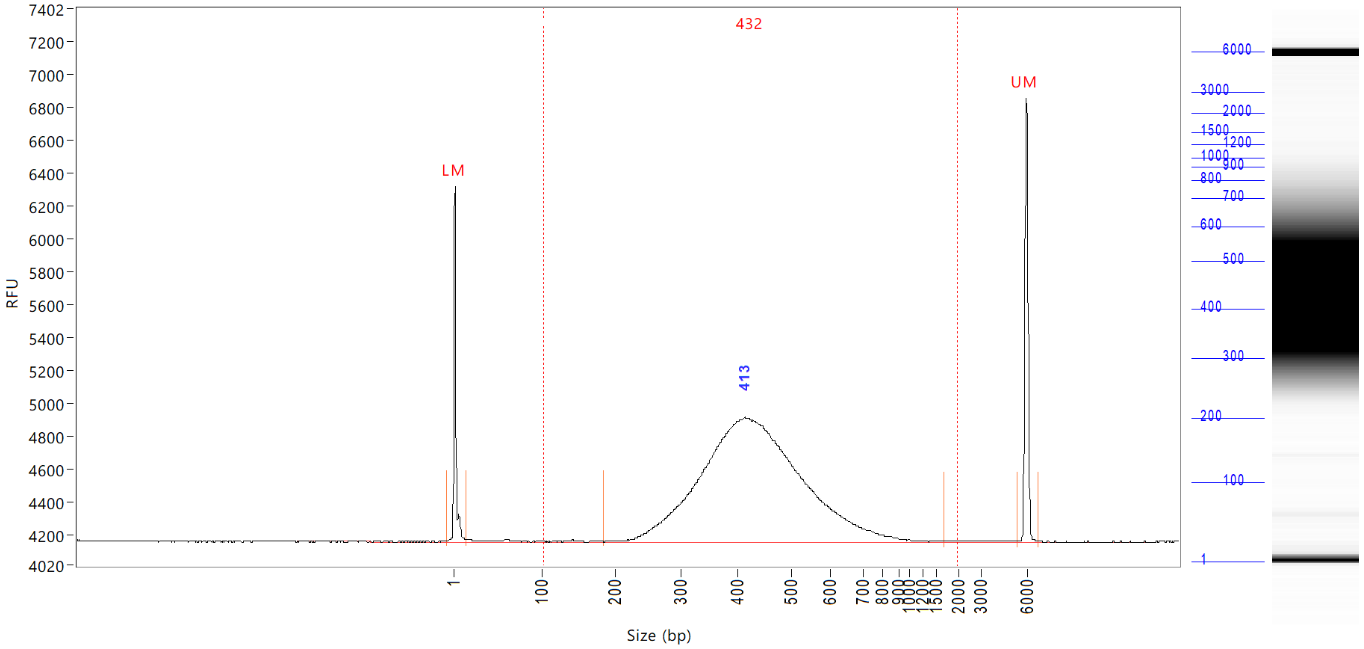


| Peak | Size (bp) | Conc. (ng/uL) | From (bp) | To (bp) | RFU |
|---|---|---|---|---|---|
| 1 | 1 (LM) | 0.0104 | 0 | 15 | 2158 |
| 2 | 413 | 1.2275 | 184 | 1692 | 757 |
| 3 | 6000 (UM) | 0.0084 | 5424 | 6788 | 2703 |

| | | |
|---|---|---|
| TIC: | 1.2275 | ng/uL |
| TIM: | 4.6750 | nmole/L |
| Total Conc.: | 1.2425 | ng/uL |

Smear Analysis 100 bp to 2000 bp 1.2308 ng/ul 99.1 %Total 4.6904 nmole/L 432 Avg. Size (b.p.) 26.27 %CV

Sample Peak Width (sec): 50 Sample Min Peak Height: 25 Sample Baseline V to V?: Y Sample Baseline V to V pts: 3
Sample Filter: Binomial # of Pts for Filter: 3 Sample Start Region (min): 0 Sample End Region (min): 50
Manual Baseline Start (min): 10 Manual Baseline End (min): 48
Marker Peak Width (sec): 5 Marker Min Peak Height: 200 Marker Baseline V to V?: Y Marker Baseline V to V pts: 3
Lower Marker Selection: First Peak > 200 RFU Upper Marker Selection: Last Peak > 200 RFU
Ladder Size (bp): 1, 100, 200, 300, 400, 500, 600, 700, 800, 900, 1000, 1200, 1500, 2000, 3000, 6000
Quantification Using: Ladder Final Concentration (ng/uL): 0.0830 Dilution Factor: 12.0


Copyright 2015 Advanced Analytical Technologies, Inc.

**Sample:** 103618-001-016
**Well Location:** H7
**Created:** Thursday, May 02, 2019 11:28:53 AM

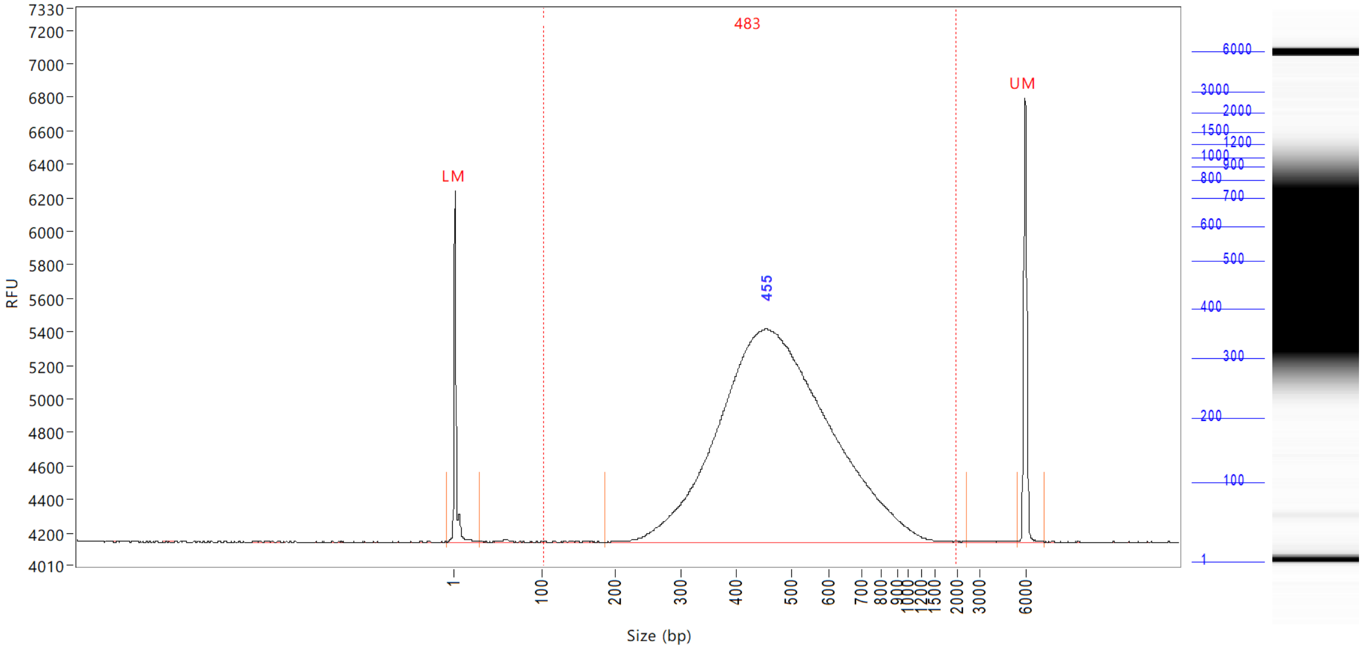


| Peak | Size (bp) | Conc. (ng/uL) | From (bp) | To (bp) | RFU |
|---|---|---|---|---|---|
| 1 | 1 (LM) | 0.0104 | 0 | 30 | 2092 |
| 2 | 455 | 2.1952 | 187 | 2413 | 1276 |
| 3 | 6000 (UM) | 0.0080 | 5448 | 7289 | 2653 |

| | | |
|---|---|---|
| TIC: | 2.1952 | ng/uL |
| TIM: | 7.4630 | nmole/L |
| Total Conc.: | 2.2074 | ng/uL |

Smear Analysis 100 bp to 2000 bp 2.1980 ng/ul 99.6 %Total 7.4807 nmole/L 483 Avg. Size (b.p.) 27.57 %CV

Sample Peak Width (sec): 50 Sample Min Peak Height: 25 Sample Baseline V to V?: Y Sample Baseline V to V pts: 3
Sample Filter: Binomial # of Pts for Filter: 3 Sample Start Region (min): 0 Sample End Region (min): 50
Manual Baseline Start (min): 10 Manual Baseline End (min): 48
Marker Peak Width (sec): 5 Marker Min Peak Height: 200 Marker Baseline V to V?: Y Marker Baseline V to V pts: 3
Lower Marker Selection: First Peak > 200 RFU Upper Marker Selection: Last Peak > 200 RFU
Ladder Size (bp): 1, 100, 200, 300, 400, 500, 600, 700, 800, 900, 1000, 1200, 1500, 2000, 3000, 6000
Quantification Using: Ladder Final Concentration (ng/uL): 0.0830 Dilution Factor: 12.0

**Sample:** 103618-001-017
**Well Location:** A8
**Created:** Thursday, May 02, 2019 11:28:53 AM

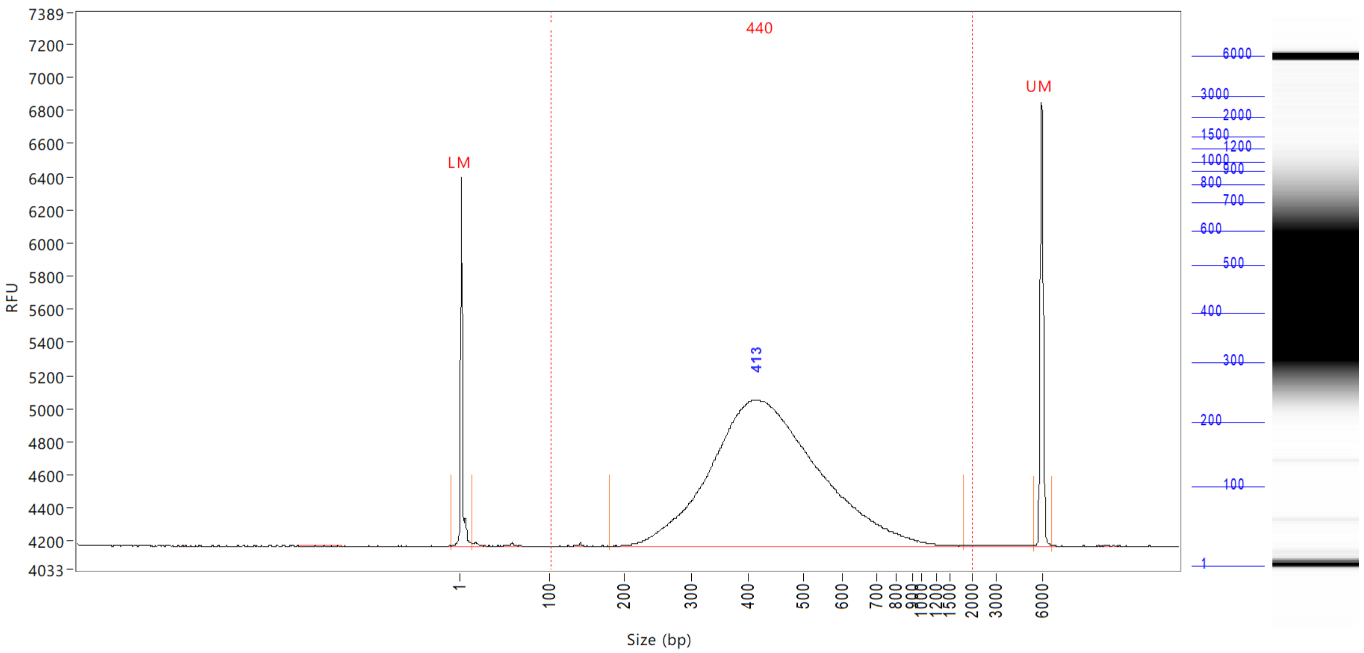


| Peak | Size (bp) | Conc. (ng/uL) | From (bp) | To (bp) | RFU |
|---|---|---|---|---|---|
| 1 | 1 (LM) | 0.0104 | 0 | 14 | 2225 |
| 2 | 413 | 1.4882 | 180 | 1834 | 882 |
| 3 | 6000 (UM) | 0.0078 | 5520 | 6644 | 2681 |

| | | |
|---|---|---|
| TIC: | 1.4882 | ng/uL |
| TIM: | 5.5653 | nmole/L |
| Total Conc.: | 1.4984 | ng/uL |

| | | | | | | |
|---|---|---|---|---|---|---|
| Smear Analysis | 100 bp to 2000 bp | 1.4899 ng/ul | 99.4 %Total | 5.5666 nmole/L | 440 Avg. Size (b.p.) | 27.30 %CV |

```
Sample Peak Width (sec): 50   Sample Min Peak Height: 25    Sample Baseline V to V?: Y    Sample Baseline V to V pts: 3
Sample Filter: Binomial       # of Pts for Filter: 3        Sample Start Region (min): 0  Sample End Region (min): 50
Manual Baseline Start (min): 10     Manual Baseline End (min): 48
Marker Peak Width (sec): 5    Marker Min Peak Height: 200   Marker Baseline V to V?: Y    Marker Baseline V to V pts: 3
Lower Marker Selection: First Peak > 200 RFU                Upper Marker Selection: Last Peak > 200 RFU
Ladder Size (bp): 1, 100, 200, 300, 400, 500, 600, 700, 800, 900, 1000, 1200, 1500, 2000, 3000, 6000
Quantification Using: Ladder          Final Concentration (ng/uL): 0.0830          Dilution Factor: 12.0
```


Copyright 2015 Advanced Analytical Technologies, Inc.

**Sample:** 103618-001-018
**Well Location:** B8
**Created:** Thursday, May 02, 2019 11:28:53 AM

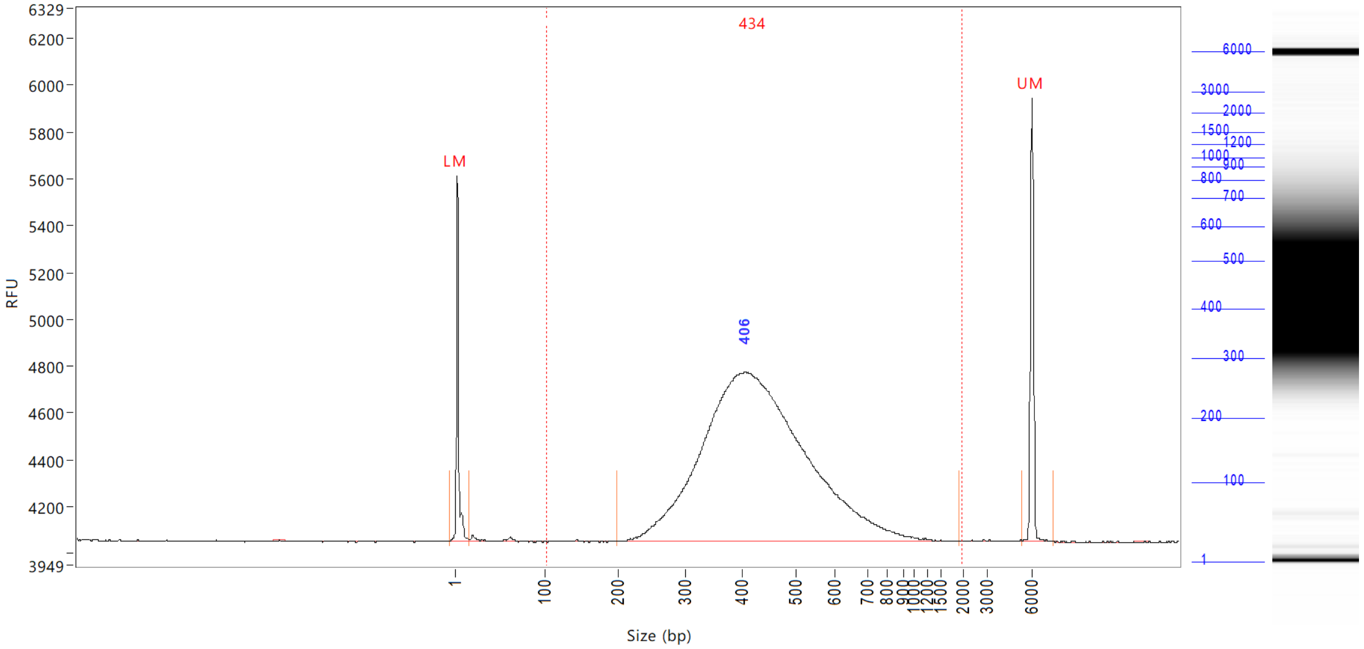


| Peak | Size (bp) | Conc. (ng/uL) | From (bp) | To (bp) | RFU |
|---|---|---|---|---|---|
| 1 | 1 (LM) | 0.0104 | 0 | 15 | 1560 |
| 2 | 406 | 1.6736 | 197 | 1942 | 725 |
| 3 | 6000 (UM) | 0.0081 | 5376 | 7361 | 1900 |

| | | |
|---|---|---|
| TIC: | 1.6736 | ng/uL |
| TIM: | 6.3595 | nmole/L |
| Total Conc.: | 1.6863 | ng/uL |

| | | | | | | |
|---|---|---|---|---|---|---|
| Smear Analysis | 100 bp to 2000 bp | 1.6744 ng/ul | 99.3 %Total | 6.3551 nmole/L | 434 Avg. Size (b.p.) | 26.92 %CV |

Sample Peak Width (sec): 50 Sample Min Peak Height: 25 Sample Baseline V to V?: Y Sample Baseline V to V pts: 3
Sample Filter: Binomial # of Pts for Filter: 3 Sample Start Region (min): 0 Sample End Region (min): 50
Manual Baseline Start (min): 10 Manual Baseline End (min): 48
Marker Peak Width (sec): 5 Marker Min Peak Height: 200 Marker Baseline V to V?: Y Marker Baseline V to V pts: 3
Lower Marker Selection: First Peak > 200 RFU Upper Marker Selection: Last Peak > 200 RFU
Ladder Size (bp): 1, 100, 200, 300, 400, 500, 600, 700, 800, 900, 1000, 1200, 1500, 2000, 3000, 6000
Quantification Using: Ladder Final Concentration (ng/uL): 0.0830 Dilution Factor: 12.0


Copyright 2015 Advanced Analytical Technologies, Inc.

**Sample:**
**Well Location:** H12:
**Created:** Thursday, May 02, 2019 11:28:53 AM
**Fit Type:** Point to Point

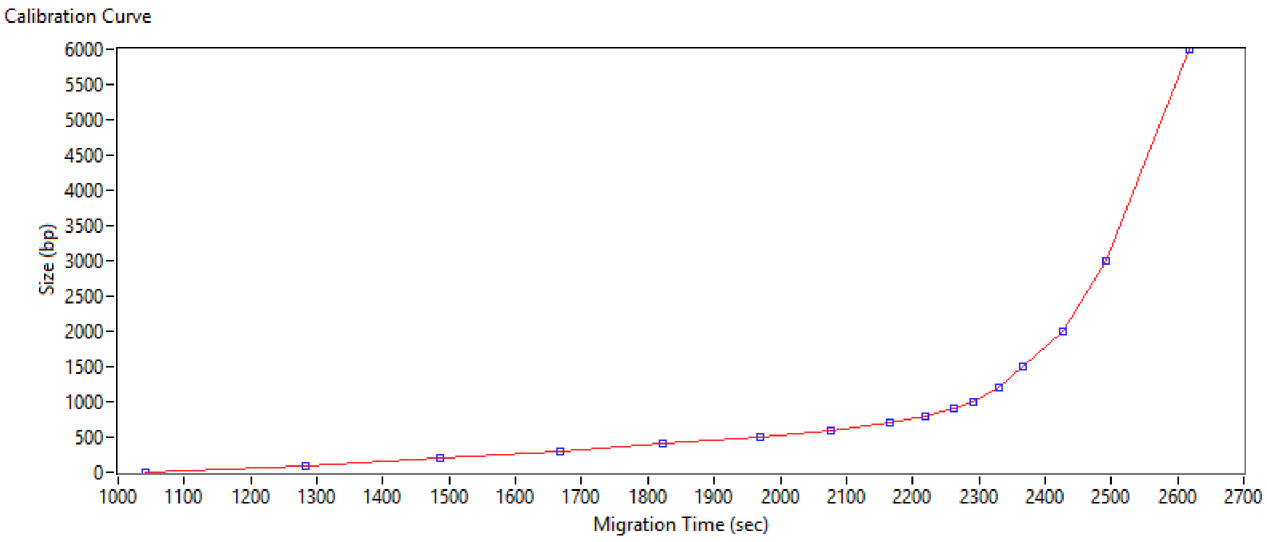


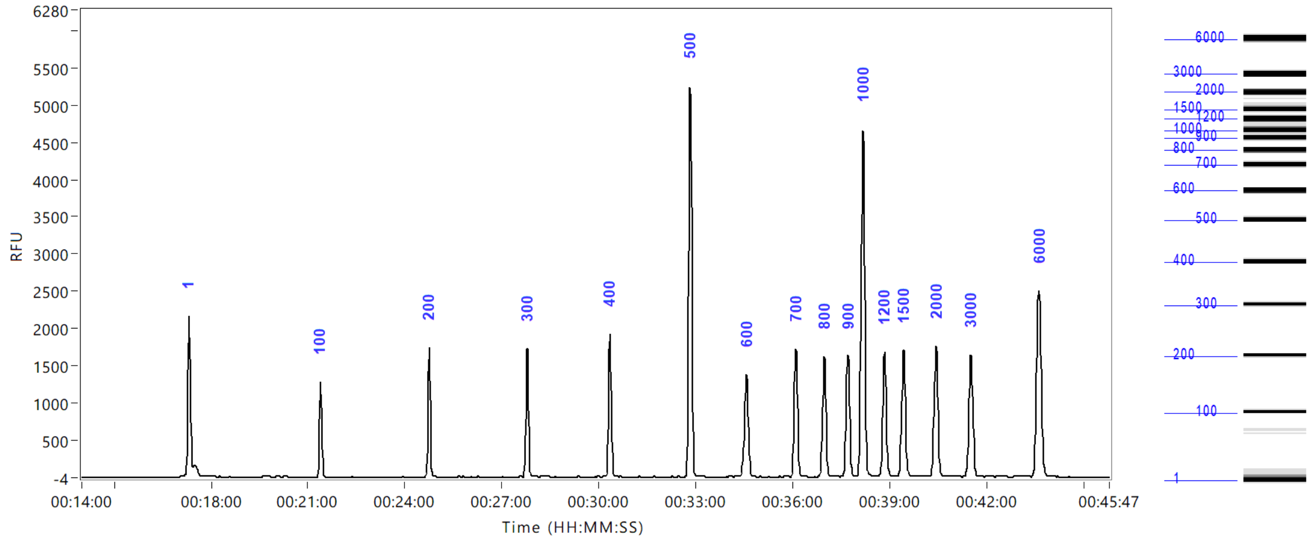



Copyright 2015 Advanced Analytical Technologies, Inc.
