## Supplementary material for "Rapid host response to an infection with Coronavirus. Study of transcriptional responses with Porcine Epidemic Diarrhea Virus": RNAseq data analysis rapport

### Data Analysis Report
#### RNA-Seq Analysis

**Customer Contact Information**

Mr M. Hulst
Wageningen Bioveterinary Research
Virology
Houtribweg 39
8221 RA Lelystad

**Project Information**

| | |
|---|---|
| Reference | AGS19055 / 103618 |
| Project start date | 2019-03-04 |
| Project manager | David van der Meer |
| Researcher(s) | Amrish Mahes (Bioinformatician) |
| Service type | Data analysis |
| Application | RNA-Seq analysis |
| Document status | Final v1 |

**Contact Information**

David van der Meer

☏ +31 71 568 1050

#### Summary

RNA-Seq analysis using our RNA-Seq and differential expression analysis pipeline was performed on a short read data set obtained using Illumina next generation sequencing technology. The pipeline has several steps, statistics, and intermediate products that will be reviewed in this report.

#### Introduction

This document describes the general workflow of the data analysis. Please refer to the Guidelines (provided on the data disk) for further details.

#### Materials

A data set of 17 sample(s) generated using an Illumina NovaSeq 6000 was used for RNA-Seq data analysis. The samples ID linkage between the GenomeScan IDs, customer IDs, sample groups, and raw input files are shown in Table 1.

**Table 1.** Sample information

| Sample ID | Customer ID | Sample Group | # Data files |
|---|---|---|---|
| 103618-001-001 | 0-PEDV | A1 | 4 |
| 103618-001-002 | 0-PEDV control | B1 | 4 |
| 103618-001-003 | 0-MRV | C1 | 4 |
| 103618-001-004 | 0-MRV control | D1 | 4 |
| 103618-001-005 | 0-PEDV-MRV | E1 | 2 |
| 103618-001-006 | 0-PEDV-MRV control | F1 | 4 |
| 103618-001-007 | 4-PEDV | G1 | 2 |
| 103618-001-008 | 4-PEDV control | H1 | 4 |
| 103618-001-009 | 4-MRV | A2 | 4 |
| 103618-001-011 | 4-PEDV-MRV | C2 | 4 |
| 103618-001-012 | 4-PEDV-MRV control | D2 | 4 |
| 103618-001-013 | 6-PEDV | E2 | 4 |
| 103618-001-014 | 6-PEDV control | F2 | 4 |
| 103618-001-015 | 6-MRV | G2 | 4 |
| 103618-001-016 | 6-MRV control | H2 | 4 |
| 103618-001-017 | 6-PEDV-MRV | A3 | 4 |
| 103618-001-018 | 6-PEDV-MRV control | B3 | 4 |

The raw FASTQ data files corresponding with the file names in the sample ID table (Table 1) can be found in the Raw Data directory on the data disk.

#### Experimental procedures

GenomeScan delivers the alignment (BAM with index), feature counts, and differential expression analysis using the RNA-Seq v4 pipeline that was developed in-house and described in standard operating procedure SOP 70 (RNA-Seq Analysis) (Fig. 1). The workflow includes raw data quality control (INS 92), adapter trimming (INS 88), alignment of short reads (INS 61), feature counting and differential expression (INS 79).

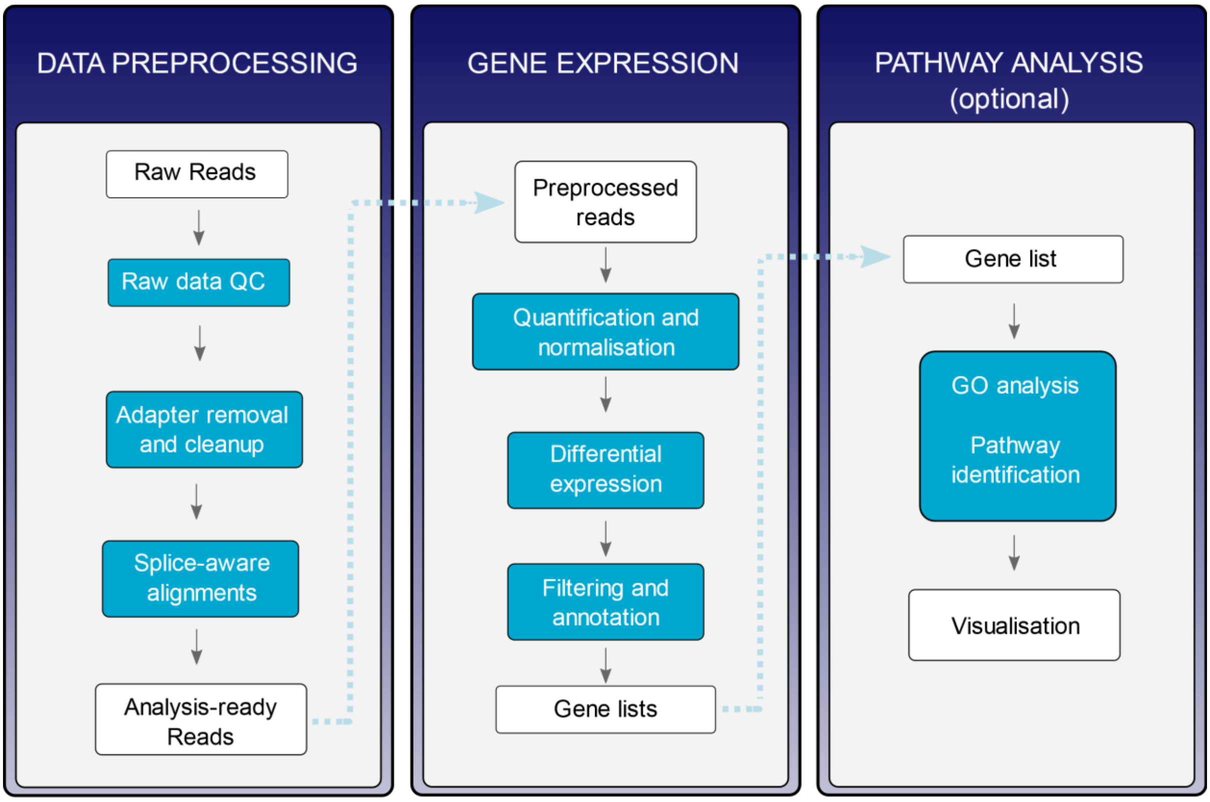


**Fig. 1.** Data analysis workflow.

#### Results

##### Raw data quality

The pipeline started with a quality control stage. In this stage checks for possible sample and barcode contamination were performed and a set of standard quality metrics for the raw data set was determined using our in-house QC v1 tool. The results are shown in Table 2.

**Table 2.** Raw data quality

| Sample ID | Read 1 (forward) | | | | Read 2 (reverse) | | | |
|---|---|---|---|---|---|---|---|---|
| | Reads | Length | Avg Q | Bases | Reads | Length | Avg Q | Bases |
| 103618-001-001 | 21465636 | 151 | 36.1 | 3241311036 | 21465636 | 151 | 35.7 | 3241311036 |
| 103618-001-002 | 19006413 | 151 | 36.1 | 2869968363 | 19006413 | 151 | 35.7 | 2869968363 |
| 103618-001-003 | 30874087 | 151 | 36.1 | 4661987137 | 30874087 | 151 | 35.7 | 4661987137 |
| 103618-001-004 | 16787245 | 151 | 36.1 | 2534873995 | 16787245 | 151 | 35.8 | 2534873995 |
| 103618-001-005 | 21844585 | 151 | 36.3 | 3298532335 | 21844585 | 151 | 36.0 | 3298532335 |
| 103618-001-006 | 21939961 | 151 | 36.2 | 3312934111 | 21939961 | 151 | 35.9 | 3312934111 |
| 103618-001-007 | 18341732 | 151 | 36.2 | 2769601532 | 18341732 | 151 | 35.8 | 2769601532 |
| 103618-001-008 | 21373119 | 151 | 36.2 | 3227340969 | 21373119 | 151 | 35.5 | 3227340969 |
| 103618-001-009 | 18632923 | 151 | 36.2 | 2813571373 | 18632923 | 151 | 35.6 | 2813571373 |
| 103618-001-011 | 19275279 | 151 | 36.3 | 2910567129 | 19275279 | 151 | 35.8 | 2910567129 |
| 103618-001-012 | 23147037 | 151 | 36.2 | 3495202587 | 23147037 | 151 | 35.9 | 3495202587 |
| 103618-001-013 | 21673452 | 151 | 36.1 | 3272691252 | 21673452 | 151 | 35.7 | 3272691252 |
| 103618-001-014 | 16185666 | 151 | 36.2 | 2444035566 | 16185666 | 151 | 35.8 | 2444035566 |
| 103618-001-015 | 18985012 | 151 | 36.1 | 2866736812 | 18985012 | 151 | 35.7 | 2866736812 |

| | | | | | | | | |
|---|---|---|---|---|---|---|---|---|
| 103618-001-016 | 18734320 | 151 | 36.2 | 2828882320 | 18734320 | 151 | 35.2 | 2828882320 |
| 103618-001-017 | 16950382 | 151 | 36.2 | 2559507682 | 16950382 | 151 | 35.7 | 2559507682 |
| 103618-001-018 | 18889320 | 151 | 36.1 | 2852287320 | 18889320 | 151 | 35.7 | 2852287320 |

#### Adapter clipping

Prior to alignment, the reads were trimmed for adapter sequences using Trimmomatic v0.30. Presumed adapter sequences were removed from the read when the bases matched a sequence in the adapter sequence set (TruSeq adapters) with 2 or less mismatches and an alignment score of at least 12. The number of the reads and bases before and after trimming are shown in Table 3.

**Table 3.** Adapter clipping

| Sample ID | Read pairs | | Read 1 (forward) | | | Read 2 (reverse) | | |
|---|---|---|---|---|---|---|---|---|
| | Kept | % kept | Length | Bases | % kept | Length | Bases | % kept |
| 103618-001-001 | 21344808 | 99.4 | 149.0 | 3196269379 | 98.6 | 150.9 | 3222256881 | 99.4 |
| 103618-001-002 | 18927958 | 99.6 | 149.1 | 2833841716 | 98.7 | 151.0 | 2857531956 | 99.6 |
| 103618-001-003 | 30591562 | 99.1 | 148.4 | 4563860037 | 97.9 | 151.0 | 4618453858 | 99.1 |
| 103618-001-004 | 16722681 | 99.6 | 149.1 | 2505144001 | 98.8 | 151.0 | 2524719195 | 99.6 |
| 103618-001-005 | 21805325 | 99.8 | 149.8 | 3266347306 | 99.0 | 151.0 | 3292359380 | 99.8 |
| 103618-001-006 | 21898461 | 99.8 | 149.2 | 3281985194 | 99.1 | 150.9 | 3305462873 | 99.8 |
| 103618-001-007 | 18324761 | 99.9 | 149.9 | 2747762725 | 99.2 | 151.0 | 2766419095 | 99.9 |
| 103618-001-008 | 21331419 | 99.8 | 149.2 | 3198609187 | 99.1 | 151.0 | 3220887552 | 99.8 |
| 103618-001-009 | 18350672 | 98.5 | 148.2 | 2736776825 | 97.3 | 150.9 | 2770110633 | 98.5 |
| 103618-001-011 | 19250245 | 99.9 | 149.1 | 2880642264 | 99.0 | 151.0 | 2906337280 | 99.9 |
| 103618-001-012 | 23005431 | 99.4 | 146.7 | 3399202064 | 97.3 | 150.8 | 3471407916 | 99.3 |
| 103618-001-013 | 21629853 | 99.8 | 148.7 | 3233108375 | 98.8 | 151.0 | 3265673516 | 99.8 |
| 103618-001-014 | 16075520 | 99.3 | 147.6 | 2392094470 | 97.9 | 151.0 | 2426948081 | 99.3 |
| 103618-001-015 | 18955105 | 99.8 | 149.1 | 2836823198 | 99.0 | 150.9 | 2861328729 | 99.8 |
| 103618-001-016 | 18726292 | 100.0 | 150.2 | 2816148654 | 99.5 | 151.0 | 2827152387 | 99.9 |
| 103618-001-017 | 16907918 | 99.7 | 149.4 | 2531076732 | 98.9 | 150.9 | 2551594999 | 99.7 |
| 103618-001-018 | 18865675 | 99.9 | 148.8 | 2820553698 | 98.9 | 150.9 | 2847478055 | 99.8 |

#### Quality filtering

For this application and type of data quality filtering is not required to obtain proper count data from this data set.

#### Mapping of reads to the reference

The GCF_000409795.2_Chlorocebus_sabeus_1.1_genomic reference was used for alignment of the reads for each sample. The reference files are provided in the References directory. The reads were mapped to the reference sequence using a short read aligner based on Burrows—Wheeler Transform (Tophat v2.0.14) with default settings. The alignment files (BAM files sorted on coordinates and indexed with the samtools v1.3 package) containing the mapping information are provided in the Mapping folder on the hard disk (Table 4).

**Table 4.** Mapping of reads

| Sample ID | Filtered reads | Mapped reads | % Mapping | Multiple mapped | % Multiple |
|---|---|---|---|---|---|
| 103618-001-001 | 21344808 | 18372494 | 86.1 | 1115911 | 5.2 |
| 103618-001-002 | 18927958 | 16350900 | 86.4 | 916827 | 4.8 |
| 103618-001-003 | 30591562 | 26177817 | 85.6 | 1570924 | 5.1 |

| | | | | | |
|---|---|---|---|---|---|
| 103618-001-004 | 16722681 | 14469826 | 86.5 | 792785 | 4.7 |
| 103618-001-005 | 21805325 | 19085203 | 87.5 | 1048210 | 4.8 |
| 103618-001-006 | 21898461 | 19014562 | 86.8 | 1034550 | 4.7 |
| 103618-001-007 | 18324761 | 15937134 | 87.0 | 870073 | 4.7 |
| 103618-001-008 | 21331419 | 18642835 | 87.4 | 1067486 | 5.0 |
| 103618-001-009 | 18350672 | 15595904 | 85.0 | 1046330 | 5.7 |
| 103618-001-011 | 19250245 | 16671586 | 86.6 | 918282 | 4.8 |
| 103618-001-012 | 23005431 | 19119313 | 83.1 | 1116125 | 4.9 |
| 103618-001-013 | 21629853 | 18509839 | 85.6 | 1050915 | 4.9 |
| 103618-001-014 | 16075520 | 13685292 | 85.1 | 824550 | 5.1 |
| 103618-001-015 | 18955105 | 16203034 | 85.5 | 892565 | 4.7 |
| 103618-001-016 | 18726292 | 16470960 | 88.0 | 919521 | 4.9 |
| 103618-001-017 | 16907918 | 14588464 | 86.3 | 820667 | 4.9 |
| 103618-001-018 | 18865675 | 16218358 | 86.0 | 914311 | 4.8 |

#### Feature counting

Based on the mapped locations in the alignment file the frequency of how often a read was mapped on a transcript was determined with HTSeq v0.11.0. The counts were saved to count files, which serve as input for downstream RNA-Seq differential expression analysis. The count files are provided in the folder Counts on the hard disk.

Additionally, RPKM/FPKM (reads/fragments per kilobase of exon per million reads mapped) values were calculated and provided in the 'Fpkm' directory.

#### Differential expression

The read counts were loaded into the DESeq2 package v1.14.1, a statistical package within the R platform v3.3.0. DESeq2 was specifically developed to find differentially expressed genes between two conditions for RNA-Seq data with small sample size and over-dispersion and uses an expression curve model based on a negative binomial distribution and local regression to estimate the relationship between the mean and variance of each gene. Furthermore, it allows scaling factors to be easily included in the statistical test. Since DESeq2 can only handle groups or conditions with replicates, single-sample comparisons are performed with DESeq v1.30.0. The differential expression comparison grouping is provided in Table 5. For this analyses DESeq v1.30 was used.

**Table 5.** Expression comparison setup and differentially expressed genes

| Comparison | Condition A | Condition B | DE features |
|---|---|---|---|
| A1 vs B1 | 0-PEDV | 0-PEDV control | 1 |
| A3 vs B3 | 6-PEDV-MRV | 6-PEDV-MRV control | 127 |
| C1 vs D1 | 0-MRV | 0-MRV control | 2 |
| C2 vs D2 | 4-PEDV-MRV | 4-PEDV-MRV control | 95 |
| E1 vs F1 | 0-PEDV-MRV | 0-PEDV-MRV control | 7 |
| E2 vs F2 | 6-PEDV | 6-PEDV control | 1 |
| G1 vs H1 | 4-PEDV | 4-PEDV control | 2 |
| G2 vs H2 | 6-MRV | 6-MRV control | 33 |

#### Clustering and normalisation

R generates a sample heatmap based on the relative distances calculated by DESeq (Fig. 2).

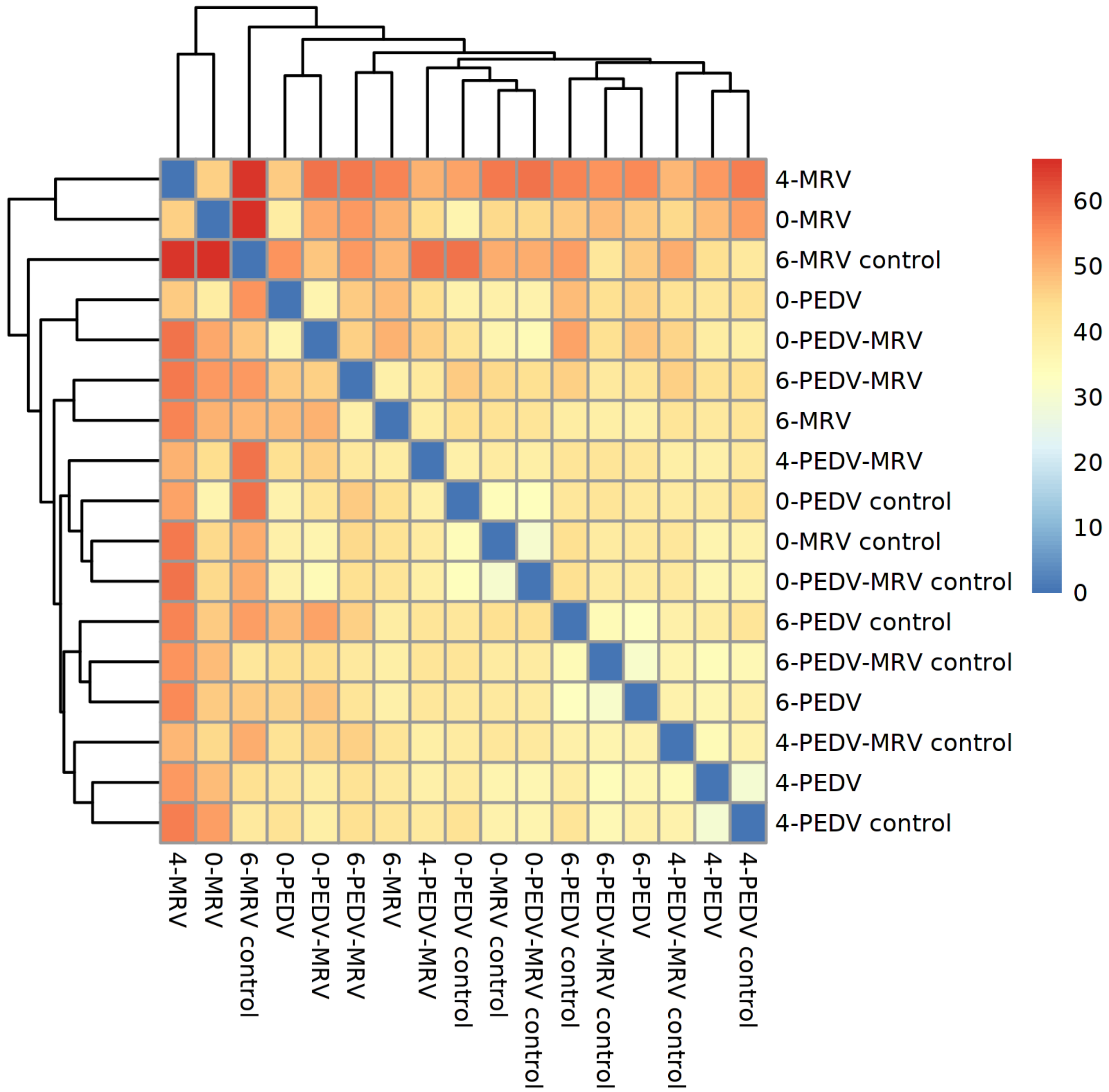


**Fig. 2**. The heatmap for all samples, based on relative distances of the samples. Blue means identical, red means very different.

To determine if the normalisation of the gene expression data successfully suppressed most deviations, MA plots were generated (Fig. 3). A MA plot can identify systematic deviations in the data, under the assumption that for most genes their expression is not significantly different, when compared between two sample groups.

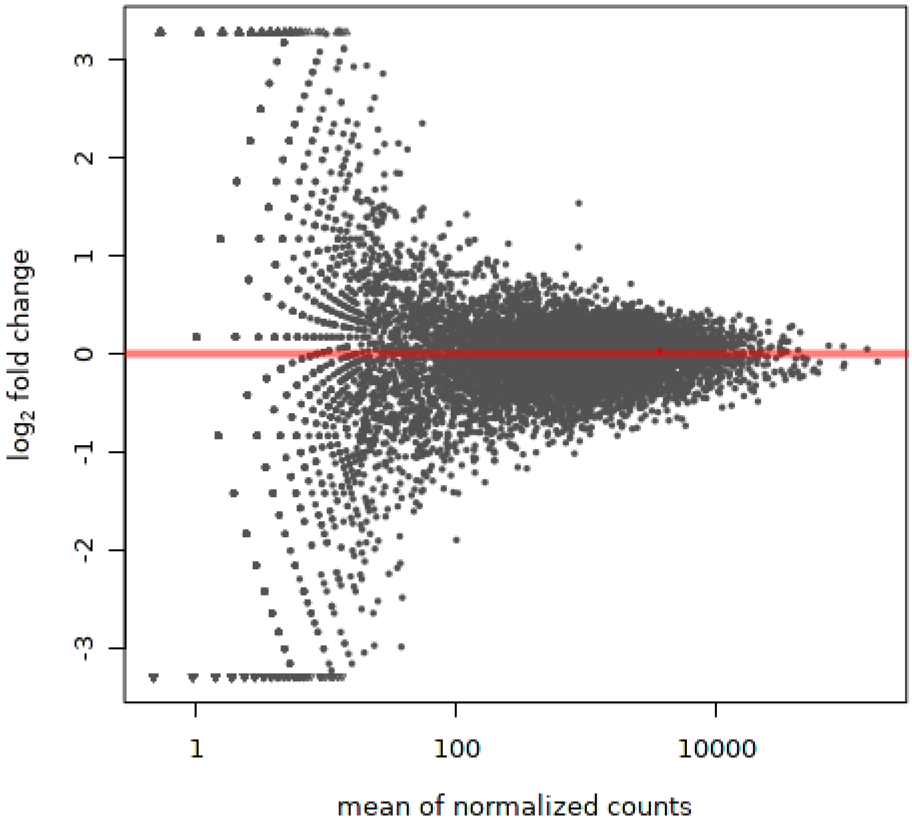


**Fig. 3.** MA plot for group A1 vs group B1.
All plots are included in the folder Plots on the hard disk.

##### Differentially expressed genes

The main output result of the differential expression analysis are the lists of differentially expressed genes (DE genes). The analysis consisted of 8 comparisons. All groups were compared according to the customer's specifications as stated in Table 5. This resulted in the amount of DE genes per comparison as shown in Table 5.

The lists with annotated differentially expressed genes are included in the folder 'Differential' on the hard disk.

#### Conclusions

Our standard procedure filtered the raw Illumina sequencing data, aligned the reads of all samples to the reference with annotation, and counted reads mapped to gene annotated regions per sample based on this mapping. Based on the counts, differential expression analysis was performed. The pipeline was executed successfully. All relevant data and overview files have been included on the data hard disk.

#### Computing hardware

The analysis was performed at the following site(s): (1) the GenomeScan SGE cluster located at GenomeScan, Plesmanlaan 1d, 2333 BZ, Leiden, The Netherlands.

#### Deviations, additions, exclusions from the test method

This project was performed in compliance with the requirements and specifications set by GenomeScan and the customer. All data are within specifications. There were no deviations from the standard test procedures.

#### Data transfer and file formats

The data files are stored on the accompanying data disk as part of the results delivery. Within the data analysis directory, the following directories and files are present:

| | |
|---|---|
| Mapping: | Alignment files in .bam format and index files in .bai format. |
| References: | Reference sequence(s) and annotation, usually provided by the customer |
| Customer: | Optional directory with data provided by the customer. |
| Plots: | Plot image files. These files include:<br>o Heatmaps<br>o MA plots<br>All plots generated per comparison where applicable. |
| Counts: | Raw count files. |
| Differential: | Lists with differentially expressed genes per comparison. |
| Fpkm: | FPKM tables. |
| Guidelines: | Guidelines for NGS services in PDF format |
| Reports: | Data analysis report in PDF format |
| Export: | Optional spreadsheet/tab-delimited versions of the variant list, filtered lists, or derived data. |

All files provided are in either plain text (.txt), tab-delimited text (.tsv, .txt, .count, .fpkm_tracking), Variant Call format (.vcf), PDF format (.pdf), PNG format (.png), or OpenDocument format (.odt, .ods). Plain text files can be opened in any text editor although some files for the Next Generation Sequencing service may be too large and an editor optimised for large files is required. Tab-delimited files can be opened in any text editor or spreadsheet application. PDF files require Adobe Reader to open and OpenDocument files can be opened in Microsoft Office 2007 SP2 or later versions.

Tab delimited files (.tsv) are formatted using a dot as decimal separator and without thousand separators. They should be opened using an English locale to avoid incorrect translation of decimal and thousand separators.

This document is digitally signed using certificates issued by the certificate authority CAcert. To validate the signatures you may need to add the CAcert root certificate (www.cacert.org) to the key store of your viewer or browser (one-time only). Alternatively, you may validate signatures via an online signature validation tool.

Signed,

Digitally signed by: Amrish Mahes
Bioinformatics Analyst
Reason: Approval
Date: 2019-06-12 15:55:21
Location: GenomeScan B.V., Leiden

Digitally signed by: Thomas Chin-A-Woeng
Project Manager Bioinformatics
Reason: Approval
Date: 2019-06-12 15:54:48
Location: GenomeScan B.V., Leiden

Date: 2019-06-12
